# Pathogen Subversion of Neuro-Epidermal Signaling Impairs Lysosomal Function to Disrupt Collagen Homeostasis

**DOI:** 10.1101/2025.06.28.662087

**Authors:** Qian Li, Yating Liu, Hanyi Chen, Weilie Xiao, Bin Qi

## Abstract

The epidermis relies on collagen-rich extracellular matrices (ECMs) to maintain barrier integrity against pathogens. Lysosomes regulate cuticle collagen turnover, yet how neuronal signaling modulates epidermal lysosomal function and collagen organization during infection remains unclear. Using *Pseudomonas aeruginosa* PA14-*Caenorhabditis elegans* infection model, we demonstrate that pathogen-induced neuronal signaling disrupts epidermal lysosomal activity and collagen remodeling. PA14 infection triggers neurons to secrete NSIF-1 (Neuronal Secreted Immune Factor 1), which translocates to the epidermis and impairs lysosomal acidification, maturation, and degradation by suppressing the transcription factor ELT-3. This disruption leads to disorganized collagen structure, compromising cuticle integrity and host resistance. Genetic mutation of *nsif-1* restores lysosomal function, enhances collagen density, and improves survival, while neuron-specific *nsif-1* knockdown confirms its neuronal origin. Moreover, NSIF-1 inhibits ELT-3 nuclear localization, blocking its role in lysosomal-dependent ECM repair. Our study reveals a neuro-epidermal axis wherein pathogens exploit neuronal signals to disrupt lysosomal function and collagen homeostasis, identifying NSIF-1 and ELT-3 as potential targets to counteract infection-driven ECM dysregulation.

## Introduction

The epidermis serves as the primary barrier against pathogens, relying on structural components such as collagen to maintain its integrity and immune function. In mammals, the extracellular matrix (ECM) of the skin is rich in collagens, which not only provide mechanical strength but also act as scaffolds for immune signaling pathways. Many pathogens, including bacteria and viruses, exploit ECM components and associated molecules to adhere to host tissues, degrade structural barriers, and facilitate invasion [1–5]. Excessive ECM deposition, known as fibrosis, is commonly observed in chronic infectious diseases caused by various microbial agents, including bacteria, viruses such as HIV, influenza, and SARS-CoV-2, as well as parasites like *Leishmania* [5–7]. Therefore, regulating collagen turnover in the ECM has been proposed as a potential therapeutic strategy. Similarly, in *Caenorhabditis elegans*, the cuticle—a specialized extracellular matrix primarily composed of collagen—plays a crucial role in innate immunity and aging, shielding the organism from environmental stressors and microbial infections [8–12]. Recent studies have shown that neuronal signaling via a G protein-coupled receptor (GPCR) regulates cuticle barrier defense, suggesting that the nervous system can rapidly transmit signals to the epidermis to modulate ECM remodeling in response to environmental stimuli [12]. However, the underlying mechanisms of this neuron-epidermis communication for ECM remodeling remain largely unknown.

Lysosomes play a pivotal role in regulating collagen biosynthesis, cuticle turnover, and tissue repair. In mammals, lysosomal enzymes are essential for extracellular matrix (ECM) remodeling, ensuring balanced collagen degradation and synthesis. For instance, proteolytic collagen fragments can be internalized via the collagen receptor urokinase plasminogen activator receptor-associated protein (uPARAP/Endo180) and subsequently degraded in lysosomes by intracellular cathepsins [13–15]. Moreover, cancer cells exploit lysosomal enzyme secretion to degrade ECM components, facilitating tissue invasion [16]. Additionally, lysosomal function has been implicated in collagen production regulation via mTORC1 signaling, suggesting an indirect role in maintaining ECM composition and quality [17]. In *Caenorhabditis elegans*, lysosomes in the epidermis are activated at each molt when the apical ECM (cuticle) is replaced, ensuring proper ECM turnover and developmental progression [18]. Disruptions in lysosomal function impair endocytic cargo degradation, suppress protein synthesis required for molting, and result in molting defects [18]. These findings underscore the necessity of lysosomal activity for cuticle remodeling and epidermal integrity during development. However, the role of lysosomes in epidermal collagen remodeling under pathogen-induced stress remains poorly understood. Given the critical function of lysosomes in ECM maintenance, elucidating their regulation during infection is essential for understanding host defense mechanisms.

Neuronal signaling has emerged as a crucial regulator of epidermal immunity and collagen homeostasis. In mammals, pathogens can directly activate sensory neurons to release neuropeptides that contribute to skin immunity. For example, bacterial toxins such as α-toxin/hemolysin (Hla), phenol-soluble modulins (PSMs), and leukocidin HlgAB from *Staphylococcus aureus* and streptolysin S (SLS) from *Streptococcus pyogenes* have been shown to induce pain and inflammatory responses by directly activating sensory neurons [19–21]. Sensory neurons also play a fundamental role in regulating innate immunity and tissue repair across species. In *Caenorhabditis elegans*, sensory neurons such as ASI and ASH integrate environmental cues to modulate immune responses and enhance pathogen defense [22]. Neuronal signaling pathways, including the GPCRs OCTR-1 and NPR-1, as well as neuropeptide-mediated communication, contribute to the regulation of immune responses [22–25]. Traditionally, the cuticle has been regarded as a static physical barrier providing innate defense against pathogens [26]. However, recent studies indicate that *C. elegans* actively remodels its cuticle structure in response to infection. Notably, neuronal GPCRs such as NPR-8 have been shown to inhibit the expression of cuticle collagen, suggesting a dynamic regulation of barrier defenses through neuronal signaling [12]. Understanding how the nervous system modulates ECM remodeling via neuroendocrine signaling will provide valuable insights into host-pathogen interactions and the regulation of epidermal defenses.

In this study, using the *Pseudomonas aeruginosa* PA14–*Caenorhabditis elegans* infection model, we investigate how neuronal signaling regulates epidermal lysosomal activity, which in turn modulates collagen remodeling. We discover that *Pseudomonas aeruginosa* PA14 infection disrupts the structural integrity of the *Caenorhabditis elegans* cuticle by impairing lysosomal function and that this disruption is mediated by pathogen-induced neuronal signaling. Our findings reveal a novel neuro-epidermal pathway in which a neuronally secreted factor, NSIF-1, acts to inhibit epidermal lysosomal activity through suppression of the transcription factor ELT-3, leading to compromised collagen organization and increased host susceptibility to infection.

## Results

### PA14 compromises nematode cuticle collagen and triggers epidermal immunity

The nematode cuticle functions similarly to human skin as a physical barrier and first line of immune defense. This protective structure, composed primarily of collagen proteins secreted by epidermal cells, forms a complex extracellular matrix (ECM) [27, 28]. We investigated whether PA14 directly damages cuticle collagen and thereby triggers an immune response in the epidermis.

We first validated the infection protocol by initiating infections at the L4 stage (48 h post-L1), thereby bypassing the earlier larval molts (L1-L3). L4-stage synchronization was achieved using vulval substaging criteria [29] (S1A Fig). Following 24 h of exposure to PA14, we evaluated multiple developmental parameters, including body length and egg morphology. No significant differences were observed between PA14-exposed and control animals for either parameter (S1B-S1C Fig), confirming that the 24-h PA14 exposure protocol does not cause detectable developmental delay.

DPY-7 collagen, secreted by epidermal cells and deposited into the cuticle’s annular furrows [30], serves as a reliable marker for cuticle morphology when tagged with GFP (DPY-7::sfGFP) [18]. To determine whether PA14 affects the intact cuticle or specifically targets molting phases, we performed a time-course experiment. Cuticle morphology was visualized using DPY-7::sfGFP, and molting status was assessed at five time points: L4+0h, L4+3h, L4+6h, L4+12h, and L4+24h (S2A Fig). Molting was identified by the presence of a mouth plug—a distinctive cap-like structure formed by the loosening of the old cuticle, which temporarily halts feeding [31]. At early intervals (L4+3h), both OP50-fed and PA14-infected animals exhibited mouth plugs, indicating active molting. During these periods, cuticle damage characterized by fragmentation and disorganized DPY-7 patterning was observed in both conditions, reflecting normal cuticle remodeling during molting. At 12 hours post-PA14 exposure, cuticle abnormalities began to appear, while animals were not in molting phases (no mouth plugs observed). By 24 hours infection, severe cuticle damage (fragmentation and loss of DPY-7 patterning) was evident, again occurring outside molting windows (Figs 1A, S2B-S2E). These observations indicate that PA14 directly compromises the structural integrity of DPY-7 in the cuticle. The phenotype emerging at 24 hours post-L4 suggests an active pathogenic mechanism that disrupts cuticle homeostasis independent of developmental remodeling.

**Figure 1.**
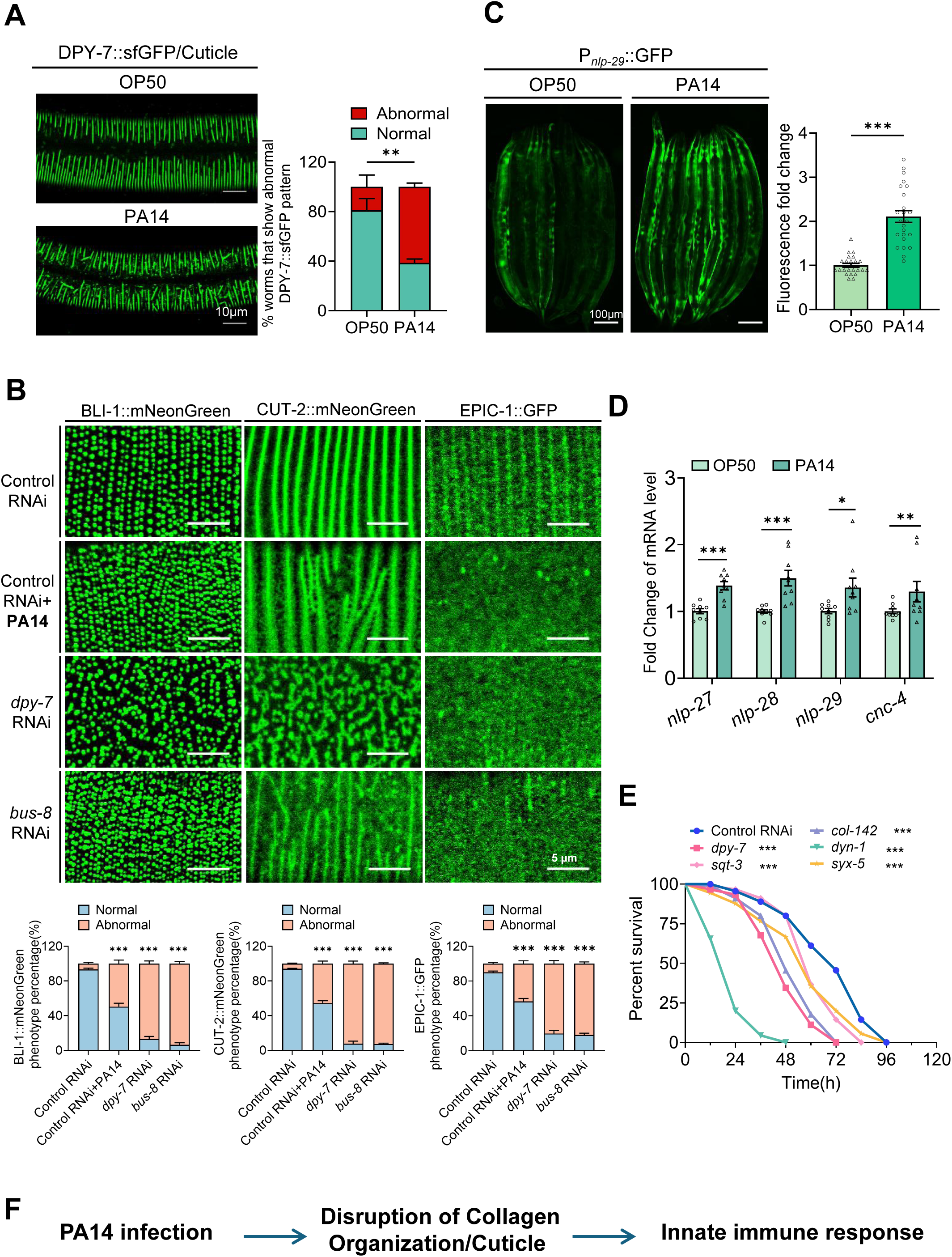
*Pseudomonas aeruginosa* PA14 disrupts cuticle collagen integrity and activates epidermal immunity in *C. elegans*. **(A)** Confocal fluorescence images of DPY-7::sfGFP in wild-type (WT) young adults after 24h PA14 exposure (infection initiated at L4 stage), and the quantification of worms with abnormal DPY-7 pattern is calculated in 300 worms (n=300). Scale bar:10µm. **(B)** Confocal fluorescence images and quantitative analysis of BLI-1::mNeonGreen (n=300 animals), CUT-2::mNeonGreen (n=300 animals) and EPIC-1::GFP (n=300 animals) in young adults animals after 24h PA14 exposure (infection initiated at L4 stage). RNAi knockdown of *dpy-7* and *bus-8 act as* positive control, which resulted in abnormal patterning and mislocalization of all markers examined. Scale bars: 5 µm. **(C)** Representative microscope images and quantitative analysis of P*_nlp-29_*::GFP fluorescence intensity in L4-stage animals following 24-hour exposure to PA14. n=24 worms, Scale bar:100µm. **(D)** qRT-PCR analysis of antimicrobial peptide (AMP) gene expression (*nlp-27, nlp-28, nlp-29, cnc-4*) in L4 wild-type animals following 24 hours exposure to OP50 or PA14 exposure. **(E)** Survival analysis of wild-type worm fed either control RNAi bacteria or RNAi bacteria targeting *dpy-7*, *sqt-3*, *col-142*, *dyn-1*, and *syx-5* in the PA14 slow-killing assay. PA14 survival assays were performed in three independent replicates, detailed statistical data are presented in source data. **(F)** Schematic model of PA14 infection in disrupting collagen organization. For all quantification, data are presented as mean ± SEM. Statistical comparisons were performed using unpaired two-tailed Student’s t-test (A: worms with abnormal DPY-7 pattern in OP40 vs PA14.), one-way ANOVA followed by Tukey’s multiple comparisons test (B: indicated RNAi and PA14 treatment vs control RNAi), Mann–Whitney U test (B) and Log rank (Mantel-Cox) test between indicated RNAi vs control RNAi (E), * p<0.05, **p< 0.01, ***p<0.001, ns., not significant.. All experiments were performed independently at least three times. The data underlying this figure can be found in **S1 Data**.

To rule out the possibility that the observed effects reflect a time dependent impact on cuticle biosynthesis rather than direct damage, we performed an additional experiment using young adults (L4+24 h, well beyond the L4 to adult molt) expressing DPY 7::sfGFP. These animals were transferred to either OP50 (control) or PA14 lawns and imaged at 0, 3, 6, 12, and 24 h post transfer (S3A Fig). PA14-exposed adults exhibited a significant increase in animals with disorganized DPY-7 pattern as early as 3 h post-transfer, and this proportion increased progressively at 6, 12, and 24 h (S3B-S3E Fig). Since the animals were already post molt young adults at the time of transfer, these results rule out any artifacts related to molting or cuticle assembly and clearly demonstrate that PA14 directly disrupts cuticle homeostasis.

The cuticle of *C. elegans* is a complex extracellular matrix composed of distinct layers, each characterized by specific structural proteins that collectively maintain barrier function and mechanical stability [32]. To determine whether *P. aeruginosa* PA14-induced cuticle damage is specific to DPY-7 or broadly affects cuticular architecture, we examined markers representing multiple layers. Using translational fusions for CUT-2 (cortical layer)[33], BLI-1 (struts)[34] and EPIC-1 (epicuticle) [35], we assessed cuticle integrity following 24-hour PA14 infection from L4 stage. In control animals fed *E. coli*, all markers displayed organized patterns consistent with their established localizations. As a positive control, RNAi knockdown of *dpy-7* and *bus-8*—genes essential for maintaining the cuticle barrier [36]—resulted in abnormal patterning and mislocalization of all markers examined (Fig 2B). After 24 hours of PA14 infection, we observed disrupted patterning and mislocalization of all markers tested (Fig 2B). These results indicate that the pathogenic effect extends beyond DPY 7 and broadly impairs multiple cuticular layers.

**Figure 2.**
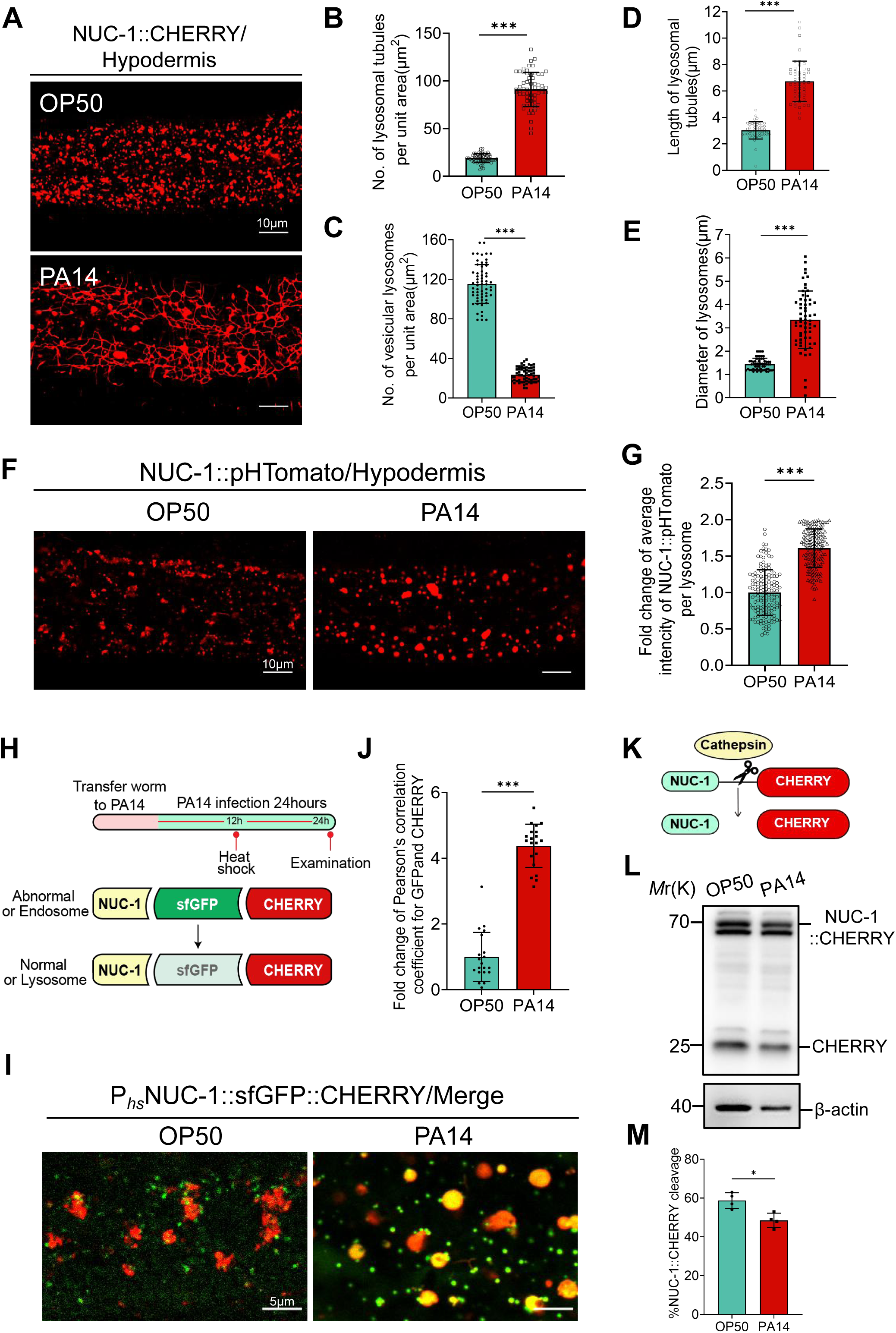
PA14 infection disrupts cuticle structure by impairing lysosomal function. **(A)** Confocal fluorescence images of epidermal lysosomes in young adults NUC-1::CHERRY animals after 24h PA14 exposure (infection initiated at L4 stage). Scale bar:10µm **(B)** As shown in (A), the number of tubular lysosomes was quantified by counting within three 35×25(µm^2^) unit areas per individual worm, with a total of 20 worms analyzed (n=20 worms). **(C)** As shown in (A), the number of vesicular lysosomes was quantified by counting within three 35×25(µm^2^) unit areas per individual worm, with a total of 20 worms analyzed (n=20 worms). **(D)** As shown in (A), the length of tubular lysosomes was quantified by measuring the longest five tubular lysosomes within a 35×25(µm^2^) unit areas per individual worm, with a total of 10 worms analyzed (n=10 worms). **(E)** As shown in (A), the diameter of vesicular lysosomes was quantified by measuring the six largest vesicular lysosomes within a 35×25(µm^2^) unit areas per worm, with a total of 10 worms analyzed (n=10 worms). **(F-G)** Confocal fluorescence imaging (F) and quantitative analysis (G) were performed on the hypodermis of young adults wild-type (WT) expressing NUC-1::pHTomato under the control of a heat-shock promoter after 24h PA14 exposure (infection initiated at L4 stage). Fluorescence quantification was performed on 150 lysosomes from 20 worms (n=20 worms). Scale bar:10µm. **(H-J)** Merged confocal fluorescence images (I) of the epidermis were acquired from young adults wild-type worm expressing NUC-1::sfGFP::CHERRY at 12 hours post-heat shock (HS) following exposure to OP50 or PA14 24h (infection initiated at L4 stage). A schematic diagram of the experiment is presented in (H), and Pearson’s correlation coefficient for GFP and CHERRY fluorescence is shown in (J), and least 20 worms were analyzed (n=20 worms). Scale bar:5µm. **(K)** A schematic diagram of cleavage of CHERRY from NUC-1::CHERRY. **(L-M)** Immunoblots (L) and quantification (M) showing cleavage of CHERRY from NUC-1::CHERRY transgenic young adults animals after 24h PA14 exposure (infection initiated at L4 stage). N=4 independent biological replicates in (L); For all quantification, data are presented as mean ± SEM. Statistical comparisons were performed using Mann–Whitney U-test (B, C, D, E, G, M). **p<0.01, ***p<0.001. All experiments were performed independently at least three times. The data underlying this figure can be found in **S1 Data**.

To directly assess cuticle barrier integrity, we performed a quantitative cuticle permeability assay using Hoechst 33342 dye penetration [37]. Consistent with published data [36], RNAi knockdown of *bus-8* and *dpy-7* resulted in a significant increase in Hoechst 33342 fluorescence intensity in the head and tail regions of the animals (S4A Fig). Our findings show that 24-hour PA14 infection significantly increases Hoechst 33342 fluorescence intensity in the head and tail regions of wild-type animals compared with uninfected controls (S4A Fig). This enhanced dye penetration provides direct physiological evidence that PA14 infection compromises the structural integrity of the cuticle barrier.

Since damage to the nematode epidermis is known to induce an immune response—specifically through the upregulation of the antimicrobial peptide NLP-29 [38, 39], we hypothesized that PA14-induced cuticle damage might similarly trigger epidermal immunity. To test this, we monitored the expression of the epidermal immunity reporter P*_nlp-29_*::GFP and found a significant increase in fluorescence following PA14 infection (Fig 1C). This finding was further corroborated by elevated expression levels of several antimicrobial peptide (AMP) genes in the epidermis, including *nlp-27*, *nlp-28*, *nlp-29*, and *cnc-4* (Fig 1D).

To further explore the relationship between cuticle integrity and epidermal immunity, we disrupted cuticle synthesis and structural organization using RNA interference (RNAi) against several key genes *dpy-7* [36], *sqt-3* [40], *col-142* [41], *dyn-1* and *syx-5* [18, 42]. The results demonstrated that compromising cuticle integrity via gene knockdown led to an upregulation of epidermal immunity (S4B Fig). Notably, knockdown of *dpy-7*—a major furrow collagen [36], and *dyn-1*—a gene essential for the endocytosis [43] —resulted in the most significant immune activation (S4B Fig). Furthermore, we found that directly knocking down genes related to cuticle synthesis and assembly also reduced the survival rate of nematodes when exposed to PA14 (Fig 1E). This highlights the critical role of cuticle integrity in protecting nematodes against PA14 infection.

In summary, our findings extend the established paradigm of cuticle-immune crosstalk in nematodes [44–46] by demonstrating that pathogens like *P. aeruginos*a PA14 infection damages the structural integrity of the nematode cuticle, which in turn triggers an upregulation of epidermal immunity (Fig 1F).

### PA14 infection disrupts cuticle structure by impairing epidermal lysosomal function

A previous study revealed that enhanced epidermal lysosomal activity is crucial for cuticle collagen turnover during the molting process in *C. elegans* [18]. Certain pathogens evade host defenses by secreting proteins that inhibit V-ATPase, thereby disrupting lysosomal acidification, fusion, and overall function [47–50]. Given that PA14 infection causes aberrant DPY-7 localization (Fig 1A), we hypothesized that PA14 impairs lysosomal function, which in turn damages the cuticle. To test this hypothesis, we examined several lysosomal properties—morphology, acidification, and degradation activity—in animals infected with PA14.

Firstly, we evaluated lysosomal morphology using an established reporter-*qxIs383*(P*_hyp-7_*::NUC-1::CHERRY [18, 51] after 24 hours of PA14 infection. We observed a significant increase in tubular lysosomal structures under PA14 treatment (Fig 2A-B). We also observed a decrease in the number of vesicular lysosomes (Fig 2C), an increase in the length of tubular lysosomes (Fig 2D), and an enlargement in lysosomal diameter (Fig 2E) under PA14 infection.

Secondly, to assess lysosomal acidification, we employed an established transgenic reporter, NUC-1::pHTomato [51, 52], in which the pH-sensitive fluorescent protein pHTomato (with a pKa near 7.8) is fused to NUC-1 under the control of a heat-shock promoter. pHTomato exhibits increased fluorescence at higher pH. We found that animals infected with PA14 displayed significantly higher NUC-1::pHTomato fluorescence compared to those treated with OP50 (Fig 2F-2G), indicating that PA14 infection leads to an increase in lysosomal pH, thereby disrupting acidification—a critical condition for the activity of lysosomal hydrolases.

Thirdly, we monitored lysosomal maturation using the transgenic reporter-*qxIs612*(P*_hs_*NUC-1::sfGFP::CHERRY) [18, 51]. In this system, a heat shock at 34°C for 30 minutes induces the expression of the NUC-1::sfGFP::CHERRY fusion protein, which is then trafficked to lysosomes via the endocytic pathway. GFP fluorescence was quenched in the acidic lysosomal environment, while CHERRY fluorescence remains stable, allowing us to assess lysosomal acidification and maturation by the loss of GFP signal (Fig 2H). In PA14-infected animals, GFP fluorescence was not quenched and colocalized strongly with CHERRY (Fig 2I-2J), indicating impaired lysosomal maturation and acidification.

Finally, we evaluated lysosomal degradation activity using the transgenic reporter-*qxIs383*(P*_hyp-7_*NUC-1::CHERRY) [18], which expresses the NUC-1::CHERRY fusion protein in the epidermis. Upon delivery to lysosomes, CHERRY is cleaved by cathepsins, and the cleaved product can be quantified by western blot (Fig 2K). We observed a reduction in cleaved CHERRY levels following PA14 infection (Fig 2L-2M), demonstrating a decline in lysosomal degradation activity.

To investigate the role of specific bacterial virulence factors in modulating lysosomal morphology and cuticle collagen organization, we compared the effects of wild-type *P. aeruginosa*-PA14 with those of attenuated mutant strains lacking key virulence regulators (Δ*gacA,* Δ*lasR,* Δ*mvfR,* Δ*pqsF*) [53]. In contrast to wild-type PA14, these mutants significantly attenuated the PA14-induced lysosomal alterations and cuticle disruption (S5A-S5B Fig), indicating that functional virulence expression is required for PA14-dependent dysregulation of both lysosomal architecture and cuticle integrity.

In summary, our findings reveal that PA14 infection drives lysosomes into an inactive state characterized by altered morphology, reduced acidification, impaired maturation, and diminished degradation activity. This impairment of lysosomal function likely contributes to the disruption of cuticle collagen, thereby compromising the nematode’s physical barrier and triggering epidermal immunity.

### NSIF-1, expressed in neuron, is a signaling factor that responds to PA14 infection and modulates lysosomal and cuticle integrity

*C. elegans* uses various sensory neurons to detect pathogenic bacteria, triggering both avoidance behaviors and immune responses [54]. Neuronal signaling also influences peripheral mitochondrial homeostasis, affecting aging, lipid metabolism, and immunity [55–58]. However, whether neuronal signals can regulate lysosomal functions in peripheral tissues remains unclear.

Given that PA14 infection disrupts epidermal lysosomal function (Fig 2A) and collagen integrity (Fig 1A), an important question arises: Does the nervous system relay signals to epidermal lysosomes, thereby exacerbating cuticle damage during infection? Since secreted proteins are crucial mediators of intercellular communication—as hormones, cytokines, or ligands binding to receptors [59], we hypothesized that PA14 infection might induce the secretion of specific proteins that impair lysosomal function (Fig 3A).

**Figure 3.**
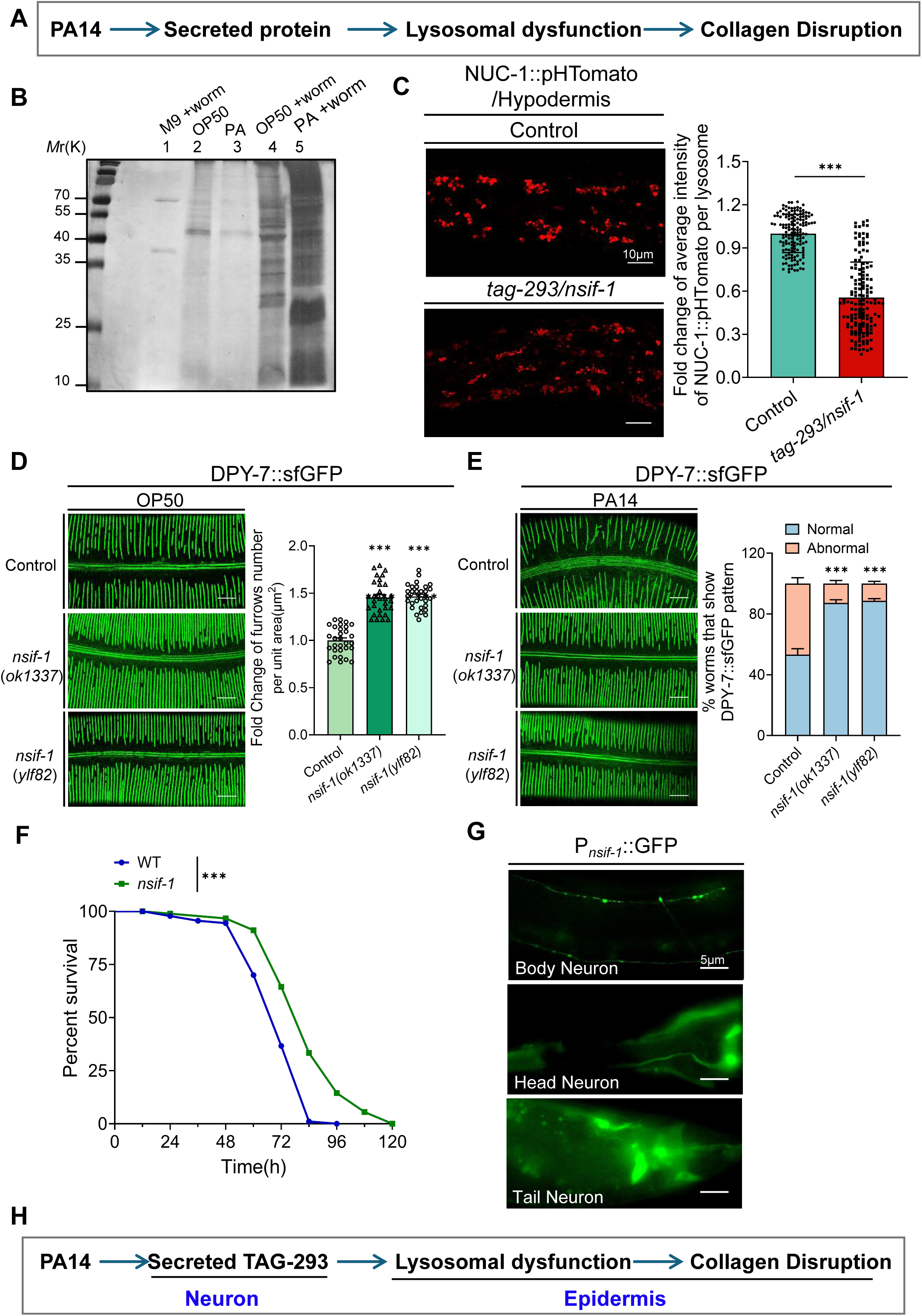
NSIF-1, expressed in neuron, is a signaling factor that responds to PA14 infection and modulates lysosomal and cuticle integrity. **(A)** Simple diagram of the strategy for identifying secreted proteins that affect lysosome function and subsequently influence cuticle morphology. **(B)** Silver staining of secreted proteins from wild-type (WT) animals under different treatment conditions. L1-stage animals were grown in liquid culture for 4 hours (lane 1) or exposed to *E. coli* OP50 (lane 4) or *P. aeruginosa* PA14 (lane 5). Control lanes 2 and 3 contained the same liquid culture treated with OP50 or PA14, respectively, but without animals. After treatment, animals and bacteria were removed by centrifugation, and the medium was filtered through a 0.22μm filter to obtain secreted proteins for sliver staining analysis. **(C)** Confocal fluorescence imaging and quantitative analysis were performed on the hypodermis of wild-type (WT) and *tag-293/nsif-1(ok1337)* mutant expressing NUC-1::pHTomato under the control of a heat-shock promoter (young adults wild-type after 24h PA14 exposure and infection initiated at L4 stage). The average pHTomato fluorescence intensity was performed on 150 lysosomes from 20 worms (n=20 animals, total 150 lysosomes). Scale bar:10µm. **(D)** Confocal fluorescence images and quantitative analysis were performed on the hypodermis of wild-type (WT), *tag-293/nsif-1(ok1337)* and *tag-293/nsif-1(ylf82)* expressing DPY-7::sfGFP, all of these animals are adult under OP50 condition. Quantification of cuticle furrows density in wild-type (WT), *nsif-1(ok1337)* and *nsif-1(ylf82)* were performed on 30 worms (n=30 animals). Scale bar:10µm. **(E)** Confocal fluorescence images and quantitative analysis were performed of DPY-7::sfGFP in young adults wild-type (WT) and *nsif-1(ok1337)* mutant after 24h PA14 exposure (infection initiated at L4 stage). The statistical analysis of worms with abnormal DPY-7 pattern in WT and *nsif-1(ok1337)* after 24h PA14 exposure performed on 300 worms (n=300 animals). Scale bar:10µm. **(F)** Survival analysis of wild-type (WT) and *nsif-1(ok1337)* mutant in the PA14 slow-killing assay. PA14 survival assays were performed in three independent replicates, detailed statistical data are presented in source data. **(G)** Confocal fluorescence imaging of P*_nsif-1_*::GFP. GFP was expressed in various neuron, particularly within the head region. Scale bar:5µm. **(H)** The simple diagram of the conclusive findings from this section. PA14 induced neuronal secreted NSIF-1 may disrupt epidermal lysosomal function and collagen organization. For all quantification, data are presented as mean ± SEM. Statistical comparisons were performed using Mann–Whitney U-test (C), one-way ANOVA followed by Tukey’s multiple comparisons test (D, E) and Log rank (Mantel-Cox) test (F). ***p<0.001. All experiments were performed independently at least three times. The data underlying this figure can be found in **S1 Data**.

To identify these factors, we collected supernatants from worms treated with PA14 and various control conditions (M9 buffer+worms; OP50+worms; OP50 only; PA14 only). Silver staining revealed distinct protein bands in PA14-treated samples (Fig 3B), suggesting that infection stimulates the secretion of certain proteins. Mass spectrometry identified 35 candidate proteins induced by PA14 (S6A Fig, S1 Table). By applying two criteria—(1) the presence of signal peptides (as predicted by SignalP-5.0) [60] and (2) neuronal expression based on the *C. elegans* Neuronal Gene Expression Map & Network (CeNGEN https://www.cengen.org/) database [61], we narrowed this list to 5 potential neuronal secreted proteins (S6A Fig).

We then investigated whether any of these PA14-induced proteins impair lysosomal function, consequently affecting collagen organization (Fig 3A). If PA14-stimulated neuronal proteins disrupt lysosomal activity, then knocking them down should restore lysosomal function. Using RNAi to reduce the expression of *rla-5, dao-2, nlp-77, tag-293* and *lec-5,* we assessed lysosomal acidification with NUC-1::pHTomato. Remarkably, knockdown of *tag-293* significantly enhanced lysosomal acidification under PA14 infection (S6B Fig). Consistently, the *tag-293(ok1337)* deletion mutant exhibited elevated lysosomal acidification (Fig 3C). Furthermore, analysis of supernatants from worms expressing TAG-293::Flag::GFP showed that PA14 infection indeed promotes the secretion of TAG-293 (S6C Fig). These data imply that PA14-induced TAG-293 secretion may impair lysosomal function. Given that TAG-293 is a neuron-derived secreted factor that responds to pathogen infection and likely plays a role in immune regulation, we have renamed TAG-293 as **NSIF-1** (**<u>N</u>**euronal **<u>S</u>**ecreted **<u>I</u>**mmune **<u>F</u>**actor-1).

Next, we assessed whether the enhanced lysosomal activity in *nsif-1* mutants impacts cuticle collagen integrity. We generated a second null-like allele, *nsif-1(ylf82)*, using CRISPR/Cas9 to introduce a stop codon in the second exon (S7A Fig). Under normal OP50 feeding conditions, two independent *nsif-1* mutant alleles (*ok1337, ylf82*) displayed a higher density of cuticular structural units compared to wild-type worms (Figs 3D, S7B-S7D). Moreover, during PA14 infection, the disrupted collagen organization observed in wild-type worms was significantly alleviated in *nsif-1* mutants (Figs 3E, S7B-7D).

Survival assays further confirmed that *nsif-1(ok1337)* mutants exhibit enhanced resistance to PA14 infection relative to wild-type animals (Figs 3F, S8). In contrast, overexpression of NSIF-1 under its native promoter in wild-type animals did not result in increased susceptibility to PA14 (S8 Fig), indicating that only pathogen-induced NSIF-1 expression contributes to infection progression. This supports the idea that PA14-induced NSIF-1 contributes to cuticle damage.

To determine the expression and localization of NSIF-1, we generated transcriptional (P*_nsif-1_*::GFP) and translational (P*_nsif-1_*NSIF-1::GFP) reporters. The transcriptional reporter confirmed that *nsif-1* is exclusively expressed in neurons (Fig 3G), whereas the translational reporter revealed that the NSIF-1 protein is distributed across various tissues, including head and body neurons, muscle, and intestine (S9A Fig). Using a CRISPR knock-in strain (*syb9508*) expressing NSIF-1::mNeonGreen::3xFLAG from the endogenous locus, we further confirmed that NSIF-1 also localizes to the epidermis (S9B Fig). Together, these findings indicate that neuronally expressed NSIF-1 may be secreted and act on peripheral tissues, likely modulating epidermal lysosomal function and collagen organization (Fig 3H).

Under OP50 feeding conditions, this NISF-1 (endogenous reporter: NSIF-1::mNeonGreen::3xFLAG) exhibited low-level fluorescence. However, upon 24-hour infection with wild-type *P. aeruginosa*-PA14, we observed a marked increase in fluorescence intensity specifically within the epidermis (S10A Fig). Furthermore, the NSIF-1::mNeonGreen signal showed clear co-localization with epidermal lysosomal markers (S10B Fig). These effects were significantly attenuated when infection was carried out with attenuated PA14 strains lacking key virulence factors (*gacA, lasR, mvfR, pqsF*) (S10B Fig). These results confirm that pathogen-induced upregulation and epidermal lysosomal localization of NSIF-1 are specific responses to active virulence.

In summary, we propose that PA14 infection triggers the neuronal secretion of NSIF-1—a signaling factor that modulates epidermal lysosomal function and cuticle integrity (Fig 3H). This disruption of lysosomal activity leads to impaired collagen organization, compromising the cuticle barrier and increasing susceptibility to PA14 infection.

### PA14 infection induces the secretion of NSIF-1 from neurons to the epidermis

To directly demonstrate that NSIF-1 is synthesized in neurons and secreted to epidermis, we employed a GFP-binding nanobody (GBP) capture system in epidermis. In this system, tissue-specific expression of GBP::mKate2 enables detection of trace amounts of secreted GFP-tagged proteins originating from other tissue [62, 63]. We generated transgenic *C. elegans* strains expressing GBP::mKate2 in the epidermis (P*_col-12_*GBP::mKate2) in a background where NSIF-1::GFP is expressed pan-neuronally under the P*rgef-1* promoter (P*_rgef-1_*NSIF-1::GFP). If NSIF-1 is secreted from neurons and reaches the epidermis, the epidermal GBP should capture NSIF-1::GFP, resulting in detectable green fluorescence (Fig 4A).

**Figure 4.**
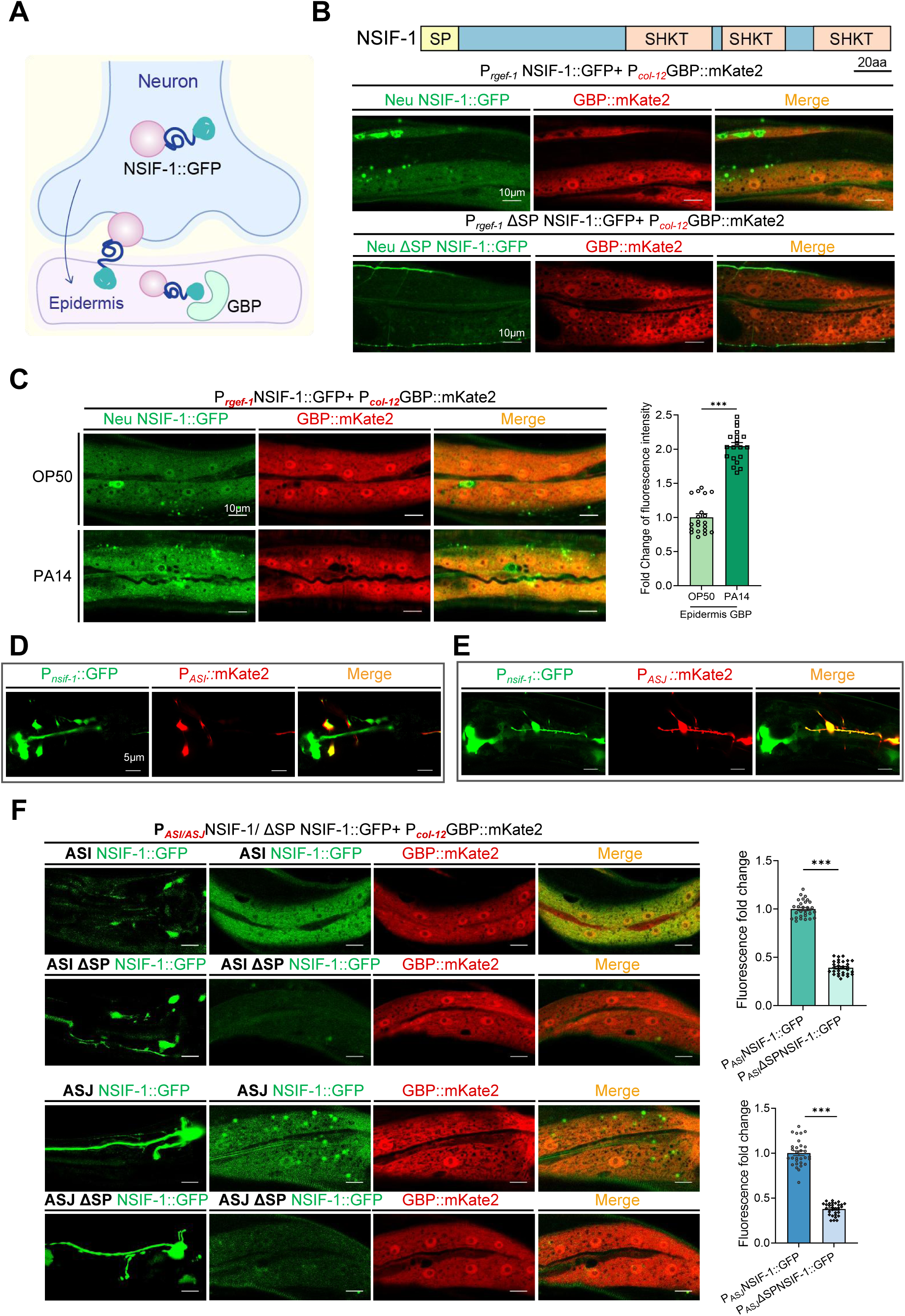
PA14 infection promotes neuronal-derived NSIF-1 release to epidermis and intestine. **(A)** Schematic illustrating how GBP::mKate, expressed in non-neuronal tissues (epidermis), captures neuronal secreted NSIF-1::GFP. **(B)** Confocal fluorescence imaging of transgenic animals expressing NSIF-1::GFP (or the signal-peptide deletion mutant, ΔSP-NSIF-1::GFP) in pan-neurons (P*rgef-1*) and GBP::mKate in the epidermis (P*col-12*). While the secreted NSIF-1::GFP was robustly detected in the epidermis (green), the ΔSP abolished this epidermal localization, confirming the requirement of the signal peptide for secretion. Scale bar:10µm. **(C)** Confocal fluorescence imaging of transgenic animals co-expressing pan-neuronal (P*rgef-1)* NSIF-1::GFP and epidermal (P*col-12*) GBP::mKate2. Following 24 h of PA14 exposure (initiated at the L4 larval stage), secreted neuronal NSIF-1::GFP (green) was more abundantly detected in the epidermis of young adult animals. Quantification of epidermal NSIF-1::GFP fluorescence intensity after PA14 exposure is shown (n=20 animals). Scale bar:10µm. **(D-E)** Analysis of the localization distribution of P*_nsif-1_*::GFP in ASI(P*_ASI (str-3)_*::mKate2) (D) and ASJ (P*_ASJ(trx-1)_*::mKate2) (E) neurons for adult animals. Scale bars, 5µm. **(F)** Confocal fluorescence imaging of the transgenic line expressing NSIF-1::GFP and ΔSPNSIF-1::GFP in ASI(P*_str-3_*)/ASJ(P*_trx-1_*) and GBP::mKate2 in epidermis(P*_col-12_*). Expression of ΔSPNSIF-1 in ASJ and ASI neurons abolished the secretion of NSIF-1::GFP into the epidermis. Quantification of epidermal NSIF-1::GFP fluorescence intensity after PA14 exposure is shown (n=30 animals). Scale bar:10µm. For all quantification, data are presented as mean ± SEM. Statistical comparisons were performed using Mann–Whitney U-test (C, F). ***p<0.001. All experiments were performed independently at least three times. The data underlying this figure can be found in **S1 Data**.

Our results confirmed that GBP::mKate2 expressed in the epidermis successfully captured neuron-derived NSIF-1, producing clear green fluorescence signals in these tissues (Fig 4B). In contrast, when the signal peptide (SP) of NSIF-1 was removed, no green fluorescence was detected in epidermis, despite the presence of GBP::mKate2 (Fig 4B). These results demonstrate that NSIF-1 is indeed secreted from neurons and taken up by the epidermis. Furthermore, upon PA14 infection, we observed a marked increase in NSIF-1 signal specifically in the epidermis (Fig 4C), indicating that infection induces enhanced secretion of NSIF-1 from neurons to epidermis. These effects were suppressed when worms were infected with attenuated PA14 strains-lacking key virulence regulators (*gacA*, *lasR*, *mvfR*, *pqsF*) (S11 Fig), indicating that NSIF-1 secretion is specifically dependent on PA14 virulence factors.

To determine whether NSIF-1 secretion is a specific response to PA14 virulence rather than a general response to cuticle damage, we disrupted cuticle integrity using RNAi against collagen and extracellular matrix genes (*col-142, dpy-7, dyn-1, sqt-3, syx-5*). None of these RNAi induced NSIF-1 secretion into epidermis (S12 Fig), confirming that NSIF-1 is specifically upregulated in response to PA14 virulence factors rather than general barrier damage.

We further investigated whether NSIF-1 secretion depends on canonical neuropeptide processing pathways by testing in animals with RNAi of *unc-31*(regulating dense-core vesicle secretion) [64] and *egl-3* (proprotein convertase) [65]. NSIF-1 secretion remained unchanged in these RNAi animals (S13 Fig), indicating that its release occurs through a non-canonical pathway independent of traditional neuropeptide processing mechanisms.

To identify which specific neurons express *nsif-1* and thus secret it to the epidermis, we first examined its expression profile. Analysis using the *C. elegans* Neuronal Gene Expression Map & Network (CeNGEN) database revealed that *nsif-1* is predominantly expressed in ASJ neurons (S6A Fig). Subsequent co-localization studies, using transcriptional (P*_nsif-1_*::GFP) and translational (P*_nsif-1_*NSIF-1::GFP) reporters along with multiple neuronal markers, confirmed that nsif-1 is expressed exclusively in ASI and ASJ neurons, rather than in RIS or ADF neurons. (Figs 4D-E, S14).

To test ASI/ASJ (neuron)-to-hypodermis NSIF-1 transfer, we generated four transgenic lines expressing full-length or ΔSP NSIF-1::GFP under ASI-(*str-3*) or ASJ-specific (*trx-1*) promoters, in a hypodermal GBP capture background (P*_col-12_*GBP::mKate2). Full-length yielded robust hypodermal GFP signal, whereas ΔSP yielded little to none (Fig 4F). These results directly demonstrate that NSIF-1 produced specifically in ASI or ASJ neurons is secreted and taken up by the hypodermis. Thus, ASI/ASJ-derived NSIF-1 may influence epidermal lysosomal function.

### NSIF-1 enhances susceptibility to PA14 by inhibiting lysosomal function

To investigate how NSIF-1 modulates immune responses and lysosomal activity in *C. elegans*, we performed transcriptome analysis (RNA-seq) on wild-type N2 worms and *nsif-1(ok1337)* mutants. KEGG pathway enrichment analysis of the upregulated genes in *nsif-1(ok1337)* mutants revealed a significant enrichment of lysosomal-related genes—54 in total (S15A-S15B Fig, S2 Table, S3 Table). Basic categorization showed that among these genes, 32 encode membrane proteins, 12 encode proteases, and 6 encode non-hydrolytic enzymes (S15B Fig, S3 Table). Subsequent qPCR analysis confirmed the RNA-seq results (S15C Fig), demonstrating that genes related to lysosomal proteases and non-hydrolytic enzymes were significantly upregulated. These data suggest that lysosomal function is enhanced in the *nsif-1* mutant.

We next examined lysosomal morphology under normal OP50 feeding and PA14 infection conditions. In wild-type animals, PA14 infection increased the number of tubular lysosomes and decreased the number of vesicular lysosomes compared to OP50-fed controls; however, these morphological changes were restored in the *nsif-1* mutant background (Fig 5A-C). Moreover, by measuring lysosomal degradation activity using *qxIs383*(P*_hyp-7_*NUC-1::CHERRY), we found that cleaved CHERRY is increased in *nsif-1(ok1337)* mutant animals compared with wild-type under *E. coli* (Fig 5D) or PA14 (Fig 5E) condition, indicating an increased lysosomal degradation activity in *nsif-1*mutant animals.

**Figure 5.**
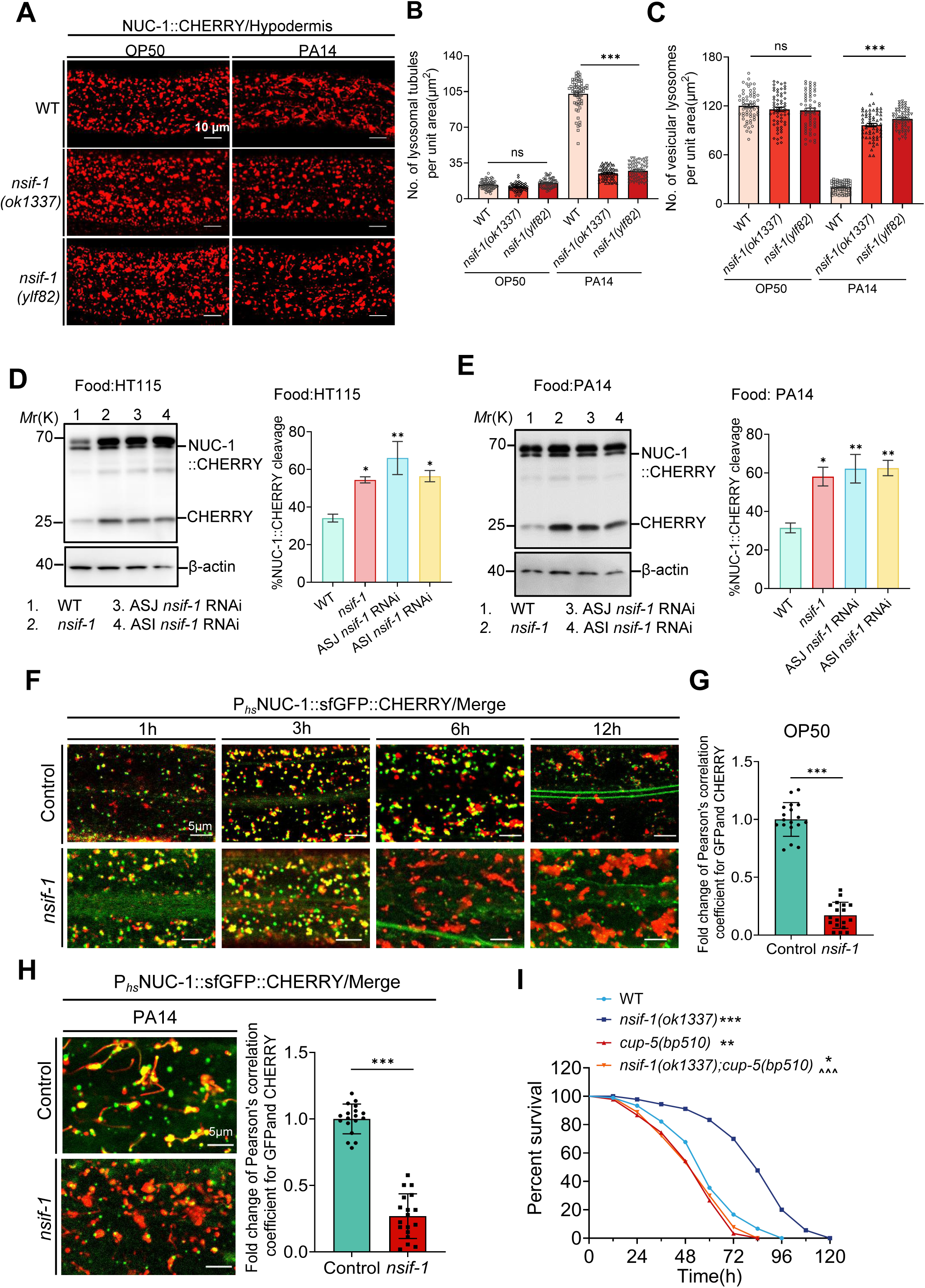
NSIF-1 enhances susceptibility to PA14 by inhibiting lysosomal Function. **(A-C)** Confocal fluorescence images (A) of lysosomes in the epidermis of young adults wild-type (WT), *nsif-1(ok1337)* and *nsif-1(ylf82)* expressing NUC-1::CHERRY after 24h PA14 exposure (infection initiated at L4 stage). The number of tubular lysosomes (B) and vesicular lysosomes (C) were quantified by counting within three 35×25(µm^2^) unit areas per individual worm, with a total of 20 worms analyzed (n=20 animals). Scale bars, 10µm. **(D)** Immunoblots and quantification showing cleavage of CHERRY from NUC-1::CHERRY expressed in the epidermis of wild-type (WT), *nsif-1(ok1337)*, ASJ neuron *nsif-1* knock down and ASI neuron *nsif-1* knock down animals (Adult) under E. coli-HT115 feeding condition. N=3 independent replicates. **(E)** Immunoblots and quantification showing cleavage of CHERRY from NUC-1::CHERRY expressed in the epidermis of wild-type (WT), *nsif-1(ok1337)*, ASJ neuron *nsif-1* knockdown strain and ASI neuron *nsif-1* knockdown strain in young adults animals after 24h PA14 exposure (infection initiated at L4 stage). N=3 independent replicates. **(F-G)** Merged fluorescence images (F) and quantification (G) of the epidermis in WT and *nsif-1(ok1337)* expressing NUC-1::sfGFP::CHERRY at 1, 3, 6 and 12 h post-Heat shock at L4 stages under OP50 feeding condition. Pearson’s correlation coefficient for GFP and CHERRY was shown in (G). n=18 animals. Scale bars, 5µm. **(H)** Merged fluorescence images and quantification of the epidermis in WT and *nsif-1(ok1337)* expressing NUC-1::sfGFP::CHERRY at 12 hours post-heat shock (HS) (24h PA14 exposure from L4 stage animals). Pearson’s correlation coefficient for GFP and CHERRY was shown in (F). n=18 animals. Scale bars, 5µm. **(I)** Survival analysis of wild-type(WT), *nsif-1(ok1337), cup-5(bp510)* and *nsif-1(ok1337); cup-5(bp510)* mutant in the PA14 slow-killing assay. PA14 survival assays were performed in three independent replicates, detailed statistical data are presented in source data. For all quantification, data are presented as mean ± SEM. Statistical comparisons were performed using one-way ANOVA followed by Tukey’s multiple comparisons test ( B, C, D, E), unpaired two-tailed Mann–Whitney U-test (G, H) and Log rank (Mantel-Cox) test (I). *Indicates comparison with the wild-type, ^ indicates comparison with the *nsif-1(ok1337)* mutant; */^ p< 0.05, **/^^ p< 0.01, ***/^^^ p< 0.001; ns., not significant. All experiments were performed independently at least three times. The data underlying this figure can be found in **S1 Data**.

Using a NUC-1::sfGFP::CHERRY marker to monitor lysosomal maturation, we observed that in *nsif-1(ok1337)* mutant, lysosomes matured within 6 hours after heat shock, whereas wild-type lysosomes had not yet fully matured, as evidenced by a higher degree of colocalization between GFP and CHERRY (Fig 5F-5G) under OP50 feeding condition. Moreover, we observed lysosomal maturation in NUC-1::sfGFP::CHERRY, infected them with PA14 for 12 hours, subjected them to a heat shock at 34°C for 30 minutes, followed by a 20°C recovery and continued infection for an additional 12 hours. The results revealed that GFP fluorescence was quenched in the *nsif-1* mutant but not wild-type worms after PA14 infection (Fig. 5H). Together, these data indicate that *nsif-1* mutation promotes lysosomal maturation, increases acidification and degradation activity.

Loss of the *C. elegans* lysosomal Ca^²^ channel CUP-5 (homologous to human TRPML1) is known to disrupt lysosomal biogenesis and transport, leading to impaired degradation of endocytic and autophagic cargo[51]. When we introduced a *cup-5* mutation into the *nsif-1* deletion background to disrupt lysosomal function, the enhanced resistance to PA14 infection observed in *nsif-1* mutants was significantly reduced (Fig 5I). In summary, NSIF-1 acts as a negative regulator of lysosomal function, thereby increasing *C. elegans* susceptibility to PA14 infection.

### Neuronal secreted NSIF-1 regulates epidermal lysosomes

NSIF-1 is secreted by neurons and taken up by the epidermis under PA14 infection. To test whether neuronally secreted NSIF-1 specifically regulates epidermal epidermal lysosomal and cuticular function, we expressed NSIF-1 under muscle-specific (*myo-3*) or intestine-specific (*vha-6*) promoters in *nsif-1(ok1337)* mutants. Neither construct reversed the protective cuticle (DPY-7::sfGFP) and lysosomal phenotypes observed in the mutant during PA14 infection, ruling out these tissues as primary targets (S16A-S16B Fig). In contrast, hypodermal expression (*col-12*) of either full-length NSIF-1::GFP or signal peptide-deleted (ΔSP) NSIF-1::GFP abolished these protective benefits and even disrupted lysosomes and cuticle organization under non-pathogenic (OP50) conditions (S16C-S16D Fig). Importantly, ΔSP NSIF-1 was as effective as the full-length protein when expressed locally, confirming that NSIF-1 acts within hypodermal cells. Together, these data indicate that neuronally secreted NSIF-1 functions in the hypodermis to impair lysosomal health and cuticle integrity.

To test whether ASI/ASJ-derived NSIF-1 influences epidermal lysosomal function, we generated ASI/ASJ-specific RNAi strains *qxIs383*;*sid-1*(*qt9*);*ylfEx432*(P*_ASI_*SID-1::3×FLAG;P*_ASI_*NSIF-1::GFP) and *qxIs383*;*sid-1*(*qt9*);*ylfEx433*(P*_ASJ_*SID-1::3×FLAG;P*_ASJ_*NSIF-1::GFP) to knock down *nsif-1* exclusively in these two neurons. Comparing animals fed *nsif-1* RNAi versus the HT115 empty-vector control, NSIF-1::GFP fluorescence in ASI/ASJ was markedly and significantly reduced upon *nsif-1* RNAi, confirming efficient neuron-specific knockdown (Fig 6A-6B). Under PA14 infection, worms with ASI/ASJ-specific *nsif-1* knockdown maintained healthier lysosomal morphology, similar to that observed in *nsif-1(ok1337)* mutants (Fig 6A-6B). Direct measurement of NUC-1::CHERRY cleavage efficiency further confirmed that lysosomal proteolytic activity was significantly higher in the ASI/ASJ-specific *nsif-1* knockdown animals—both under PA14 infection and normal feeding—relative to wild-type worms (Fig 5D-5E).

**Figure 6.**
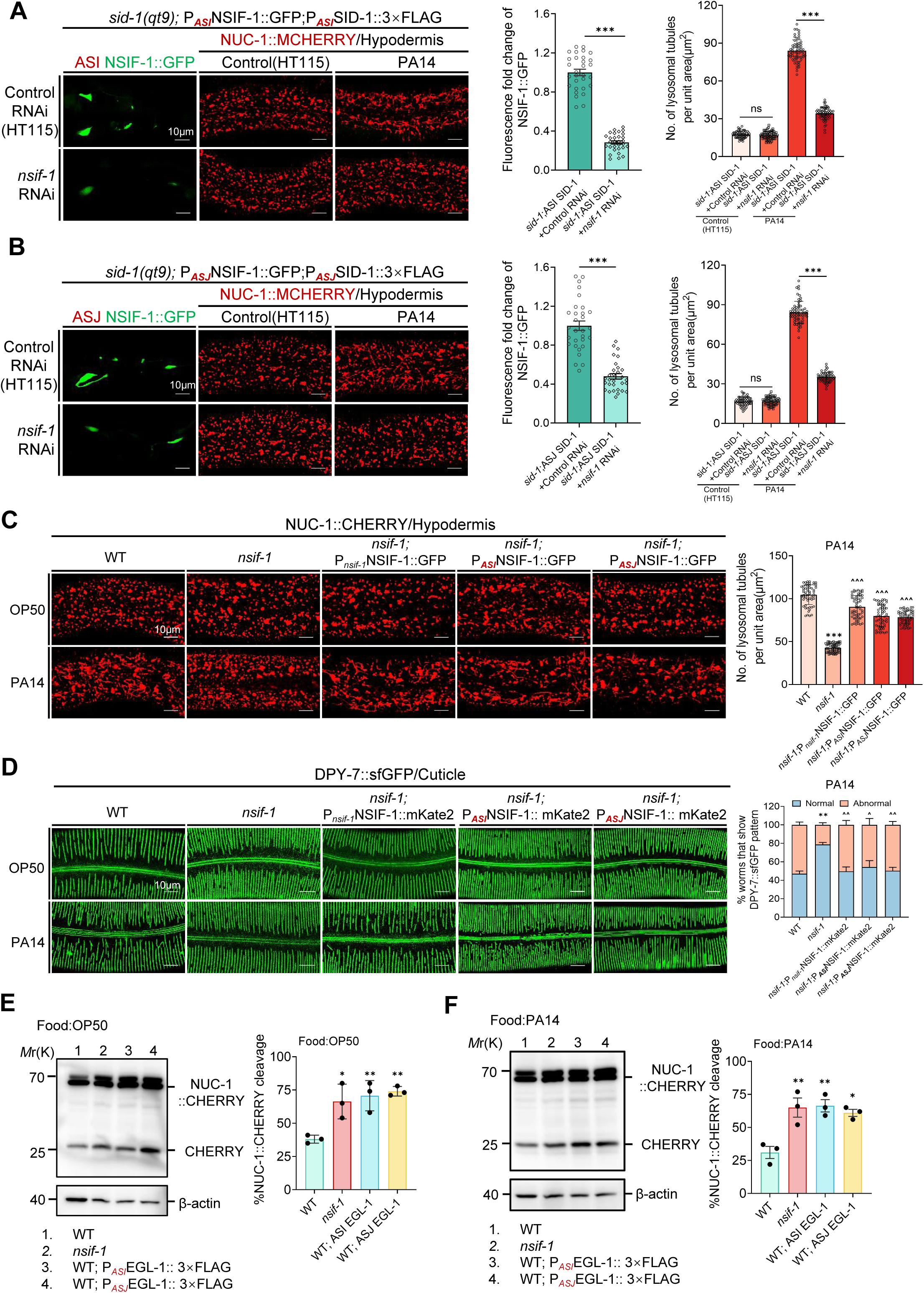
Neuronal secreted NSIF-1 regulates epidermal lysosomes homeostasis. **(A)** Confocal microscopy images and quantification of ASI neuron-specific *nsif-1* knockdown efficiency and lysosomal morphology. Compared with Control RNAi, ASI neuron-specific *nsif-1* knockdown significantly reduced NSIF-1::GFP fluorescence intensity in ASI neurons and decreased the number of tubular lysosomes following 24-h PA14 exposure (infection initiated at the L4 stage). NSIF-1::GFP fluorescence intensity was quantified in 30 worms. Tubular lysosomes were quantified by counting three 35×25µm² regions per worm (n=20 worms). Scale bars, 10µm. **(B)** Confocal microscopy images and quantification of ASJ neuron-specific *nsif-1* knockdown efficiency and lysosomal morphology. Compared with Control RNAi, ASI neuron-specific *nsif-1* knockdown significantly reduced NSIF-1::GFP fluorescence intensity in ASI neurons and decreased the number of tubular lysosomes following 24-h PA14 exposure (infection initiated at the L4 stage). NSIF-1::GFP fluorescence intensity was quantified in 30 worms. Tubular lysosomes were quantified by counting three 35×25µm² regions per worm (n=20 worms). Scale bars, 10µm. **(C)** Confocal fluorescence imaging and quantitative analysis of NUC-1::CHERRY marked lysosome in wild-type(WT), *nsfi-1(ok1337)*, *nsif-1(ok1337)*;P*_nsif-1_*NSIF-1::GFP, *nsif-1(ok1337)*;P*_ASI_*NSIF-1::GFP, *nsif-1(ok1337)*;P*_ASJ_*NSIF-1::GFP animals following 24-h PA14 exposure (infection initiated at the L4 stage). Quantification of tubular lysosomes were quantified by counting three 35 × 25 µm² regions per worm (n =20 worms). Scale bars: 10 µm. **(D)** Confocal fluorescence imaging and quantitative analysis of DPY-7::sfGFP marked cuticle in wild-type (WT), *nsfi-1(ok1337)*, *nsif-1(ok1337)*;P*_nsif-1_*NSIF-1::mKate2, *nsif-1(ok1337)*;P*_ASI_*NSIF-1::mKate2, *nsif-1(ok1337)*;P*_ASJ_*NSIF-1::mKate2 animals following 24-h PA14 exposure (infection initiated at the L4 stage). Quantification of abnormal DPY-7 pattern performed on 300 worms (n=300 animals). Scale bars: 10 µm. **(E)** Immunoblots and quantification showing cleavage of CHERRY from NUC-1::CHERRY expressed in the epidermis of wild-type (WT), *nsif-1(ok1337)*, and ASJ and ASI neurons ablate animals under OP50 feeding conditions. N=3 independent replicates. **(F)** Immunoblots and quantification showing cleavage of CHERRY from NUC-1::CHERRY expressed in the epidermis of wild-type (WT), *nsif-1(ok1337)*, and ASJ and ASI neurons ablate animals under PA14 feeding conditions. N=3 independent replicates. For all quantification, data are presented as mean ± SEM. Statistical comparisons were performed using one-way ANOVA followed by Tukey’s multiple comparisons test (C, D, E, F) and Mann–Whitney U-test (A, B). *Indicates comparison with the wild-type, ^ indicates comparison with the *nsif-1(ok1337)* mutant; **/^^ p < 0.01, ***/^^^ p< 0.001; ns., not significant. All experiments were performed independently at least three times. The data underlying this figure can be found in **S1 Data**.

To ensure that our findings reflect the physiological role of NSIF-1, we performed comprehensive rescue experiments in the *nsif-1(ok1337)* mutant background. We expressed the NSIF-1 coding sequence under its native promoter, as well as under ASJ- (*trx-1*) or ASI-(*str-3*) specific promoters. Under PA14 infection, both the native and neuron-specific expression restored wild-type lysosomal morphology and rescued cuticle organization in the mutant (Fig 6C-6D). These results support our model that neuronal secreted NSIF-1 acts non-cell-autonomously to regulate lysosomal function in peripheral tissues. However, we also examined versions lacking the signal peptide (ΔSP). Unexpectedly, ΔSP NSIF 1 expressed from ASI- or ASJ-specific promoters still rescued lysosomal morphology and cuticle organization in the mutant (S17A-S17B Fig). One plausible explanation is that the elevated level of ΔSP NSIF 1 in ASI or ASJ may trigger other unknown signaling pathways that restore wild type lysosomal morphology and rescue cuticle organization in the mutant.

To further validate the neuronal origin of NSIF-1 in regulation epidermal lysosomes, we genetically ablated ASJ/ASI neurons using the cell death activator EGL-1 [66]. To confirm efficacy of EGL-1-mediated ASI/ASJ ablation, we co-expressed cell-death activator EGL-1 with ASI/ASJ-specific NSIF-1::GFP in the *qxIs383* background. Compared with sibling controls lacking EGL-1, animals carrying the EGL-1 transgene showed a marked reduction in NSIF-1::GFP fluorescence in the targeted neurons, consistent with successful cell ablation (S18A Fig). Ablation of ASJ/ASI neurons significantly reduced NSIF-1 secretion (S18B Fig), as measured by GFP-binding nanobody (GBP) capture system, confirming that these neurons are the primary source of NSIF-1.

Moreover, pathogen-induced expansion of tubular lysosomes was significantly suppressed in both *nsif-1(ok1337)* mutants and animals subjected to ASI/ASJ neuronal ablation (S18A, S19A-S19B Fig). Furthermore, assessment of lysosomal proteolytic activity revealed consistent trends: cleaved CHERRY levels—a direct indicator of lysosomal degradation efficiency—were elevated in *nsif-1(ok1337)* mutants and neuron-ablated animals relative to wild-type controls under both standard (OP50) and pathogenic (PA14) conditions (Fig 6E-6F).

In summary, our data indicate that ASI/ASJ neuronal secreted NSIF-1 acts non-cell-autonomously to impair lysosomal health and cuticle integrity in the hypodermis.

### NSIF-1 impairs lysosomal function via inhibiting ELT-3

We have demonstrated that PA14-induced impairment of lysosomal function requires NSIF-1, particularly its neuronal secretion, which depends on its signal peptide (Fig 4). To directly test whether NSIF-1 is sufficient to impair epidermal lysosomal function, we expressed either full-length NSIF-1 (FL NSIF-1) or a signal peptide-deleted version (ΔSP NSIF-1) under the native *Pnsif-1* promoter or the epidermis-specific *col-12* promoter in wild-type animals. Overexpression of either construct from the native *Pnsif-1* promoter in wild-type animals produced no detectable change in hypodermal lysosomal morphology (S20A Fig) or cuticle organization (S20B Fig) compared to wild-type control under OP50 feeding condition. In contrast, when the same constructs were driven by the hypodermal *col-12* promoter, both FL NSIF-1 and ΔSP NSIF-1 markedly disrupted lysosomal morphology (S20A Fig) and cuticle organization (S20B Fig), phenocopying the effects of PA14 infection (Fig 1A, 2A). These results indicate two key points: (i) overexpression of NSIF-1 under non-infection conditions from its native promoter is well tolerated, consistent with our model that it is the PA14-induced secretion of NSIF-1 that drives pathology; and (ii) forced expression of NSIF-1 directly in the hypodermis is sufficient to recapitulate the lysosomal and cuticle defects, supporting the conclusion that the hypodermis is the site of NSIF-1 action.

Using cNLS Mapper (https://nls-mapper.iab.keio.ac.jp/), we identified a putative NLS sequence within NSIF-1 with a score of 3.3 (S21 Fig). According to the cNLS Mapper scoring system established by Kosugi et al. (2009) [67], scores of 3- 5 indicate proteins that localize to both the nucleus and the cytoplasm. Moreover, we found that ΔSP NSIF-1 localizes to the nucleus (S22A Fig). This observation led us to explore its potential interaction with ELT-3, a transcription factor known to enhance lysosomal function [18] and regulate collagen organization [8]. Consistent with this hypothesis, we observed strong nuclear co-localization between ΔSP NSIF-1 and ELT-3 (S22A Fig). Furthermore, transcription factor enrichment analysis through WormExp (https://wormexp.zoologie.uni-kiel.de/wormexp/) [68] revealed approximately 200 upregulated genes in *nsif-1* mutants that are putative ELT-3 targets (S22B Fig; S3 Table). This suggests that loss of NSIF-1 may enhance ELT-3 activity, potentially driving lysosomal functional enhancement. To further test this hypothesis, we examined ELT-3 localization in *nsif-1* mutant animals. We found that loss of *nsif-1* resulted in significantly enhanced nuclear expression of ELT-3 (Fig 7A). Moreover, we performed the qRT-PCR on synchronized L4 + 24 h wild-type and *nsif-1(ok1337)* animals and find that endogenous *elt-3* mRNA is significantly upregulated in *nsif-1(ok1337)* compared with wild type (S22C Fig). These data indicate that NSIF-1 normally suppresses ELT-3 nuclear expression.

**Figure 7.**
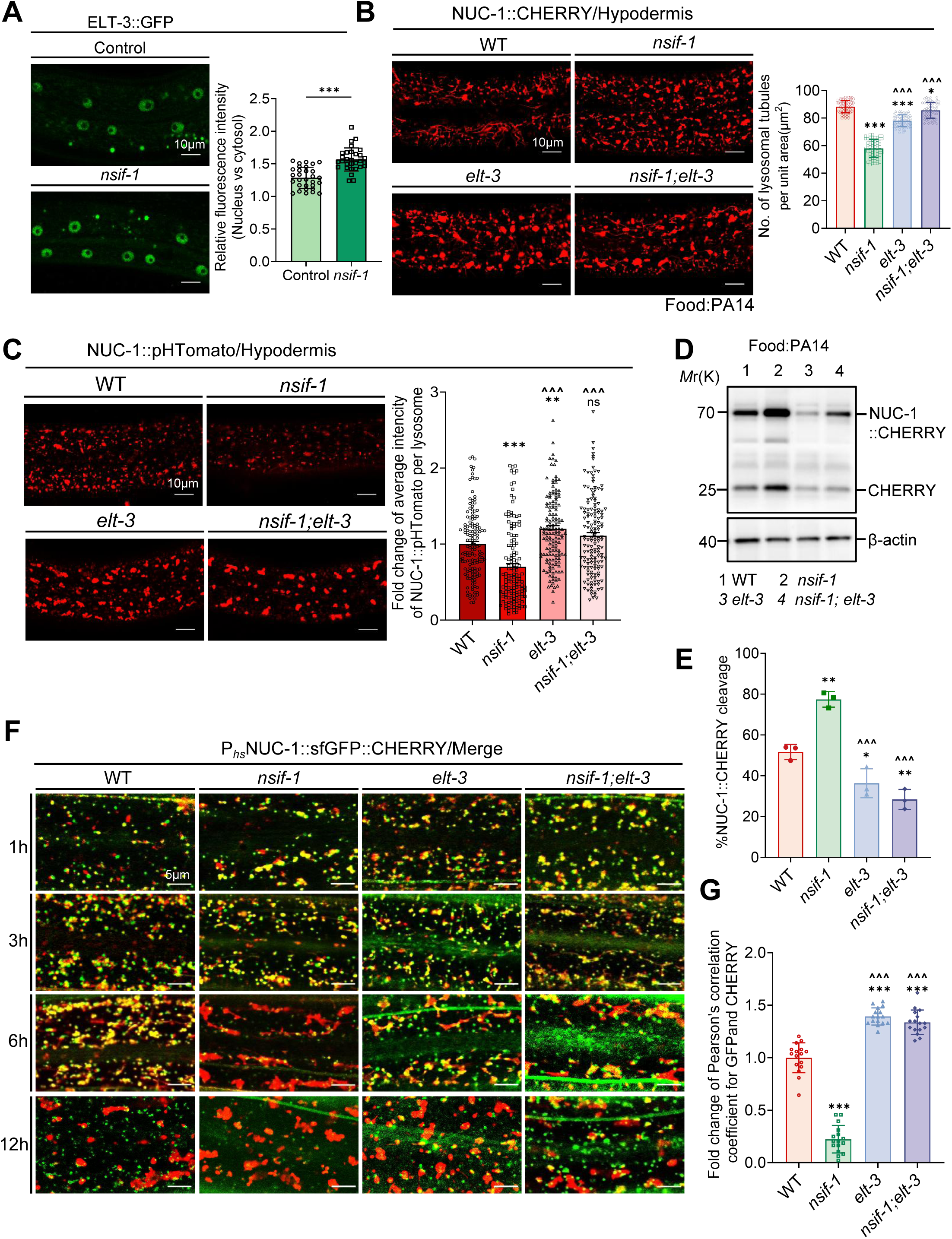
NSIF-1 impairs lysosomal function via inhibiting ELT-3. **(A)** Confocal fluorescence images and quantification of epidermal ELT-3::GFP in wild-type (WT), *nsif-1(ok1337)*. n=30 animals. Scale bars, 10µm. **(B)** Confocal fluorescence images of the hypodermis in young adults wild-type (WT), *nsif-1(ok1337)*, *elt-3(vp1), nsif-1(ok1337);elt-3(vp1)* expressing NUC-1::CHERRY after 24h PA14 exposure (infection initiated at L4 stage). The number of tubular lysosomes was quantified by counting within three 35×25(µm^2^) unit areas per individual worm, with a total of 20 worms analyzed (n=20 animals). Scale bars, 10µm. **(C)** Confocal fluorescence imaging and quantitative analysis were performed on the hypodermis of wild-type (WT), *nsif-1(ok1337)*, *elt-3(vp1), nsif-1(ok1337);elt-3(vp1)* expressing NUC-1::pHTomato under the control of a heat-shock promoter following 24-hours PA14 exposure (infection initiated at L4 stage). Fluorescence quantification was performed on 150 lysosomes from 20 worms (n=20 animals). Scale bars, 10µm. **(D-E)** Immunoblots (D) and quantification (E) showing cleavage of CHERRY from NUC-1::CHERRY expressed in the epidermis of wild-type(WT), *nsif-1(ok1337)*, *elt-3(vp1), nsif-1(ok1337);elt-3(vp1)* after 24h PA14 exposure (infection initiated at L4 stage). N=3 replicates in (E). **(F-G)** Merged fluorescence images (F) and quantification (G) of the epidermis in WT, *nsif-1(ok1337)*, *elt-3(vp1) and nsif-1(ok1337);elt-3(vp1)* expressing NUC-1::sfGFP:: CHERRY at 1, 3, 6 and 12 h post-heat shock (24h PA14 exposure from L4 stage animals). Pearson’s correlation coefficient for GFP and CHERRY was shown in (G). n=16 animals. Scale bars, 5µm. For all quantification, data are presented as mean ± SEM. Statistical comparisons were performed using one-way ANOVA followed by Tukey’s multiple comparisons test (B, C, E, G) and Mann–Whitney U-test (A). *Indicates comparison with the wild-type, ^ indicates comparison with the *nsif-1(ok1337)* mutant; */^ p < 0.05, **/^^ p < 0.01, ***/^^^ p< 0.001; ns., not significant. All experiments were performed independently at least three times. The data underlying this figure can be found in **S1 Data**.

If NSIF-1 functions through ELT-3 inhibition, then the lysosomal enhancement observed in *nsif-1* mutants should depend on ELT-3 activity. To validate this hypothesis, we performed a series of genetic epistasis experiments using *nsif-1;elt-3* double mutants. Firstly, we observed lysosomal morphology. The reduction in tubular lysosomes observed in *nsif-1* mutants under PA14 infection was suppressed in the *nsif-1;elt-3* double mutants, where tubular lysosomes, vesicular lysosomes were restored to levels comparable to *elt-3* single mutants (Figs 7B, S22D). Additionally, the lysosomal diameter in *nsif-1*; *elt-3* double mutants exhibited an enlarged lysosome diameter similar to that of *elt-3* single mutants (Figs 7B, S22E). This indicates that the morphological changes caused by *nsif-1* mutation are dependent on ELT-3.

Secondly, we measured the lysosomal acidification. In *nsif-1* mutants, lysosomes exhibited enhanced acidification, as indicated by reduced NUC-1::pHTomato fluorescence intensity (Fig 7C). However, in *nsif-1;elt-3* double mutants, lysosomal acidification was diminished, resembling the phenotype of *elt-3* mutants (Fig 7C). This demonstrates that the enhanced acidification caused by *nsif-1* loss is dependent on ELT-3. Thirdly, lysosomal degradation activity, measured by NUC-1::CHERRY cleavage, was elevated in *nsif-1* mutants but returned to *elt-3* single mutant levels in the *nsif-1;elt-3* double mutants under both PA14 infection and normal feeding conditions (Figs 7D-7E, S22F). This further supports that ELT-3 is necessary for the increased degradation activity observed in the absence of *nsif-1*.

Finally, we monitored lysosomal maturation following heat shock and found that lysosomes in *nsif-1* mutants matured within 6 hours, whereas lysosomes in *nsif-1; elt-3* double mutants displayed delayed maturation, with persistent colocalization between GFP and CHERRY signals (Fig 7F-7G). This indicates that accelerated lysosomal maturation in *nsif-1* mutants also requires ELT-3.

Taken together, these findings demonstrate that NSIF-1 impairs lysosomal function by inhibiting ELT-3. Loss of *nsif-1* leads to increased nuclear localization of ELT-3, which subsequently enhances lysosomal acidification, degradation capacity, and maturation. Therefore, ELT-3 functions downstream of NSIF-1 and is both necessary and sufficient for mediating the lysosomal improvements observed in *nsif-1* mutants.

### NSIF-1 damages the cuticle by inhibiting ELT-3

Our RNA-seq analysis comparing wild-type N2 and *nsif-1(ok1337)* mutants, revealed that *nsif-1* mutants exhibit elevated expression of collagen-related genes (Fig 8A; S3 Table). This upregulation likely contributes to the observed densification of the cuticle structure in *nsif-1(ok1337)* mutants (Figs 3D, 8B).

**Figure 8.**
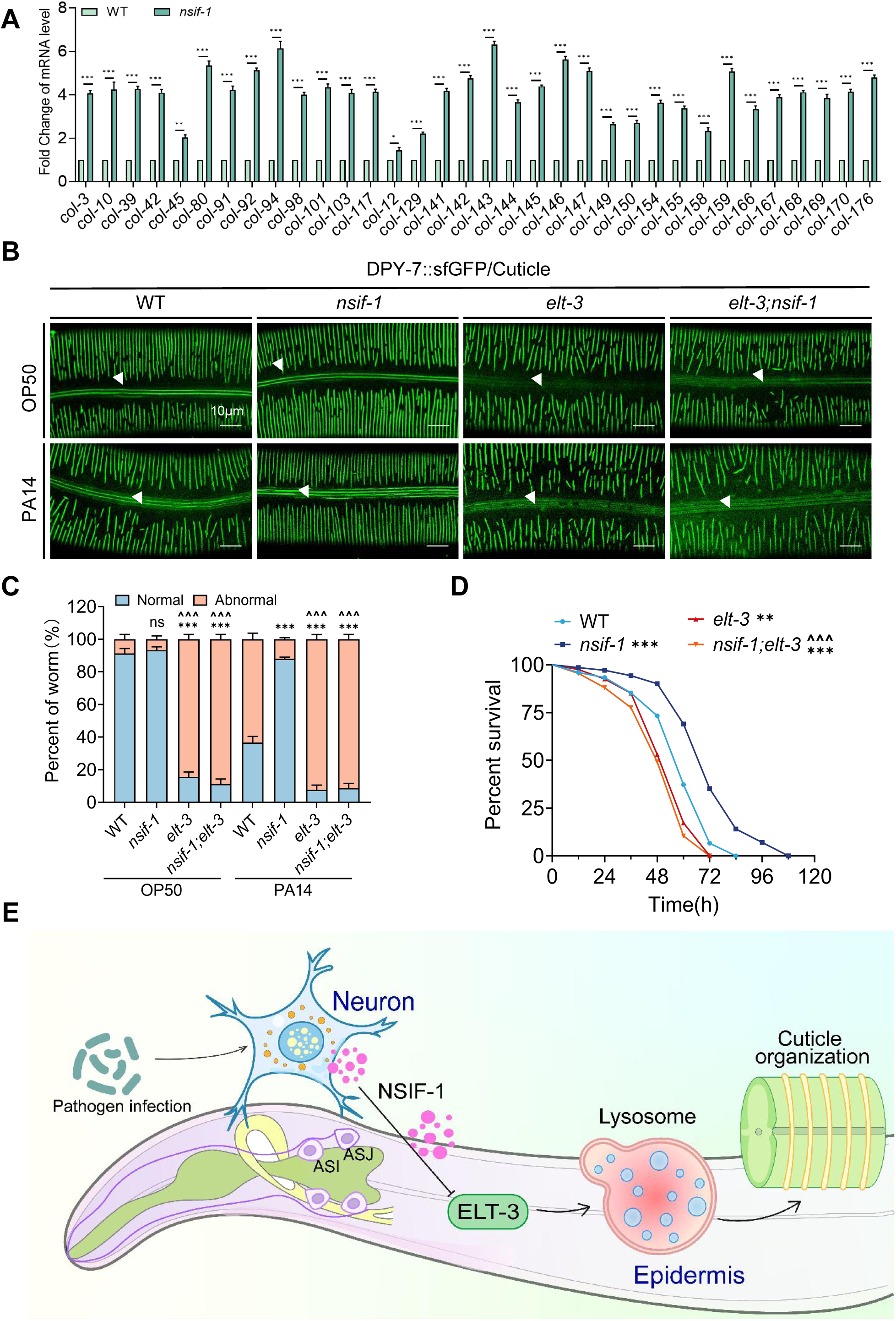
NSIF-1 disrupts collagen organization through ELT-3 which facilitates PA14 infection. **(A)** Upregulated collagen genes in *nsif-1(ok1337)* which was extracted from the RNA-seq data; **(B-C)** Confocal fluorescence images (B) of DPY-7::sfGFP in wild-type (WT), *nsif-1(ok1337)elt-3(vp1)* and *nsif-1(ok1337);elt-3(gk121)* following 24-hours PA14 expo-sure (infection initiated at L4 stage). Quantification of abnormal DPY-7::sfGFP pattern (C) is calculated in 90 Worms (n=90 animals). The fluorescence signal associated with alae structures (Arrowhead) constitutes autofluorescence in (B). **(D)** Survival analysis of wild-type (WT), *nsif-1(ok1337)*, *elt-3(gk121), nsif-1(ok1337); elt-3(gk121)* mutant in the PA14 slow-killing assay. PA14 survival assays were performed in three independent replicates, detailed statistical data are presented in source data. **(E)** Schematic model of a neuro-epidermal signaling axis in which *Pseudomonas aeruginosa* PA14 exploits neuronal NSIF-1 to disrupt epidermal lysosomal function, compromise collagen integrity, and suppress host immunity in *C. elegans*. NSIF-1 acts as a neuronal secreted factor that mediates PA14-induced pathogenesis, inhibiting epidermal lysosomal activity via the transcription factor ELT-3. For all quantification, data are presented as mean ± SEM. Statistical comparisons were performed using one-way ANOVA followed by Tukey’s multiple comparisons test (A, C) and Log rank (Mantel-Cox) test (D). *Indicates comparison with the wild-type, ^ indicates comparison with the *nsif-1(ok1337)* mutant; **/^^p < 0.01, ***/^^^p < 0.001; ns., not significant. All experiments were performed independently at least three times. The data underlying this figure can be found in **S1 Data**.

Given that (i) ELT-3, a transcription factor, is known to activate lysosomal function and regulate collagen organization, and (ii) *nsif-1* mutation enhances the nuclear localization of ELT-3, we hypothesized that the restoration of cuticle collagen organization observed in *nsif-1* mutants under PA14 infection depends on ELT-3.

Under normal OP50 conditions, *elt-3* mutants display fragmented and disorganized annular structures in the cuticle, and *nsif-1; elt-3* double mutants exhibit similar cuticle features to *elt-3* single mutants (Fig 8B-8C). This observation indicates that ELT-3 functions downstream of NSIF-1 in maintaining proper collagen organization. During PA14 infection, the disrupted collagen organization seen in wild-type worms is significantly alleviated in *nsif-1* mutants. However, in *nsif-1; elt-3* double mutants, the dense furrows seen in *nsif-1* mutants is lost, replaced by a fragmented furrows pattern similar to that observed in *elt-3* mutants (Fig 8B–8C). This finding demonstrates that the maintenance of cuticle collagen organization in *nsif-1* mutants under PA14 infection is dependent on ELT-3. Moreover, the loss of ELT-3 renders *nsif-1* mutants as susceptible to PA14 infection as *elt-3* mutants (Fig 8D). Collectively, our data indicate that PA14-stimulated NSIF-1 impairs epidermal cuticle collagen by inhibiting ELT-3, thereby facilitating PA14 infection.

In summary, we identify NSIF-1 as a neurosecretory inhibitory factor released in response to PA14 infection. Upon secretion, NSIF-1 is taken up by epidermis, where NSIF-1 disrupts lysosomal function by inhibiting ELT-3 activity, thereby impairing cuticle collagen organization (Fig 8E).

## Discussion

This study uncovers a novel neuro-epidermal signaling axis through which the pathogen *Pseudomonas aeruginosa* PA14 hijacks neuronal communication to impair lysosomal function, disrupt collagen organization, and suppress host immunity in *C. elegans*. We identify NSIF-1 (TAG-293) as a neuronal secreted factor that acts as a critical mediator of PA14-induced pathogenesis, inhibiting lysosomal activity in the epidermis via the transcription factor ELT-3 (Fig 8E). These findings expand our understanding of how pathogens exploit host neuronal signaling to compromise barrier defenses and immune responses.

### Neuronal Regulation of Epidermal Lysosomal Function

Lysosomes are central to ECM remodeling and barrier maintenance, with their activity tightly regulated during development and stress responses. Previous studies in *C. elegans* demonstrated that lysosomal activation is essential for cuticle collagen turnover during molting [18]. Our work extends these findings by showing that pathogens like PA14 disrupt this process through neuronal signaling. NSIF-1, secreted from ASI/ASJ neurons in response to infection, inhibits epidermal lysosomal acidification, degradation, and maturation. This mechanism mirrors strategies employed by intracellular pathogens such as *Mycobacterium tuberculosis* and *Salmonella*, which impair lysosomal function to evade host defenses [47–50]. However, our study uniquely highlights neurons as initiators of epidermal lysosomal dysfunction during infection, a pathway not previously described in metazoans.

In our model, we propose that neuronal NSIF-1 is secreted and subsequently taken up by hypodermal cells. Because the signal peptide is co-translationally removed during biogenesis in the neuron, the internalized protein in the hypodermal cell is already the mature (SP-cleaved) form, yet retains its intact internal NLS. Once inside the hypodermal cell, this mature protein may escape from endosomal compartments into the cytosol and then translocate to the nucleus via NLS-mediated import. However, the precise molecular mechanism by which NSIF-1 is diverted from the secretory pathway to the nucleus remains to be determined.

The nuclear co-localization of ΔSP NSIF-1 with ELT-3 and the restoration of lysosomal defects in *nsif-1;elt-3* double mutants reveal that NSIF-1 acts by antagonizing ELT-3, a transcription factor known to enhance lysosomal activity and collagen organization. This interaction aligns with reports that ELT-3 regulates stress-responsive genes in the epidermis [8, 69], but our discovery of its suppression by a neuronal derived factor adds a new layer of complexity to neuro-epidermal crosstalk. Notably, the dual role of NSIF-1 in both lysosomal and collagen regulation suggests that pathogens target centralized regulatory nodes to maximize tissue damage—a strategy analogous to bacterial toxins that disrupt multiple host pathways simultaneously.

### Collagen Remodeling Regulated by Lysosomes as a Host–Pathogen Battleground

Collagen disruption is a common feature in infections ranging from *Staphylococcus aureus*-induced skin lesions to *Leishmania*-associated cutaneous ulcers [70–73]. In *C. elegans*, PA14-induced cuticle damage activates epidermal immunity (Fig. 1), consistent with previous findings that defects in epidermal or cuticle collagen trigger compensatory immune responses [38, 74]. While earlier studies have shown that disruption of the attachments between the cuticle, epidermis, and basement membrane activates lysosomal activity in the epidermis [18], our work reveals a paradox: although PA14 damages collagen to breach the host barrier, the nematode’s own neuronal response—mediated by NSIF-1—further exacerbates this damage by suppressing lysosomal-dependent repair mechanisms.

Under normal conditions, epidermal damage activates lysosomal pathways to degrade damaged ECM components and promote the synthesis of new cuticle, thereby maintaining tissue homeostasis [18]. This process is analogous to the lysosomal activation observed during the molting period, where new collagen is synthesized. However, upon PA14 infection, the pathogen exploits host neuronal signals by inducing the secretion of NSIF-1 to suppress epidermal lysosomal activity. This suppression hinders the normal repair and remodeling of the ECM, thus facilitating pathogen invasion. Our findings suggest that PA14 hijacks host factors regulating extracellular matrix (ECM) and collagen integrity as a virulence strategy, disrupting tissue repair to facilitate infection. Therefore, our findings underscore that the modulation of lysosomal function represents a critical battleground in host-pathogen interactions, with potential implications for therapeutic interventions in chronic infectious and wound-healing disorders.

### Implications for Host-Directed Therapies

Our findings suggest that targeting neuro-immune signaling could mitigate pathogen-induced tissue damage. For instance, blocking NSIF-1 secretion or enhancing ELT-3 activity might restore lysosomal function and collagen integrity during infection. This approach aligns with emerging therapies for fibrosis and chronic infections, where lysosomal reactivation (e.g., via TRPML1 agonists) shows promise [75]. Although the role of neuronal secreted NSIF-1 in disrupting epidermal lysosomal function is specific to *C. elegans*, the evolutionary conservation of ELT-3/GATA transcription factors across species raises the possibility that similar neuro-epidermal pathways operate in mammals, offering translational relevance.

In summary, we delineate a pathway where PA14 exploits neuronal signaling to impair lysosomal function and disrupt collagen. By linking neurosecretion, lysosomal regulation, and ECM dynamics, this work advances our understanding of host-pathogen interactions and highlights neuronal signaling as a vulnerable target for therapeutic intervention.

### Limitations of the Study

While our findings reveal a neuro-epidermal pathway linking neuronal signaling, lysosomal dysfunction, and collagen disorganization, several limitations should be acknowledged.

First, although we demonstrate that NSIF-1 disrupts lysosomal function via ELT-3, the precise mechanism by which NSIF-1 inhibits ELT-3 nuclear localization remains unresolved. Similarly, the intracellular itinerary of NSIF-1—specifically, how it escapes the endosomal compartment and translocates to the nucleus—is still unknown. Identifying the receptor(s) responsible for NSIF-1 uptake and the machinery governing endosomal escape will be important directions for future research. Furthermore, whether signal peptide (SP) cleavage efficiency or alternative trafficking routes contribute to NSIF-1 nuclear localization *in vivo* during infection warrants further investigation.

Second, our study does not fully distinguish whether the observed lysosomal dysfunction primarily affects collagen degradation, synthesis, or recycling. Addressing these mechanistic gaps could provide deeper insights into how pathogens exploit neuro-epidermal extracellular matrix (ECM) pathways and may inform potential therapeutic strategies for combating chronic infections and promoting tissue repair.

## Materials and methods

### *C. elegans* strains and maintenance

1. The following strain/alleles was obtained from *Caenorhabditis Genetics Center* (CGC) and SunyBiotech (Fuzhou, China): N2 Bristol (wild type strain); VC798: *tag-293/nsif-1(ok1337)*; KU25: *pmk-1(km25)*; VC143: *elt-3(gk121)*; IG274: *frIs7* [P*_nlp-29_*GFP+P*_col-12_*DsRed]; AU78: *agIs219* P*_sysm-1_*GFP*:: unc-54* 3’UTR + P*_ttx-3_*GFP*+unc54* 3’UTR]; OP559: *unc-119(tm4063);wgIs559 elt-3:*:TY1::EGFP::3×FLAG *+ unc-119(+)*] DV3327: *pmk-1(re170)* pmk-1::mNeonGreen::3xFlag]; DR96: *unc-76(e911)*; SD1809: *elt-3(vp1)*; HZ108: *cup-5(bp510)*; JDW389: *bli-1(wrd84) bli-1*::linker::mNeonGreen::3xFLAG(internal)::linker]; JDW461: *noah-2(wrd120)* [*noah-2*::mNeonGreen::3xFLAG]; JDW655: *cut-2(wrd233) cut-2*::mNeonGreen::3xFLAG]. BHX9508: *nsif-1(syb9508) nsif-1*::mNeonGreen::3xFLAG knock-in], (This strain was generated by SunyBiotech.
2. The following strain/alleles was obtained from Dr. Xiaochen Wang lab or Dr. Chonglin Yang lab: XW5399: *qxIs257* P*_ced-1_*NUC-1::CHERRY] [76]; XW17096: *qxIs383* P*_hyp-7_*NUC-1::CHERRY] [18, 51]; XW13734: *qxIs612* P*_hsp_*NUC-1::sfGFP::CHERRY] [18, 51]; XW18042: *qxIs722* P*_dpy-7_*DPY-7::sfGFP, single-copy insertion] [18]; XW19180: *qxIs750* [P*_hsp_*NUC-1::pHTomato] [18, 51]; XW17147: *cup-5(bp510);qxIs383* P*_hyp-7_*NUC-1::CHERRY]
3. The following strains were constructed by this study: YNU6: *unc-76(e911);tag-293/nsif-1(ok1337)*; YNU210: *elt-3(gk121);nsif-1(ok1337*) YNU467: *nsif-1(ok1337);qxIs383* P*_hyp-7_*NUC-1::CHERRY]; YNU470: *nsif-1(ok1337); qxIs612* P*_hsp_*NUC-1::sfGFP::CHERRY]; YNU471: *nsif-1(ok1337);qxIs750* [P*_hsp_*NUC-1::pHTomato]; YNU473: *nsif-1(ok1337);qxIs722* P*_dpy-7_*DPY-7::sfGFP, single-copy insertion]; YNU474*: nsif-1(ok1337);wgIs559 elt-3:*:TY1::EGFP::3×FLAG *+ unc-119(+)*]; YNU512: *cup-5(bp510);nsif-1(ok1337);qxIs383*; YNU529: *elt-3(vp1); qxIs383* P*_hyp-7_*NUC-1::CHERRY]; YNU504: e*lt-3(gk121); qxIs750* [P*_hsp_*NUC-1::pHTomato]; YNU510: e*lt-3(gk121); qxIs612* P*_hsp_*NUC-1::sfGFP::CHERRY] YNU534: e*lt-3(gk121); qxIs722* P*_dpy-7_*DPY-7::sfGFP, single-copy insertion] YNU569: *nsif-1(ok1337)*;*elt-3(vp1); qxIs383* P*_hyp-7_*NUC-1::CHERRY]; YNU527: *nsif-1(ok1337)*;*elt-3(vp1); qxIs612* P*_hsp_*NUC-1::sfGFP::CHERRY]; YNU541: *nsif-1(ok1337)*;*elt-3(vp1);qxIs750* [P*_hsp_*NUC-1::pHTomato]; YNU514: *nsif-1(ok1337)*;*elt-3(vp1); qxIs722* P*_dpy-7_*DPY-7::sfGFP, single-copy insertion]; YNU18: *unc-76(e911);ylfEx17* P*_nsif-1_*::GFP*; unc-76*(+)]; YNU22: *unc-76(e911);ylfIs1* P*_nsif-1_*NSIF-1::GFP*; unc-76*(+)]; YNU134: *unc-76(e911);ylfIs4* P*_nsif-1_*NSIF-1::3×FLAG::GFP*; unc-76*(+)]; YNU138: *ylfIs8* P*_rgef-1_*NSIF-1::GFP+ P*_odr-1_*RFP]; YNU270: *ylfIs22* P*_vha-6_*NSIF-1::GFP+ P*_odr-1_*RFP]; YNU275: *ylfIs27* P*_myo-3_*NSIF-1::GFP+ P*_odr-1_*RFP]; YNU278: *unc-76(e911);ylfIs30* P*_nsif-1_*ΔSPNSIF-1::GFP+*unc-76*(+)]; YNU468: *ylfEx261* P*_col-12_*GBP::mKate2+*ylfIs8*+*rol-6*(*su1006*)]; YNU469: *ylfEx262* P*_rgef-1_*ΔSPNSIF-1::GFP;P*_col-12_*GBP::mKate2;*rol-6*(*su1006*)]; YNU284: *ylfEx151* P*_sra-6_*mKate2+ P*_nsif-1_*GFP]; YNU285: *ylfEx152* P*_trx-1_*mKate2+ P*_nsif-1_*GFP]; YNU286: *ylfEx153* P*_tph-1_*mKate2+ P*_nsif-1_*GFP]; YNU287: *ylfEx154* P*_ggr-2_*mKate2+ P*_nsif-1_*GFP]; YNU554: *ylfEx281 qxIs257*; P*_col-12_*NSIF-1::GFP+P*_odr-1_*RFP]; YNU555: *ylfEx282 qxIs257*; P*_col-1_*_2_ΔSPNSIF-1::GFP+P*_odr-1_*RFP]; YNU777: *epic-1(ylf93) epic-1*::GFP::3xFLAG knock-in]; YNU738: *nsif-1(ylf82)*; YNU763: *syb9508*;*qxIs383* P*_hyp-7_*NUC-1::CHERRY]; YNU764: *nsif-1(ylf82)*;*qxIs383* P*_hyp-7_*NUC-1::CHERRY]; YNU747: *ylfIs52* P*_col-12_*GBP::mKate2]; YNU765: *ylfIs52* P*_col-12_*GBP::mKate2];*ylfIs8* P*_rgef-1_*NSIF-1::GFP+ P*_odr-1_*RFP]; YNU556: *ylfEx283 qxIs383;nsif-1(ok1337)*; P*_trx-1_*NSIF-1::GFP+P*_odr-1_*RFP]; YNU557: *ylfEx284 qxIs383;nsif-1(ok1337)*; P*_trx-1_*ΔSPNSIF-1+P*_odr-1_*RFP]; YNU558: *ylfEx285 qxIs383;nsif-1(ok1337)*; P*_str-3_*NSIF-1::GFP+P*_odr-1_*RFP]; YNU559: *ylfEx286 qxIs383;nsif-1(ok1337)*; P*_str-3_*ΔSPNSIF-1+P*_odr-1_*RFP]; YNU755: *ylfEx357 qxIs383;*P*_str-3_*EGL-1::3×FLAG + P*_odr-1_*RFP]; YNU756: *ylfEx358 qxIs383;*P*_trx-1_*EGL-1::3×FLAG+ P*_odr-1_*RFP]; YNU757: *ylfEx359 qxIs722;nsif-1(ok1337);*P*_str-3_*NSIF-1::mKate2+ P*_odr-1_*GFP]; YNU758: *ylfEx359 qxIs722;nsif-1(ok1337);*P*_str-3_*ΔSPNSIF-1::mKate2+ P*_odr-1_*GFP]; YNU759: *ylfEx361 qxIs722;nsif-1(ok1337);*P*_trx-1_*NSIF-1::mKate2+ P*_odr-1_*GFP]; YNU760: *ylfEx362 qxIs722;nsif-1(ok1337);*P*_trx-1_*ΔSPNSIF-1::mKate2+ P*_odr-1_*GFP]; YNU761: *qxIs383;nsif-1(ok1337);ylfIs1* P*_nsif-1_*NSIF-1::GFP]; YNU762: *qxIs383;nsif-1(ok1337);ylfIs30* P*_nsif-1_*ΔSPNSIF-1::GFP]; YNU779: *ylfEx363 qxIs722;nsif-1(ok1337);*P*_nsif-1_*NSIF-1::mKate2+ P*_odr-1_*GFP]; YNU786: *ylfEx370 qxIs722;nsif-1(ok1337);*P*_nsif-1_*ΔSPNSIF-1::mKate2+ P*_odr-1_*GFP]; YNU778: *sid-1(qt9);qxIs383* P*_hyp-7_*NUC-1::CHERRY]; YNU780: *ylfEx364* [*sid-1(qt9);qxIs383;* P*_str-3_*SID-1::3×FLAG+P*_odr-1_*RFP]; YNU781: *ylfEx365 sid-1(qt9);qxIs383;*P*_trx-1_*SID-1::3×FLAG+P*_odr-1_*RFP]; YNU782: *ylfEx366 ylfIs52;ylfIs8+*P*_str-3_*EGL-1::3×FLAG+*rol-6*(*su1006*)]; YNU783: *ylfEx367 ylfIs52;ylfIs8+*P*_trx-1_*EGL-1::3×FLAG+*rol-6*(*su1006*)]; YNU794: *nsif-1(ylf82);qxIs722* P*_dpy-7_*DPY-7::sfGFP, single-copy insertion]; YNU811: *nsif-1(ylf82);bli-1(wrd84) bli-1*::linker::mNeonGreen::3xFLAG(internal) ::linker]; YNU813: *nsif-1(ylf82); cut-2(wrd233) cut-2*::mNeonGreen::3xFLAG]; YNU814: *nsif-1(ylf82); epic-1(ylf93) epic-1*::GFP::3xFLAG knock-in]; YNU815: *nsif-1(ok1337);bli-1(wrd84) bli-1*::linker::mNeonGreen::3xFLAG(internal) ::linker]; YNU817: *nsif-1(ok1337); cut-2(wrd233) cut-2*::mNeonGreen::3xFLAG]; YNU818: *nsif-1(ok1337); epic-1(ylf93) epic-1*::GFP::3xFLAG knock-in]; YNU858: *ylfEx411 ylfIs52*;P*_str-3_*NSIF-1::GFP+ *rol-6*(*su1006*)]; YNU859: *ylfEx412 ylfIs52*;P*_str-3_*ΔSPNSIF-1::GFP+ *rol-6*(*su1006*)]; YNU860: *ylfEx413 ylfIs52*;P*_trx-1_*NSIF-1::GFP+ *rol-6*(*su1006*)]; YNU861: *ylfEx414 ylfIs52*;P*_trx-1_*ΔSPNSIF-1::GFP+ *rol-6*(*su1006*)]; YNU862: *ylfEx415 qxIs383;nsif-1(ok1337)*+P*_col-12_*NSIF-1::GFP+P*_odr-1_*RFP]; YNU863: *ylfEx416 qxIs383*;*nsif-1(ok1337)*+P*_col-12_*ΔSPNSIF-1::GFP+P*_odr-1_*RFP]; YNU864: *ylfEx417 qxIs383*+P*_nsif-1_*NSIF-1::GFP+P*_odr-1_*RFP]; YNU865: *ylfEx418 qxIs383*+P *_nsif-1_*ΔSPNSIF-1::GFP+P*_odr-1_*RFP]; YNU868: *ylfEx421 qxIs722* +P*_nsif-1_*NSIF-1::mKate2+ P*_odr-1_*GFP]; YNU869: *ylfEx422 qxIs722*+P *_nsif-1_*ΔSPNSIF-1::mKate2 P*_odr-1_*GFP]; YNU870: *ylfEx423 qxIs722*+P*_col-12_*NSIF-1:: mKate2+ P*_odr-1_*GFP]; YNU871: *ylfEx424 qxIs722*+P*_col-12_*ΔSPNSIF-1:: mKate2 P*_odr-1_*GFP]; YNU872: *ylfEx425 qxIs722*;*nsif-1(ok1337)*+P*_myo-3_*NSIF-1::mKate2+ P*_odr-1_*GFP]; YNU873: *ylfEx426 qxIs722*;*nsif-1(ok1337)*+P*_vha-6_*NSIF-1:: mKate2 P*_odr-1_*GFP]; YNU874: *ylfEx427 qxIs722;nsif-1(ok1337)*+P*_col-12_*NSIF-1:: mKate2+ P*_odr-1_*GFP]; YNU875: *ylfEx428 qxIs722;nsif-1(ok1337)*+P*_col-12_*ΔSPNSIF-1:: mKate2+P*_odr-1_*GFP]; YNU876: *ylfEx429 syb9508*+P*_col-12_*::mKate2]; YNU879: *ylfEx432 sid-1(qt9)*;*qxIs383*+P*_str-3_*NSIF-1::GFP+P*_str-3_*SID-1::3×FLAG+P*_odr-1_*RFP]; YNU880: *ylfEx433 sid-1(qt9)*;*qxIs383*+P*_trx-1_*NSIF-1::GFP+P*_trx-1_*SID-1::3×FLAG+P*_odr-1_*RFP]; YNU881: *ylfEx435 qxIs383*+P*_str-3_*NSIF-1::GFP + P*_odr-1_*RFP]; YNU882: *ylfEx434 qxIs383*+P*_str-3_*NSIF-1::GFP+P*_str-3_*EGL-1::3×FLAG+ P*_odr-1_*RFP]; YNU883: *ylfEx437 qxIs383*+P*_trx-1_*NSIF-1::GFP + P*_odr-1_*RFP]; YNU884: *ylfEx436 qxIs383*+P*_trx-1_*NSIF-1::GFP+P*_trx-1_*EGL-1::3×FLAG+ P*_odr-1_*RFP]; YNU885: *qxIs383*;*nsif-1(ok1337)*;*ylfIs22* P*_vha-6_*NSIF-1::GFP+ P*_odr-1_*RFP]; YNU886: *qxIs383*;*nsif-1(ok1337)*;*ylfIs27* P*_myo-3_*NSIF-1::GFP+ P*_odr-1_*RFP]; YNU887: *ylfEx438 ylfIs1*+P*_str-3_*mKate2]; YNU888: *ylfEx439 ylfIs1*+P*_trx-1_*mKate2]; YNU889: *ylfEx440 ylfIs1*+P*_tph-1_*mKate2]; YNU890: *ylfEx441 ylfIs1*+P*_ggr-2_*mKate2].

### Bacterial strains

*Escherichia coli* OP50, *Escherichia coli* HT115, and *Pseudomonas aeruginosa* PA14 were cultured in LB medium at 37°C. Overnight bacterial cultures were seeded onto Nematode Growth Medium (NGM) plates for subsequent experiments.

### Generation of transgene strains

1. To construct the *C. elegans* plasmid for expression of P*_nsif-1_*::GFP, 490bp promoter of *nsif-1* was inserted into the pPD49.26-GFP vector. DNA plasmid mixture containing P*_nsif-1_*::GFP (20ng/µl) and *unc-76*(+) (p76-16B)(50ng/µl) was injected into the gonads of adult *unc-76(e911)*.
2. To construct the *C. elegans* plasmid for expression of P*_nsif-1_*NSIF-1::GFP, 490bp promoter and 881bp of *nsif-1* genomic DNA was inserted into the pPD49.26-GFP vector. DNA plasmid mixture containing P*_nsif-1_*NSIF-1::GFP (20ng/µl) and *unc-76*(+) (p76-16B)(50ng/µl) was injected into the gonads of adult *unc-76(e911)*.
3. To construct the *C. elegans* plasmid for expression of P*_nsif-1_*ΔSPNSIF-1::GFP, 490bp promoter and 824bp of ΔSP*nsif-1* genomic DNA was inserted into the pPD49.26-GFP vector. DNA plasmid mixture containing P*_nsif-1_*ΔSPNSIF-1::GFP (20ng/µl) and *unc-76*(+) (p76-16B)(50ng/µl) was injected into the gonads of adult *unc-76(e911)*.
4. To construct the *C. elegans* plasmid for expression of P*_nsif-1_*NSIF-1::3×FLAG::GFP, 490bp promoter, 881bp of *nsif-1* genomic DNA and 3×FLAG was inserted into the pPD49.26-GFP vector. DNA plasmid mixture containing P*_nsif-1_*NSIF-1::3×FLAG::GFP (20ng/µl) and *unc-76*(+) (p76-16B)(50ng/µl) was injected into the gonads of adult *unc-76(e911)*.
5. To construct the *C. elegans* plasmid for expression of P*_rgef-1_*NSIF-1::GFP, 3057bp promoter of *rgef-1* and 881bp of *nsif-1* genomic DNA was inserted into the pPD49.26-GFP vector. DNA plasmid mixture containing P*_rgef-1_*NSIF-1::GFP(20ng/µl) and P*_odr-1_*RFP(50ng/µl) was injected into the gonads of adult wild type N2.
6. To construct the *C. elegans* plasmid for expression of P*_myo-3_*NSIF-1::GFP, 2403bp promoter of *myo-3* and 881bp of *nsif-1* genomic DNA was inserted into the pPD49.26-GFP vector. DNA plasmid mixture containing P*_myo-3_*NSIF-1::GFP(20ng/µl) and P*_odr-1_*RFP(50ng/µl) was injected into the gonads of adult wild type N2.
7. To construct the *C. elegans* plasmid for expression of P*_vha-6_*NSIF-1::GFP, 1593bp promoter of *vha-6* and 881bp of *nsif-1* genomic DNA was inserted into the pPD49.26-GFP vector. DNA plasmid mixture containing P*_vha-6_*NSIF-1::GFP(20ng/µl) and P*_odr-1_*RFP(50ng/µl) was injected into the gonads of adult wild type N2.
8. To construct the *C. elegans* plasmid for expression of P*_col-12_*GBP::mKate, 2, 613bp promoter of *col-12* and 345bp of GBP DNA sequence was inserted into the pPD49.26-mKate2 vector. DNA plasmid mixture containing P*_col-12_*GBP::mKate2 (20ng/µl) and *rol-6*(*su1006*) (50ng/µl) was injected into the gonads of adult YNU138: ylfIs8 Prgef-1NSIF-1::GFP+ Podr-1RFP].
9. To construct the *C. elegans* plasmid for expression of P*_rgef-1_*ΔSPNSIF-1::GFP, 3057bp promoter of *rgef-1* and 824bp of ΔSP*nsif-1* genomic DNA sequence was inserted into the pPD49.26-GFP vector. DNA plasmid mixture containing P*_rgef-1_*ΔSPNSIF-1::GFP (20ng/µl), P*_col-12_*GBP::mKate2 (20ng/µl) and *rol-6*(*su1006*) (50ng/µl) was injected into the gonads of wild type N2.
10. To construct the *C. elegans* plasmid for expression of P*_sra-6_*mKate2, 2932bp promoter of *sra-6* was inserted into the pPD49.26-mKate2 vector. DNA plasmid mixture containing P*_sra-6_*mKate2(20ng/µl) and P*_nsif-1_*::GFP(20ng/µl) was injected into the gonads of adult wild type N2.
11. To construct the *C. elegans* plasmid for expression of P*_trx-1_*mKate2, 496bp promoter of *trx-1* was inserted into the pPD49.26-mKate2 vector. DNA plasmid mixture containing P*_trx-1_*mKate2(20ng/µl) and P*_nsif-1_*::GFP(20ng/µl) was injected into the gonads of adult wild type N2.
12. To construct the *C. elegans* plasmid for expression of P*_tph-1_*mKate2, 2023bp promoter of *tph-1* was inserted into the pPD49.26-mKate2 vector. DNA plasmid mixture containing P*_tph-1_*mKate2(20ng/µl) and P*_nsif-1_*::GFP(20ng/µl) was injected into the gonads of adult wild type N2.
13. To construct the *C. elegans* plasmid for expression of P*_ggr-2_*mKate2, 2115bp promoter of *ggr-2* was inserted into the pPD49.26-mKate2 vector. DNA plasmid mixture containing P*_ggr-2_*mKate2(20ng/µl) and P*_nsif-1_*::GFP(20ng/µl) was injected into the gonads of adult wild type N2.
14. To construct the *C. elegans* plasmid for expression of P*_col-12_*NSIF-1::GFP, 613bp promoter of *col-12* and 881bp of *nsif-1* genomic DNA were inserted into the pPD49.26-GFP vector. DNA plasmid mixture containing P*_col-12_*NSIF-1::GFP(20ng/µl) and P*_odr-1_*RFP(50ng/µl) were injected into the gonads of adult XW5399: *qxIs257* P*_ced-1_*NUC-1::CHERRY].
15. To construct the *C. elegans* plasmid for expression of P*_col-12_*ΔSPNSIF-1::GFP, 613bp promoter of *col-12* and 824bp of ΔSP*nsif-1* genomic DNA was inserted into the pPD49.26-GFP vector. DNA plasmid mixture containing P*_col-1_*_2_ΔSPNSIF-1::GFP(20ng/µl) and P*_odr-1_*RFP(50ng/µl) were injected into the gonads of adult XW5399: *qxIs257* P*_ced-1_*NUC-1::CHERRY].
16. To construct the *C. elegans* plasmid for expression of P*_trx-1_*NSIF-1::GFP, 496bp promoter of *trx-1* and 881bp of *nsif-1* genomic DNA was inserted into the pPD49.26-GFP vector. DNA plasmid mixture containing P*_trx-1_*NSIF-1::GFP (20ng/µl) and P*_odr-1_*RFP (50ng/µl) was injected into the gonads of adult YNU467: *nsif-1(ok1337);qxIs383* P*_hyp-7_*NUC-1::CHERRY].
17. To construct the *C. elegans* plasmid for expression of P*_trx-1_*ΔSPNSIF-1::GFP, 496bp promoter of *trx-1* and 824bp of ΔSP*nsif-1* genomic DNA was inserted into the pPD49.26-GFP vector. DNA plasmid mixture containing P*_trx-1_*ΔSPNSIF-1::GFP (20ng/µl) and P*_odr-1_*RFP (50ng/µl) was injected into the gonads of adult YNU467: *nsif-1(ok1337);qxIs383* P*_hyp-7_*NUC-1::CHERRY].
18. To construct the *C. elegans* plasmid for expression of P*_str-3_*NSIF-1::GFP, 259bp promoter of *str-3* and 881bp of *nsif-1* genomic DNA was inserted into the pPD49.26-GFP vector. DNA plasmid mixture containing P*_str-3_*NSIF-1::GFP (20ng/µl) and P*_odr-1_*RFP (50ng/µl) was injected into the gonads of adult YNU467: *nsif-1(ok1337);qxIs383* P*_hyp-7_*NUC-1::CHERRY].
19. To construct the *C. elegans* plasmid for expression of P*_str-3_*ΔSPNSIF-1::GFP, 259bp promoter of *str-3* and 824bp of ΔSP*nsif-1* genomic DNA was inserted into the pPD49.26-GFP vector. DNA plasmid mixture containing P*_str-3_*ΔSPNSIF-1::GFP (20ng/µl) and P*_odr-1_*RFP (50ng/µl) was injected into the gonads of adult YNU467: *nsif-1(ok1337);qxIs383* P*_hyp-7_*NUC-1::CHERRY].
20. To construct the *C. elegans* plasmid for expression of P*_str-3_*EGL-1::3×FLAG, 259bp promoter of *str-3* and 875bp of EGL-1 genomic DNA was inserted into the pPD49.26-3×FLAG vector. DNA plasmid mixture containing P*_str-3_*EGL-1::3×FLAG (25ng/µl) and P*_odr-1_*RFP (50ng/µl) was injected into the gonads of adult XW17096:*qxIs383* P*_hyp-7_*NUC-1::CHERRY].
21. To construct the *C. elegans* plasmid for expression of P*_trx-1_*EGL-1::3×FLAG, 496bp promoter of *trx-1* and 875bp of EGL-1 genomic DNA was inserted into the pPD49.26-3×FLAG vector. DNA plasmid mixture containing P*_trx-1_*EGL-1::3×FLAG (25ng/µl) and P*_odr-1_*RFP (50ng/µl) was injected into the gonads of adult XW17096: *qxIs383* P*_hyp-7_*NUC-1::CHERRY].
22. To construct the *C. elegans* plasmid for expression of P*_str-3_*NSIF-1::mKate2, 259bp promoter of *str-3* and 881bp of *nsif-1* genomic DNA was inserted into the pPD49.26-mKate2 vector. DNA plasmid mixture containing P*_str-3_*NSIF-1::mKate2 (20ng/µl) and P*_odr-1_*GFP (50ng/µl) was injected into the gonads of adult YNU473: *nsif-1(ok1337);qxIs722* P*_dpy-7_*DPY-7::sfGFP, single-copy insertion].
23. To construct the *C. elegans* plasmid for expression of P*_str-3_*ΔSPNSIF-1::mKate2, 259bp promoter of *str-3* and 824bp of *nsif-1* genomic DNA was inserted into the pPD49.26-mKate2 vector. DNA plasmid mixture containing P*_str-3_*ΔSPNSIF-1::mKate2 (20ng/µl) and P*_odr-1_*GFP (50ng/µl) was injected into the gonads of adult YNU473: *nsif-1(ok1337);qxIs722* P*_dpy-7_*DPY-7::sfGFP, single-copy insertion].
24. To construct the *C. elegans* plasmid for expression of P*_trx-1_*NSIF-1::mKate2, 496bp promoter of *trx-1* and 881bp of *nsif-1* genomic DNA was inserted into the pPD49.26-mKate2 vector. DNA plasmid mixture containing P*_trx-1_*NSIF-1::mKate2 (20ng/µl) and P*_odr-1_*GFP (50ng/µl) was injected into the gonads of adult YNU473: *nsif-1(ok1337);qxIs722* P*_dpy-7_*DPY-7::sfGFP, single-copy insertion].
25. To construct the *C. elegans* plasmid for expression of P*_trx-1_*ΔSPNSIF-1:: mKate2, 496bp promoter of *trx-1* and 824bp of *nsif-1* genomic DNA was inserted into the pPD49.26-mKate2 vector. DNA plasmid mixture containing P*_trx-1_*ΔSPNSIF-1::mKate2 (20ng/µl) and P*_odr-1_*GFP (50ng/µl) was injected into the gonads of adult YNU473: *nsif-1(ok1337);qxIs722* P*_dpy-7_*DPY-7::sfGFP, single-copy insertion].
26. To construct the *C. elegans* plasmid for expression of P*_nsif-1_*NSIF-1::mKate2, 490bp promoter of *nsif-1* and 881bp of *nsif-1* genomic DNA was inserted into the pPD49.26-mKate2 vector. DNA plasmid mixture containing P*_nsif-1_*NSIF-1::mKate2 (20ng/µl) and P*_odr-1_*GFP(50ng/µl) was injected into the gonads of adult YNU473: *nsif-1(ok1337);qxIs722* P*_dpy-7_*DPY-7::sfGFP, single-copy insertion].
27. To construct the *C. elegans* plasmid for expression of P*_nsif-1_*ΔSPNSIF-1::mKate2, 490bp promoter of *nsif-1* and 824bp of *nsif-1* genomic DNA was inserted into the pPD49.26-mKate2 vector. DNA plasmid mixture containing P*_nsif-1_*NSIF-1::mKate2 (20ng/µl) and P*_odr-1_*GFP(50ng/µl) was injected into the gonads of adult YNU473: *nsif-1(ok1337);qxIs722* P*_dpy-7_*DPY-7::sfGFP, single-copy insertion].
28. To construct the *C. elegans* plasmid for expression of P*_str-3_*SID-1::3×FLAG, 259bp promoter of *str-3* and 2331bp of *sid-1* cDNA was inserted into the pPD49.26-GFP vector. DNA plasmid mixture containing P*_str-3_*SID-1::3×FLAG (20ng/µl) and P*_odr-1_*RFP (50ng/µl) was injected into the gonads of adult YNU778: *sid-1(qt9);qxIs383* P*_hyp-7_*NUC-1::CHERRY].
29. To construct the *C. elegans* plasmid for expression of P*_trx-1_*SID-1::3×FLAG, 496bp promoter of *trx-1* and 2331bp of *sid-1* cDNA was inserted into the pPD49.26-GFP vector. DNA plasmid mixture containing P*_trx-1_*SID-1::3×FLAG (20ng/µl) andP*_odr-1_*RFP (50ng/µl) was injected into the gonads of adult YNU778: *sid-1(qt9);qxIs383* P*_hyp-7_*NUC-1::CHERRY].
30. To construct the *C. elegans* plasmid for expression of P*_str-3_*EGL-1::3×FLAG, 259bp promoter of *str-3* and 875bp of EGL-1 genomic DNA was inserted into the pPD49.26-3×FLAG vector. DNA plasmid mixture containing P*_str-3_*EGL-1::3×FLAG (25ng/µl) and *rol-6*(*su1006*) (50ng/µl) was injected into the gonads of adult YNU765: *ylfIs52* P*_col-12_*GBP::mKate2];*ylfIs8* P*_rgef-1_*NSIF-1::GFP+ P*_odr-1_*RFP].
31. To construct the *C. elegans* plasmid for expression of P*_trx-1_*EGL-1::3×FLAG, 496bp promoter of *trx-1* and 875bp of EGL-1 genomic DNA was inserted into the pPD49.26-3×FLAG vector. DNA plasmid mixture containing P*_trx-1_*EGL-1::3×FLAG (25ng/µl) and *rol-6*(*su1006*) (50ng/µl) was injected into the gonads of adult YNU765: *ylfIs52* P*_col-12_*GBP::mKate2];*ylfIs8* P*_rgef-1_*NSIF-1::GFP+ P*_odr-1_*RFP].
32. To construct the *C. elegans* expression of P*_str-3_*NSIF-1::GFP in YNU747: *ylfIs52* P*_col-12_*GBP::mKate2], DNA plasmid mixture containing P*_str-3_*NSIF-1::GFP (25ng/µl) and *rol-6*(*su1006*) (50ng/µl) was injected into the gonads of adult YNU747:*ylfIs52* P*_col-12_*GBP::mKate2].
33. To construct the *C. elegans* expression of P*_str-3_*ΔSPNSIF-1::GFP in YNU747: *ylfIs52* P*_col-12_*GBP::mKate2], DNA plasmid mixture containing P*_str-3_*ΔSPNSIF-1::GFP (25ng/µl) and *rol-6*(*su1006*) (50ng/µl) was injected into the gonads of adult YNU747:*ylfIs52* P*_col-12_*GBP::mKate2].
34. To construct the *C. elegans* expression of P*_trx-1_*NSIF-1::GFP in YNU747: *ylfIs52* P*_col-12_*GBP::mKate2], DNA plasmid mixture containing P*_trx-1_*NSIF-1::GFP (25ng/µl) and *rol-6*(*su1006*) (50ng/µl) was injected into the gonads of adult YNU747:*ylfIs52* P*_col-12_*GBP::mKate2].
35. To construct the *C. elegans* expression of P*_trx-1_*ΔSPNSIF-1::GFP in YNU747: *ylfIs52* P*_col-12_*GBP::mKate2], DNA plasmid mixture containing P*_trx-1_*ΔSPNSIF-1::GFP (25ng/µl) and *rol-6*(*su1006*) (50ng/µl) was injected into the gonads of adult YNU747:*ylfIs52* P*_col-12_*GBP::mKate2].
36. To construct the *C. elegans* expression of P*_col-12_*NSIF-1::GFP in YNU467: *nsif-1(ok1337);qxIs383* P*_hyp-7_*NUC-1::CHERRY], DNA plasmid mixture containing P*_col-12_*NSIF-1::GFP (25ng/µl) and P*_odr-1_*RFP (50ng/µl) was injected into the gonads of adult YNU467:*nsif-1(ok1337);qxIs383* P*_hyp-7_*NUC-1::CHERRY].
37. To construct the *C. elegans* expression of P*_col-12_*ΔSPNSIF-1::GFP in YNU467: *nsif-1(ok1337);qxIs383* P*_hyp-7_*NUC-1::CHERRY], DNA plasmid mixture containing P*_col-12_*ΔSPNSIF-1::GFP (25ng/µl) and P*_odr-1_*RFP (50ng/µl) was injected into the gonads of adult YNU467:*nsif-1(ok1337);qxIs383* P*_hyp-7_*NUC-1::CHERRY].
38. To construct the *C. elegans* expression of P*_nsif-1_*NSIF-1::GFP in XW17096: *qxIs383* P*_hyp-7_*NUC-1::CHERRY], DNA plasmid mixture containing P*_nsif-1_*NSIF-1::GFP (25ng/µl) and P*_odr-1_*RFP (50ng/µl) was injected into the gonads of adult XW17096:*qxIs383* P*_hyp-7_*NUC-1::CHERRY].
39. To construct the *C. elegans* expression of P*_nsif-1_*ΔSPNSIF-1::GFP in XW17096: *qxIs383* P*_hyp-7_*NUC-1::CHERRY], DNA plasmid mixture containing P*_nsif-1_*ΔSPNSIF-1::GFP (25ng/µl) and P*_odr-1_*RFP (50ng/µl) was injected into the gonads of adult XW17096:*qxIs383* P*_hyp-7_*NUC-1::CHERRY].
40. To construct the *C. elegans* expression of P*_nsif-1_*NSIF-1::mKate2 in XW18042: *qxIs722* P*_dpy-7_*DPY-7::sfGFP, single-copy insertion], DNA plasmid mixture containing P*_nsif-1_*NSIF-1:: mKate2 (25ng/µl) and P*_odr-1_*GFP (50ng/µl) was injected into the gonads of adult XW18042: *qxIs722* P*_dpy-7_*DPY-7::sfGFP, single-copy insertion].
41. To construct the *C. elegans* expression of P*_nsif-1_*ΔSPNSIF-1:: mKate2 in XW18042: *qxIs722* P*_dpy-7_*DPY-7::sfGFP, single-copy insertion], DNA plasmid mixture containing P*_nsif-1_*ΔSPNSIF-1::mKate2 (25ng/µl) and P*_odr-1_*GFP (50ng/µl) was injected into the gonads of adult XW18042: *qxIs722* P*_dpy-7_*DPY-7::sfGFP, single-copy insertion].
42. To construct the *C. elegans* expression of P*_col-12_*NSIF-1::mKate2 in XW18042: *qxIs722* P*_dpy-7_*DPY-7::sfGFP, single-copy insertion], DNA plasmid mixture containing P*_col-12_*NSIF-1::mKate2 (25ng/µl) and P*_odr-1_*GFP (50ng/µl) was injected into the gonads of adult XW18042: *qxIs722* P*_dpy-7_*DPY-7::sfGFP, single-copy insertion].
43. To construct the *C. elegans* expression of P*_col-12_*ΔSPNSIF-1:: mKate2 in XW18042: *qxIs722* P*_dpy-7_*DPY-7::sfGFP, single-copy insertion], DNA plasmid mixture containing P*_col-12_*ΔSPNSIF-1:: mKate2 (25ng/µl) and P*_odr-1_*GFP (50ng/µl) was injected into the gonads of adult XW18042: *qxIs722* P*_dpy-7_*DPY-7::sfGFP, single-copy insertion].
44. To construct the *C. elegans* plasmid for expression of P*_myo-3_*NSIF-1::mKate2, 2403bp promoter of *myo-3* and 881bp of *nsif-1* genomic DNA was inserted into the pPD49.26-mKate2 vector, DNA plasmid mixture containing P*_myo-3_*NSIF-1::mKate2 (25ng/µl) and P*_odr-1_*GFP (50ng/µl) was injected into the gonads of adult YNU473: *nsif-1(ok1337);qxIs722* P*_dpy-7_*DPY-7::sfGFP, single-copy insertion].
45. To construct the *C. elegans* plasmid for expression of P*_vha-6_*NSIF-1::mKate2, 1593bp promoter of *myo-3* and 881bp of *nsif-1* genomic DNA was inserted into the pPD49.26-mKate2 vector, DNA plasmid mixture containing P*_vha-6_*NSIF-1::mKate2 (25ng/µl) and P*_odr-1_*GFP (50ng/µl) was injected into the gonads of adult YNU473: *nsif-1(ok1337);qxIs722* P*_dpy-7_*DPY-7::sfGFP, single-copy insertion].
46. To construct the *C. elegans* expression of P*_col-12_*NSIF-1::mKate2 in YNU473: *nsif-1(ok1337);qxIs722* P*_dpy-7_*DPY-7::sfGFP, single-copy insertion], DNA plasmid mixture containing P*_col-12_*NSIF-1::mKate2 (25ng/µl) and P*_odr-1_*GFP (50ng/µl) was injected into the gonads of adult YNU473: *nsif-1(ok1337);qxIs722* P*_dpy-7_*DPY-7::sfGFP, single-copy insertion].
47. To construct the *C. elegans* expression of P*_col-12_*ΔSPNSIF-1:: mKate2 in YNU473: *nsif-1(ok1337);qxIs722* P*_dpy-7_*DPY-7::sfGFP, single-copy insertion], single-copy insertion], DNA plasmid mixture containing P*_col-12_*ΔSPNSIF-1:: mKate2 (25ng/µl) and P*_odr-1_*GFP (50ng/µl) was injected into the gonads of adult YNU473: *nsif-1(ok1337);qxIs722* P*_dpy-7_*DPY-7::sfGFP, single-copy insertion].
48. To construct the *C. elegans* plasmid for expression of P*_col-12_*mKate2, 613bp promoter of *col-12* was inserted into the pPD49.26-mKate2 vector, DNA plasmid mixture containing P*_col-12_*mKate2 (25ng/µl) was injected into the gonads of adult BHX9508: *nsif-1(syb9508) nsif-1*::mNeonGreen::3xFLAG knock-in].
49. To construct the *C. elegans* plasmid for expression of P*_str-3_*SID-1::3×FLAG and P*_str-3_*NSIF-1::GFP into YNU778: *sid-1(qt9);qxIs383* P*_hyp-7_*NUC-1::CHERRY], DNA plasmid mixture containing P*_str-3_*SID-1::3×FLAG (20ng/µl), P*_str-3_*NSIF-1::GFP (20ng/µl) and P*_odr-1_*RFP (50ng/µl) was injected into the gonads of adult YNU778: *sid-1(qt9);qxIs383* P*_hyp-7_*NUC-1::CHERRY].
50. To construct the *C. elegans* plasmid for expression of P*_trx-1_*SID-1::3×FLAG and P*_trx-1_*NSIF-1::GFP into YNU778: *sid-1(qt9);qxIs383* P*_hyp-7_*NUC-1::CHERRY], DNA plasmid mixture containing P*_trx-1_*SID-1::3×FLAG (20ng/µl), P*_trx-1_*NSIF-1::GFP (20ng/µl) and P*_odr-1_*RFP (50ng/µl) was injected into the gonads of adult YNU778: *sid-1(qt9);qxIs383* P*_hyp-7_*NUC-1::CHERRY].
51. To construct the *C. elegans* expression of P*_str-3_*NSIF-1::GFP in XW17096: *qxIs383* P*_hyp-7_*NUC-1::CHERRY], DNA plasmid mixture containing P*_str-3_*NSIF-1::GFP (25ng/µl) and P*_odr-1_*RFP (50ng/µl) was injected into the gonads of adult XW17096: *qxIs383* P*_hyp-7_*NUC-1::CHERRY].
52. To construct the *C. elegans* expression of P*_str-3_*EGL-1::3×FLAG and P*_str-3_*NSIF-1::GFP in XW17096: *qxIs383* P*_hyp-7_*NUC-1::CHERRY], DNA plasmid mixture containing P*_str-3_*EGL-1::3×FLAG (25ng/µl), P*_str-3_*NSIF-1::GFP (25ng/µl) and P*_odr-1_*RFP (50ng/µl) was injected into the gonads of adult XW17096: *qxIs383* P*_hyp-7_*NUC-1::CHERRY].
53. To construct the *C. elegans* expression of P*_trx-1_*NSIF-1::GFP in XW17096: *qxIs383* P*_hyp-7_*NUC-1::CHERRY], DNA plasmid mixture containing P*_trx-1_*NSIF-1::GFP (25ng/µl) and P*_odr-1_*RFP (50ng/µl) was injected into the gonads of adult XW17096: *qxIs383* P*_hyp-7_*NUC-1::CHERRY].
54. To construct the *C. elegans* expression of P*_trx-1_*EGL-1::3×FLAG and P*_trx-1_*NSIF-1::GFP in XW17096: *qxIs383* P*_hyp-7_*NUC-1::CHERRY], DNA plasmid mixture containing P*_trx-1_*EGL-1::3×FLAG (25ng/µl), P*_trx-1_*NSIF-1::GFP (25ng/µl) and P*_odr-1_*RFP (50ng/µl) was injected into the gonads of adult XW17096: *qxIs383* P*_hyp-7_*NUC-1::CHERRY].
55. To construct the *C. elegans* plasmid for expression of P*_str-3_*mKate2 in YNU22: *unc-76(e911);ylfIs1* P*_nsif-1_*NSIF-1::GFP*; unc-76*(+)], 259bp promoter of *str-3* was inserted into the pPD49.26-mKate2 vector. DNA plasmid mixture containing P*_str-3_*mKate2 (20ng/µl) was injected into the gonads of adult YNU22: *unc-76(e911);ylfIs1* P*_nsif-1_*NSIF-1::GFP*; unc-76*(+)].
56. To construct the *C. elegans* plasmid for expression of P*_trx-1_*mKate2 in YNU22: *unc-76(e911);ylfIs1* P*_nsif-1_*NSIF-1::GFP*; unc-76*(+)], 496bp promoter of *trx-1* was inserted into the pPD49.26-mKate2 vector. DNA plasmid mixture containing P*_trx-1_*mKate2 (20ng/µl) was injected into the gonads of adult YNU22: *unc-76(e911);ylfIs1* P*_nsif-1_*NSIF-1::GFP*; unc-76*(+)],.
57. To construct the *C. elegans* plasmid for expression of P*_tph-1_*mKate2 in YNU22: *unc-76(e911);ylfIs1* P*_nsif-1_*NSIF-1::GFP*; unc-76*(+)], 2023bp promoter of *tph-1* was inserted into the pPD49.26-mKate2 vector. DNA plasmid mixture containing P*_tph-1_*mKate2 (20ng/µl) was injected into the gonads of adult YNU22: *unc-76(e911);ylfIs1* P*_nsif-1_*NSIF-1::GFP*; unc-76*(+)].
58. To construct the *C. elegans* plasmid for expression of P*_ggr-2_*mKate2 in YNU22: *unc-76(e911);ylfIs1* P*_nsif-1_*NSIF-1::GFP*; unc-76*(+)], 2115bp promoter of *ggr-2* was inserted into the pPD49.26-mKate2 vector. DNA plasmid mixture containing P*_ggr-2_*mKate2 (20ng/µl) was injected into the gonads of adult YNU22: *unc-76(e911);ylfIs1* P*_nsif-1_*NSIF-1::GFP*; unc-76*(+)].

### Generating knock-out mutants using CRISPR

1. **Construct CRISPR-Cas9-sgRNA vector.** The *nsif-1* sgRNA sequence (5′-AGCGCAGTACACGAGGCAT-3-3′) was designed from https://crispor.gi.ucsc.edu/, and cloned into the CRISPR-Cas9-sgRNA vector pDD162 (Addgene #47549). We constructed the plasmids: P*_eft-3_*Cas9 + P*_u6_nsif-1*-sgRNA
2. **Design of the repair template.** The single-stranded DNA (ssDNA) repair template was designed with homology arms of at least 40 base pairs (bp) flanking the target editing site. To enable efficient screening and validation of successful genome editing, a NheI restriction site—along with a premature stop codon—was incorporated at the desired insertion site. To ensure optimal synthesis fidelity and efficiency, the total length of the ssDNA repair template was constrained to no more than 120 bp. The sequence of repair template: *nsif-1* 5′-ccagccgcagtcgcatgttctaatgctgtttacgatgcctgctagcgctttatccaacagaagacgaacaagggtatccagct-3′. *dpy-10* 5′-cacttgaacttcaatacggcaagatgagaatgactggaaaccgtaccgcatgcggtgcctatggtagcttcacatggcttcagaccaacagcctat-3′.
3. **Injection of DNA mixture and genotyping** During microinjection, we co-injected into P0 gonads: The Cas9-sgRNA plasmid (P*_eft-3_*Cas9 + P*_u6_nsif-1*-sgRNA) (25ng/µl), The *nsif-1* ssDNA repair template (2µM), The co-editing marker plasmid (P*_eft-3_*::Cas9 + P*_u6_*::*dpy-10*-sgRNA) (20ng/μl), and The *dpy-10* ssDNA repair template (2µM). Worm lysis and PCR: Worms were picked into 10 μl of worm lysis buffer (50mM KCl,10mM Tris-HCl pH 8.0, 2.5mM MgCl2, 0.45% NP40, 0.45% Tween-20, 0.01% Gelactin, 0.2mg/mL Proteinase K), quickly freeze-thaw three times using liquid nitrogen, incubated it at 60 °C for 90min and 95 °C for 20min. 1 μl supernatant was taken and performed for PCR analysis with the following primers: *nsif-1* KO forward 5′-gttaagatcgtggaaagcttgctagt-3′, reverse 5′-ctcctgcattccgacgc-3′, then digested with NheI endonuclease 37 overnight and identified by DNA agarose electrophoresis. We first identified F1 Roller progeny with successful *dpy-10* editing and genotyped them for *nsif-1* editing using NheI digestion. Individuals F1 showing the expected digestion pattern (three bands) were retained and propagated to the F2 generation. Single non-Roller F2 animals were isolated and re-genotyped. F2 worms that maintained the NheI-cut pattern (two bands) were mutant animals.

### Generation of EPIC-1::GFP knock-in strain using CRISPR/Cas9

1. **Construct knock-in CRISPR-Cas9-sgRNA vector.** To generate an EPIC-1::GFP knock-in strain, a fluorescent GFP tag was inserted at the C-terminus of the endogenous *epic-1* coding sequence using CRISPR–Cas9. The *epic-1* sgRNA sequence (5′-TCAGGAAGAGACAGCTCCAG-3′) was designed using the CRISPOR online tool (https://crispor.gi.ucsc.edu/) and cloned into the CRISPR-Cas9-sgRNA vector pDD162 (Addgene #47549)
2. **Construct knock-in homologous recombination vector** Homology-directed repair was carried out using the pDD282 plasmid (Addgene #66823), which contains GFP::3×FLAG and serves as a repair template. Approximately 800 bp homology arms flanking the *epic-1* locus were amplified from *C. elegans* genomic DNA and inserted into pDD282 to enhance recombination efficiency.
3. **Injection of DNA mixture and Genotyping** A DNA mixture consisting of 50 ng/μl Cas9-sgRNA plasmid (pDD162-*epic-1* sgRNA), 10 ng/μl homologous recombination plasmid (pDD282-*epic-1* homology recombinant template), and 2.5 ng/μl pCFJ90 (P*_myo-2_*mCherry) as a co-injection marker was injected into the gonads of young adult N2 hermaphrodites. After hygromycin selection, transgenic F1 progeny expressing P*_myo-2_*mCherry (co-conversion marker) and displaying the roller phenotype were isolated and screened for GFP knock-in by PCR. Positive candidates were verified by Sanger sequencing. Confirmed EPIC-1::GFP strains were subsequently outcrossed with N2 wild-type worms three times before use in further experiments. More details can be found in previous studies [77].

### Generation of transgene strains by TMP/UV

Stable transgenic *C. elegans* lines were generated using UV/trimethylpsoralen (TMP)-mediated integration. Synchronized L4-stage transgenic worms were first treated with TMP (50 µg/mL) to enhance UV sensitivity, followed by UV irradiation (350–400 J/m²) to induce DNA damage and facilitate genomic integration of extrachromosomal arrays. After recovery and expansion, single-worm lines were screened for stable inheritance of both the co-marker and target plasmid expression. Positive integrations were subsequently outcrossed with wild-type N2 worms at least four times to remove background mutations.

### *C. elegans* RNAi feeding strategy

RNAi by feeding was performed using bacterial clones from the MRC RNAi library [78] or the ORF-RNAi Library [79]. Synchronized *C. elegans* (L1-stage larvae) were transferred onto RNAi NGM plates seeded with *E. coli* HT115 expressing the corresponding double-stranded RNA. Worms were maintained at 20 °C and allowed to develop either to the L4 larval stage or the next generation, depending on the specific experimental requirements.

### *C. elegans* tissue-specific RNAi feeding strategy

To achieve tissue-specific RNA interference, we cross *sid-1(qt9)* mutant into the background (*qxIs383*) to abolish systemic RNAi. The *sid-1* gene was then selectively restored in either ASI or ASJ neurons by expressing *sid-1* under the control of neuron-specific promoters (P*_str-3_*SID-1::3×FLAG for ASI and P*_trx-1_*SID-1::3×FLAG for ASJ).

In these transgenic backgrounds, feeding RNAi was performed to knock down target genes. Briefly, worms were transferred onto NGM plates seeded with RNAi bacteria and incubated at 20℃ until the indicated developmental stages. The L4440 empty vector strain was used as a negative control.

### *Pseudomonas aeruginosa* PA14 survival assay

The slow-killing assay was performed as previously described [80]. Overnight cultures of *Pseudomonas aeruginosa* PA14 were inoculated onto modified NGM agar plates (0.35% peptone, 35 mm in diameter). The inoculated plates were air-dried at room temperature and subsequently incubated at 37°C for 24 hours, followed by an additional incubation at 25°C for 8-24 hours. Before initiating the assay, plates were equilibrated at room temperature for at least 1 hour. Synchronized L4-stage *C. elegans* were transferred onto PA14-seeded slow-killing plates and maintained at 25°C. Survival was assessed at 12-hour intervals, with mortality determined by the absence of response to mechanical stimulation of both the anterior and posterior regions using an eyebrow hair. To suppress progeny production and prevent “bagging,” 5-fluoro-2’-deoxyuridine (FUdR, 50 mg/mL; Sigma) was added to the assay medium. Worms exhibiting vulval rupture or those that migrated off the agar surface were excluded from analysis. All survival assays were conducted in triplicate to ensure statistical robustness.

### Quantitative RT-PCR

1. Sample preparation Synchronized *C. elegans* were grown to the L4 stage and washed with M9 buffer to remove residual *E. coli* OP50. Worms were then transferred to NGM plates seeded with *Pseudomonas aeruginosa* PA14 and incubated for 24 hours. Following infection, worms were washed with M9 buffer to remove surface bacteria. For uninfected samples, N2 and *nsif-1(ok1337)* worms were collected at L1+24h or the designated time point using the same procedure.
2. RNA isolation Worms were washed with M9 buffer, pelleted by centrifugation, and snap-frozen in liquid nitrogen. Samples from each 90 mm plate were homogenized in 1 mL TRIzol reagent using a tissue homogenizer. After adding 200 µL chloroform, samples were vigorously shaken and centrifuged at 12,000 × g for 15 minutes at 4°C. The aqueous phase was transferred to a fresh tube and mixed with an equal volume of isopropanol to precipitate RNA. The RNA pellet was collected by centrifugation, washed three times with 75% ethanol, air-dried, and dissolved in RNase-free water. Samples were then incubated at 55–60°C for 10 minutes to ensure complete dissolution.
3. RT-PCR cDNA was synthesized using the PrimeScript RT Reagent Kit (TaKaRa) and diluted to a final concentration of 100 ng/µL. For qRT-PCR, 2.5 µL of diluted cDNA was used as a template in a 10 µL reaction mixture containing PowerUp SYBR Green Master Mix (Thermo Fisher Scientific). Reactions were performed on an Applied Biosystems QuantStudio 7 Flex Real-Time PCR System (Life Technologies). The *act-1* gene was used as an internal reference, and fold changes in gene expression were calculated using the 2^-ΔΔCt^ method. Each experiment was conducted in triplicate, with at least three independent biological replicates. Primers are listed in S4 Table

### RNA-seq Preparation and Analysis

1. Preparation of *C. elegans* Samples for RNA-seq RNA-seq was performed using three independently generated biological replicates. Synchronized L1+12h stage animals (*N2* wild-type and *nsif-1(ok1337)* mutants) grown on *E. coli* OP50 were collected for RNA extraction.
2. RNA Sequencing and Data Processing For each sample, 1 mg of total RNA was used as input for library preparation. Sequencing libraries were constructed using the NEBNext Ultra RNA Library Prep Kit for Illumina (NEB, USA) according to the manufacturer’s protocol, with index codes incorporated for sample identification. Libraries were sequenced on an Illumina platform to generate paired-end reads. Raw sequencing data were processed using the BMKCloud (www.biocloud.net) online bioinformatics pipeline for quality control and downstream analysis.

### Fluorescent microscopy

Worms were anesthetized using 1 mM levamisole and mounted on 3% agarose pads for imaging. Fluorescence microscopy was performed using either an upright Leica DM6 B microscope equipped with a 10× objective lens or a Zeiss LSM 900 confocal laser scanning microscope with a 60× oil immersion objective. Fluorescence signals were detected using the following filter sets: GFP (excitation: 488 nm, emission: 510–550 nm) and RFP/CHERRY/mKate2 (excitation: 561 nm, emission: 570–620 nm). Images were acquired and processed using ImageJ (NIH) or ZEN imaging software v3.4 (Carl Zeiss Inc.).

### Protein extraction from *C. elegans*

Worms were collected and washed three times with M9 buffer to remove residual bacteria. The worm pellet was then resuspended in protein lysis buffer (50 mM Tris-HCl, pH 8.0; 50 mM NaCl; 0.5% deoxycholate; 10% glycerol; 1% NP-40) supplemented with PMSF (Sangon Biotech), Protease Inhibitor Cocktail (MCE, HY-K0011), and Phosphatase Inhibitor Cocktail I (MCE, HY-K0021). Worms were homogenized and lysed, followed by centrifugation at 12,000 rpm for 10 min at 4°C. The supernatant was collected, and protein concentration was determined using the Pierce BCA Protein Assay Kit (Thermo Fisher, 23227).

### Western blot

Protein samples were prepared by adding 5× SDS loading buffer to the quantified protein at a 1:4 ratio. Samples were then heated at 95°C for 10 minutes to denature the proteins. Proteins were separated via SDS-polyacrylamide gel electrophoresis (SDS-PAGE) and transferred onto a PVDF membrane for subsequent analysis.

To assess CHERRY expression, the membrane was incubated with anti-CHERRY antibody (1:3000; Proteintech, 26765-1-AP), with anti-β-actin (1:3000; Proteintech, 66009-1-lg) as a reference protein. Following primary antibody incubation, the membrane was washed and incubated with appropriate secondary antibodies. Protein bands were visualized using chemiluminescence detection.

### Isolation of secreted proteins from *C. elegans*

Synchronized L1-stage worms were cultured in 500 μL of M9 buffer, with or without the addition of 5 μL *Pseudomonas aeruginosa* PA14 (OD600 = 1) or *E. coli* OP50, in a 1.5 mL tube. The samples were gently shaken for 4 hours. Following incubation, worms were pelleted by centrifugation at 50 × g for 1 minute, and the supernatant was collected.

The collected supernatant was filtered through a 0.22 μm filter unit to isolate secreted proteins. The resulting protein samples were then subjected to mass spectrometry analysis using an Orbitrap hybrid mass spectrometer (Orbitrap Exploris 480, Thermo Scientific).

We selected synchronized L1-stage animals for the secretome analysis because bleach synchronized L1s have never been exposed to any bacterial food source, and therefore lack any prior innate immune history. When these naïve L1s are subsequently challenged with *P. aeruginosa* PA14, the secreted proteins they produce should reflect a direct response to PA14 itself, uncontaminated by adaptive effects resulting from a previous diet of *E. coli* OP50. In contrast, L4 stage animals have been reared on *E. coli* for approximately 48 h and may have already developed a baseline level of innate immune adaptation, which could attenuate or otherwise modify their response upon transfer to PA14.

### Quantification of lysosomal morphology

1. **Quantification of number of Tubular lysosomes** Fluorescence images of adult *C. elegans* expressing NUC-1::CHERRY [(XW5399: *qxIs257*(P*_ced-1_*NUC-1::CHERRY) and XW17096:*qxIs383*(P*_hyp-7_*NUC-1::CHERRY)]) at different ages or specific time points were captured using a laser scanning confocal microscope (Zeiss LSM900). The length of NUC-1::CHERRY-positive tubules in each worm was quantified using ImageJ software. Tubular lysosomes that intersected were counted as two separate tubules. For each worm, the number of tubular lysosomes within a 35 × 25 μm² unit area was quantified, with a minimum of 20 animals analyzed per strain at different time points.
2. **Quantification of vesicular lysosome number and volume** Fluorescence images of *C. elegans* adults expressing NUC-1::CHERRY (XW5399: *qxIs257*(P*_ced-1_*NUC-1::CHERRY) and XW17096:*qxIs383*(P*_hyp-7_*NUC-1::CHERRY)] were acquired using confocal fluorescence microscopy (Carl Zeiss LSM900). The volume and number of NUC-1::CHERRY-positive vesicular lysosomes within a 35 × 25 μm^²^ unit area per animal were quantified using ImageJ and ZEN Blue v3.4 software. A minimum of 20 animals were analyzed under different conditions.

All images were acquired using a Zeiss LSM 900 confocal microscope with a 63× oil-immersion objective at a resolution of 1024 x1024 pixels. For quantification, a fixed ROI (Region of Interest) of 35 × 25 μm² was generated and saved as a template in ZEN software. Because all images were acquired under identical imaging parameters, the same ROI template was applied uniformly across all images. To ensure that the ROI was positioned in the same anatomical region in every animal, all images used for lysosome quantification and cuticle morphology analysis were acquired from a defined body region located posterior to the head (terminal pharyngeal bulb) and anterior to the vulva. The ROI was placed within this region using these anatomical landmarks as references and applied consistently across all worms analyzed.

### Lysosome maturation process

L4-stage worms were subjected to heat shock at 34°C for 30 minutes to induce transient expression of NUC-1::sfGFP::CHERRY strain: XW13734: *qxIs612* P*_hsp_*NUC-1::sfGFP::CHERRY], followed by recovery at 20°C for 1, 3, 6, and 12 hours. At each time point, different worms were selected and examined using confocal fluorescence microscopy (Carl Zeiss LSM900). To assess lysosome maturation under various conditions, worms were heat-shocked and analyzed at different time points post-recovery. The colocalization of CHERRY and GFP was quantified using Pearson’s correlation coefficient in the ImageJ colocalization tool.

### Quantification of NUC-1::pHTomato fluorescence intensity

Different *C. elegans* strains were crossed with the NUC-1::pHTomato strain XW19180: *qxIs750* [P*_hsp_*NUC-1::pHTomato] to generate homozygous lines. Synchronized worms were cultured to the L4 stage and subjected to heat shock at 34°C for 30 minutes. Immediately after heat shock, worms were transferred to a 20°C incubator for a 12-hour recovery period, followed by fluorescence imaging using a confocal microscope. For *Pseudomonas aeruginosa* PA14 treatment, L4-stage worms were transferred to NGM plates seeded with PA14 before undergoing the same heat shock and recovery protocol. To ensure consistency, laser intensity and exposure time were maintained uniformly during imaging. Fluorescence intensity was quantified by measuring the 150 lysosomes from 20 worms per strain Each lysosome’s fluorescence intensity was divided by the mean fluorescence intensity of all control-group lysosomes. Specifically, for the control group, each lysosome’s value was normalized to the control mean (yielding a distribution centered around 1.0). For the mutant group, each lysosome’s value was likewise divided by the control mean, so that the mutant values can be read as fold-change relative to control.

### Quantitative analysis of ELT-3 fluorescence intensity in nuclear

ELT-3::sfGFP-expressing *C. elegans* (OP559: *wgIs559*) were imaged using confocal fluorescence microscopy (Carl Zeiss LSM900). Nuclear-localized ELT-3::sfGFP fluorescence intensity was quantified using ZEN Blue v3.4 software. To ensure reliability, at least 20 animals per strain were analyzed at different developmental stages. Fluorescence intensity within the nucleus was measured and compared to evaluate ELT-3::sfGFP expression levels across samples.

### Quantification of lysosomal degradation activity

L4-stage *C. elegans* expressing NUC-1::CHERRY in the epidermis [XW5399: *qxIs257*(P*_ced-1_*NUC-1::CHERRY) and XW17096:*qxIs383*(P*_hyp-7_*NUC-1::CHERRY)] were synchronized and collected 24 hours post-L4 (∼100 worms per stage or condition) into 1.5 mL tubes. Protein samples were mixed with 5× SDS loading buffer at a 1:4 ratio and heated at 95°C for 10 minutes to denature the proteins. Worm lysates were then analyzed by Western blot using anti-Cherry (1:3000; Proteintech, 26765-1-AP) and anti-β-actin (1:3000; Proteintech, 66009-1-lg) antibodies. The %NUC-1::CHERRY cleavage=Intensity of CHERRY/(Intensity of NUC-1::CHERRY+CHERRY)/Fold change of reference (Control=1).

### Quantification of cuticle organization density and defects

To evaluate cuticle monomer density, XW18042: *qxIs722* P*_dpy-7_*DPY-7::sfGFP, single-copy insertion] worms at the same developmental stage were imaged using a Zeiss LSM900 confocal microscope. The clearest cuticle layer was selected for each worm, and images were captured at a consistent magnification (63× oil objective). Image analysis was performed using Zeiss ZEN Blue 3.4 software, applying region selection and counting tools to quantify monomer structures within a defined area. A minimum of 20 worms per strain were analyzed, with each experiment repeated three times. To assess cuticle structural defects, worms at the same developmental stage were imaged and analyzed in real-time using an Olympus BX53 upright fluorescence microscope. Images were captured at a consistent magnification (40× objective). At least 20 worms were examined per strain, and the proportion of worms displaying cuticle damage was quantified.

### Hoechst Staining Assay

Hoechst staining was performed as previously described [81]. Briefly, synchronized young adult worms were incubated in M9 buffer containing 1 mg/ml Hoechst 33342 (Wako, Japan) at room temperature for 15 minutes, followed by several washes with M9 buffer to remove residual dye. Fluorescent images were acquired under identical imaging conditions, and nuclear fluorescence in the tail and head region was quantified using ImageJ. The intestinal area was excluded to avoid interference from autofluorescence. Specifically, the head was defined as the broader, rounded anterior end of the animal and was readily recognized by the autofluorescence of the pharynx. The tail was defined as the tapered posterior end and was identified by the autofluorescence near the anal opening. All confocal images were acquired at a resolution of 1024 × 1024. For fluorescence quantification, identical ROIs (Region of Interest) (78μm × 60μm) were positioned in the head and tail regions using these anatomical landmarks and applied consistently across all animals analyzed.

### Quantification and statistical analysis

Fluorescence intensity of worms was quantified using ImageJ software. Data are presented as the mean ± SEM and were analyzed with GraphPad Prism 10. Survival curves were generated using GraphPad Prism 10, and the significant difference of two groups of samples was analyzed by Log rank (Mantel-Cox) test. Statistical analyses were performed using a Mann–Whitney’s nonparametric test for comparisons two sample groups that were not normally distributed. Statistical significance was determined as follows: */^P < 0.05, **/^^P < 0.01, ***/^^^ P < 0.001, The symbols “*” and “^” denote statistical comparisons with different groups, as specified in the figure legend. All experiments were performed independently at least three times.

## Acknowledgments

We thank the Caenorhabditis Genetics Center (CGC) (funded by NIH P40OD010440) for strains; Dr. Xiaochen Wang (SUSTech) and Chonglin Yang (Yunnan U.) for sharing strains and suggestions.

## Supporting Information captions

**S1 Fig. PA14 infection does not cause significant developmental retardation.**

**(A)** Confocal microscopy images of the vulva of animals in L4 stage. Scale bars: 10 µm.

**(B)** Confocal microscopy images of the body length of L4 animals after 24-hour OP50 or PA14 exposure. Scale bars: 100 µm.

**(C)** Confocal microscopy images of the egg morphology of L4 animals after 24-hour OP50 or PA14 exposure. Scale bars: 20 µm.

**S2 Fig. PA14-induced cuticle damage is not confined to molting cycles but occurs progressively in post-molt, intact cuticles.**

**(A)** Schematic diagram of the timelapse microscopy method for observing cuticle morphological changes in wild type (WT) animals from the L4 stage upon PA14 infection. The molting status and the DPY-7::sfGFP was observed after transferring L4 animals to OP50 or PA14 for 0, 3, 6, 12, 24 hours.

**(B)** Confocal microscopy images of the molting status and the DPY-7::sfGFP of L4 animals after 0, 3, 6, 12, 24 hours OP50 or PA14 exposure. Scale bars: 10 µm.

**(C-E)** Quantitative analysis of the worms with abnormal DPY-7::sfGFP pattern on OP50(C) and PA14(D) in different stage. And quantitative analysis of the worms with abnormal DPY-7::sfGFP pattern percentage between OP50 and PA14 in the same stage (E). n=90 animals.

For all quantification, data are presented as mean ± SEM. Statistical comparisons were performed using one-way ANOVA followed by Tukey’s multiple comparisons test (C, D). **p<0.01, ***p<0.001, n.s., not significant. All experiments were performed independently at least three times. The data underlying this figure can be found in S2 Data.

**S3 Fig. PA14 induces cuticle damage during the adult stage.**

**(A)** Schematic diagram of the timelapse microscopy method for observing cuticle morphological changes in wild type (WT) animals from L4+24h(adult) stage upon PA14 infection. DPY-7::sfGFP was observed after transferring L4+24h(adult) stage animals to OP50 or PA14 for 0, 3, 6, 12, 24 hours.

**(B)** Confocal microscopy images DPY-7::sfGFP of L4+24h(adult) animals after 0, 3, 6, 12, 24 hours OP50 or PA14 exposure. Scale bars: 10 µm.

**(C-E)** Quantitative analysis of the worms with abnormal DPY-7::sfGFP pattern on OP50(C) and PA14(D) in different stage. And quantitative analysis of the worms with abnormal DPY-7::sfGFP pattern percentage between OP50 and PA14 in the same stage(E). n=90 animals.

For all quantification, data are presented as mean ± SEM. Statistical comparisons were performed using one-way ANOVA followed by Tukey’s multiple comparisons test (C, D). *p<0.05, **p<0.01, ***p<0.001, n.s., not significant. All experiments were performed independently at least three times. The data underlying this Figure can be found in S2 Data.

**S4 Fig. PA14 induces cuticle damage during the adult stage.**

**(A)** Confocal fluorescence images and quantitative analysis of Hoechst 33342 fluorescence intensity in young adult animals after 24h PA14 exposure (infection initiated at L4 stage). RNAi knockdown of *dpy-7* and *bus-8* act as positive control. n=20 animals, Scale bars: 10 µm.

**(B)** Representative fluorescence microscopy images and quantification of P*_nlp-29_*::GFP intensity in adult wild-type worms subjected to RNAi targeting *dpy-7*, *sqt-3*, *col-142*, *dyn-1*, and *syx-5*, compared to control RNAi. n=24 animals, Scale bars: 100 µm.

For all quantification, data are presented as mean ± SEM. Statistical comparisons were performed using one-way ANOVA followed by Tukey’s multiple comparisons test. ***p<0.001,all experiments were performed independently at least three times. The data underlying this figure can be found in S2 Data.

**S5 Fig. PA14-dependent dysregulation of lysosomal architecture and cuticle integrity requires functional virulence factor expression.**

**(A)** Confocal fluorescence imaging and quantitative analysis were performed on the epidermis of wild-type (WT) expressing NUC-1::CHERRY after 24h PA14 and PA14’s key virulence regulators deficient strain Δ*gacA*, Δ*lasR*, Δ*mvfR*, Δ*pqsF* exposure. The number of tubular lysosomes was quantified by counting within 35×25(µm^2^) unit areas per individual worm, with a total of 20 worms analyzed (n=20 animals). Scale bars: 10 µm.

**(B)** Confocal fluorescence imaging and quantitative analysis of abnormal DPY-7::sfGFP pattern in wild-type (WT) L4-stage animals following 24-hour exposure to PA14 and PA14’s key virulence regulators deficient strain Δ*gacA*, Δ*lasR*, Δ*mvfR*, Δ*pqsF*. n=300 animals. Scale bars: 10 µm.

For all quantification, data are presented as mean ± SEM. Statistical comparisons were performed using one-way ANOVA followed by Tukey’s multiple comparisons test. *Indicates comparison with the OP50, ^ indicates comparison with the PA14; ***/^^^p<0.001, ns., not significant. All experiments were performed independently at least three times. The data underlying this Figure can be found in S2 Data.

**S6 Fig. Screen of PA14 induced neuronal secretion proteins which modulates lysosomal function and susceptibility to PA14.**

**(A)** List of secreted proteins induced by PA14 infection, as identified by mass spectrometry. Candidate proteins were screened for signal peptides using SignalP-5.0 and evaluated for neuronal expression patterns via CeNGEN analysis.

**(B)** Confocal imaging and quantification of NUC-1::pHTomato fluorescence in hypodermal lysosomes of wild-type animals following RNAi targeting *rla-5, dao-2, nlp-77, tag-293,* and *lec-5*. *tag-293* RNAi significantly reduced lysosomal fluorescence intensity compared to control RNAi (150 lysosomes analyzed from 30 worms). n=30 animals. Scale bars: 10 µm.

**(C)** Immunoblot detection of secreted TAG-293 in liquid culture medium after 6-hour PA14 exposure using TAG-293::3xFLAG::GFP animals.

For all quantification, data are presented as mean ± SEM. Statistical comparisons were performed using one-way ANOVA followed by Tukey’s multiple comparisons test. ***p<0.001, ns., not significant. All experiments were performed independently at least three times. The data underlying this Figure can be found in S2 Data.

**S7 Fig. NSIF-1 disrupts collagen integrity.**

**(A)** Schematic of the *nsif-1* gene and NSIF-1 protein. Exons are shown as filled boxes and introns as connecting lines. The positions of the *ok1337* deletion and the *ylf82* point mutation are indicated. The signal peptide (SP) and SHKT domains of NSIF-1 are shown.

**(B-D)** Confocal fluorescence images and quantitative analysis of BLI-1::mNeonGreen (B), CUT-2::mNeonGreen (C) and EPIC-1::GFP (D) in wild-type (WT), *nsif-1(ok1337)* and *nsif-1(ylf82)* following 24-hours PA14 exposure (infection initiated at L4 stage) or OP50. Quantification of cuticle furrows density were performed on 30 worms (n=30). And the quantitative analysis of abnormal pattern after 24h PA14 infection were performed on 100 worms with 3 times repeats (n=300 animals). Scale bar:5µm.

For all quantification, data are presented as mean ± SEM. Statistical comparisons were performed using one-way ANOVA followed by Tukey’s multiple comparisons test. **p< 0.01, ***p<0.001. All experiments were performed independently at least three times. The data underlying this figure can be found in S2 Data.

**S8 Fig. NSIF-1 enhances susceptibility to PA14**

Survival analysis of wild-type N2, *nsif-1(ok1337)* mutants, and N2 animals overexpressing full-length NSIF-1 driven by its own native promoter (P*nsif-1*) in the PA14 slow-killing assay. Data are representative of three independent biological replicates.

For all quantification, data are presented as mean ± SEM. Statistical comparisons were performed using Log rank (Mantel-Cox) test. * Indicates comparison with the wild-type, ^ indicates comparison with the *nsif-1(ok1337)* mutant. */^p<0.05, ***/^^^p<0.001; ns., not significant. All experiments were performed independently at least three times. The data underlying this figure can be found in S2 Data.

**S9 Fig. NSIF-1 expression pattern.**

(A) Confocal fluorescence imaging of transgenic animals expressing NSIF-1::GFP under the native *nsif-1* promoter (P*_nsif-1_*NSIF-1::GFP). GFP expression was observed in various neurons, muscle cells, and the intestine. Scale bars: 5µm.

(B) Confocal fluorescence imaging of the CRISPR knock-in strain NSIF-1::mNeonGreen::3×FLAG (*syb9508*), showing that the tagged protein colocalizes with the epidermis, which is marked by P*_col-12_*::mKate2. Scale bars: 100 µm (in whole animals) and 10 µm (in epidermis).

**S10 Fig. PA14-induced upregulation and lysosomal localization of NSIF-1 are specific responses to active virulence.**

**(A)** Confocal fluorescence images and quantification of NSIF-1::mNeonGreen::3×FLAG(*syb9508*) knock-in worms following 24-h exposure to wild-type PA14 or the virulence regulator-deficient mutants Δ*gacA,* Δ*lasR,* Δ*mvfR,* and Δ*pqsF*. n=30 animals. Scale bars: 10 µm.

**(B)** Confocal fluorescence imaging of NSIF-1::mNeonGreen::3×FLAG and NUC-1::CHERRY following 24-h exposure to wild-type PA14 or the PA14 virulence regulator-deficient mutants Δ*gacA,* Δ*lasR,* Δ*mvfR,* and Δ*pqsF*. Scale bars: 10 µm.

For all quantification, data are presented as mean ± SEM. Statistical comparisons were performed using one-way ANOVA followed by Tukey’s multiple comparisons test. *Indicates comparison with the OP50, ^indicates comparison with the PA14.

***/^^^p<0.001. All experiments were performed independently at least three times. The data underlying this figure can be found in S2 Data.

**S11 Fig. PA14 virulence regulators are required for neuronal NSIF-1 secretion to the epidermis.**

Confocal fluorescence images and quantification of neuronally secreted NSIF-1::GFP detected in the epidermis of transgenic animals expressing P*_rgef-1_*NSIF-1::GFP (neuronal) and P*_col-12_*GBP::mKate2 (epidermal) following 24 h of exposure to wild-type PA14 or the virulence-regulator-deficient mutants Δ*gacA,* Δ*lasR,* Δ*mvfR,* and Δ*pqsF*. n = 20 animals per condition. Scale bars: 10 µm.

For all quantification, data are presented as mean ± SEM. Statistical comparisons were performed using one-way ANOVA followed by Tukey’s multiple comparisons test. *Indicates comparison with the OP50, ^indicates comparison with the PA14.

***/^^^p<0.001. All experiments were performed independently at least three times. The data underlying this figure can be found in S2 Data.

**S12 Fig. Collagen related genes disruption does not impair NSIF-1 secretion to the epidermis.**

Confocal fluorescence images and quantification of epidermal NSIF-1::GFP fluorescence in P*_rgef-1_*NSIF-1::GFP;P*_col-12_*GBP::mKate2 worms following *col-124, dpy-7, dyn-1, sqt-3,* or *syx-5* RNAi. No significant difference in epidermal NSIF-1::GFP fluorescence was observed compared with the control RNAi group. n=30 animals. Scale bars: 10 µm.

For all quantification, data are presented as mean ± SEM. Statistical comparisons were performed using one-way ANOVA followed by Tukey’s multiple comparisons test. ns., not significant. All experiments were performed independently at least three times. The data underlying this figure can be found in S2 Data.

**S13 Fig. NSIF-1 release occurs through a non-canonical pathway independent of traditional neuropeptide processing mechanisms.**

Confocal fluorescence images and quantification of epidermal NSIF-1::GFP fluorescence in P*_rgef-1_*NSIF-1::GFP;P*_col-12_*GBP::mKate2 worms following 24-h exposure to OP50 or PA14 with *unc-31* or *egl-3* RNAi. Neither *unc-31* nor *egl-3* RNAi significantly affected epidermal NSIF-1::GFP fluorescence. n=30 animals. Scale bars: 10 µm.

For all quantification, data are presented as mean ± SEM. Statistical comparisons were performed using one-way ANOVA followed by Tukey’s multiple comparisons test. ns., not significant. All experiments were performed independently at least three times. The data underlying this figure can be found in S2 Data.

**S14 Fig. Co-localization analysis of NSIF-1 transcriptional and translational reporters with ASI, ASJ, ADF, and RIS neurons.**

**(A)** Localization of the NSIF-1 transcriptional reporter (P*_nsif-1_*::GFP) in ASI (P*_str-3_*::mKate2), ASJ (P*_trx-1_*::mKate2), ADF (P*_tph-1_*::mKate2), and RIS (P*_ggr-2_*::mKate2) neurons. Scale bars: whole animal, 100 µm; others, 20 µm.

**(B)** Localization of the NSIF-1 transcriptional reporter (P*_nsif-1_*NSIF-1::GFP) in ASI (P*_str-3_*::mKate2), ASJ (P*_trx-1_*::mKate2), ADF (P*_tph-1_*::mKate2), and RIS (P*_ggr-2_*::mKate2) neurons. Scale bars: whole animal, 100 µm; others, 10 µm.

**S15 Fig. Loss of *nsif-1* upregulates lysosome-associated gene expression.**

**(A)** KEGG pathway analysis of differentially upregulated genes in *nsif-1(ok1337)* relative to wild-type (WT) from RNA-seq. Upregulated genes for KEGG pathway analysis were defined as those with a false discovery rate (FDR) < 0.05 and a fold change≥1.5.

**(B)** Functional categorization of 54 lysosomal genes upregulated in *nsif-1(ok1337)* (identified in A) from RNA-seq.

**(C)** qRT-PCR validation of lysosomal gene expression changes in WT and adult *nsif-1(ok1337)*, grouped by functional class: membrane proteins, lipid hydrolases, proteases, non-proteases, and v-ATPase components.

For all quantification, data are presented as mean±SEM. Statistical comparisons were performed using Mann–Whitney U-test. *p<0.05, **p<0.01, ***p<0.001; ns., not significant. All experiments were performed independently at least three times. The data underlying this figure can be found in S2 Data.

**S16 Fig. Functional analysis of multi-tissue rescue of NSIF-1 in regulating lysosomal homeostasis and cuticle integrity.**

(A) Confocal fluorescence imaging and quantitative analysis of DPY-7::sfGFP marked cuticle in wild-type(WT), *nsfi-1(ok1337)*, *nsfi-1(ok1337)*;P*_myo-3_*NSIF-1::mKate2, *nsfi-1(ok1337)*;P*_vha-6_*NSIF-1::mKate2 animals following 24-h PA14 exposure (infection initiated at the L4 stage). Quantification of abnormal DPY-7::sfGFP pattern performed on 300 worms (n=300 animals). Scale bars: 10 µm.

(B) Confocal fluorescence imaging and quantitative analysis of NUC-1::CHERRY marked lysosome in wild-type(WT), *nsfi-1(ok1337)*, *nsfi-1(ok1337)*;P*_myo-3_*NSIF-1::mKate2, *nsfi-1(ok1337)*;P*_vha-6_*NSIF-1::mKate2 animals following 24-h PA14 exposure (infection initiated at the L4 stage). Quantification of tubular lysosomes were quantified by counting three 35 × 25 µm² regions per worm (n=20 worms). Scale bars: 10 µm.

(C) Confocal fluorescence imaging and quantitative analysis of DPY-7::sfGFP marked cuticle in wild-type(WT), *nsfi-1(ok1337)*, *nsfi-1(ok1337)*;P*_col-12_*NSIF-1::mKate2, *nsfi-1(ok1337)*;P*_col-12_*ΔSPNSIF-1::mKate2 animals following 24-h PA14 exposure (infection initiated at the L4 stage). Quantification of abnormal DPY-7::sfGFP pattern performed on 300 worms (n=300 animals). Scale bars: 10 µm.

(D) Confocal fluorescence imaging and quantitative analysis of NUC-1::CHERRY marked lysosome in wild-type(WT), *nsfi-1(ok1337)*, *nsfi-1(ok1337)*;P*_col-12_*NSIF-1::mKate2, *nsfi-1(ok1337)*;P*_col-12_*ΔSPNSIF-1::mKate2 animals following 24-h PA14 exposure (infection initiated at the L4 stage). Quantification of tubular lysosomes were quantified by counting three 35 × 25 µm² regions per worm (n = 20 worms). Scale bars: 10 µm.

For all quantification, data are presented as mean ± SEM. Statistical comparisons were performed using one-way ANOVA followed by Tukey’s multiple comparisons test. *Indicates comparison with the wild-type, ^ indicates comparison with the *nsif-1(ok1337)* mutant; **/^^ p < 0.01, ***/^^^ p< 0.001; ns., not significant. All experiments were performed independently at least three times. The data underlying this figure can be found in S2 Data.

**S17 Fig. Neuronal NSIF-1 in ASI/ASJ neurons is sufficient to impact lysosomal homeostasis and cuticle integrity.**

**(A)** Confocal fluorescence images and quantitative analysis of NUC-1::CHERRY-labeled lysosomes in wild-type (WT), *nsif-1(ok1337)* mutants, and *nsif-1(ok1337)* mutants expressing full-length NSIF-1::GFP or the signal-peptide-deleted variant (ΔSP-NSIF-1::GFP) under the control of the endogenous *nsif-1* promoter (P*nsif-1*), the ASI promoter (P_ASI_), or the ASJ promoter (P_ASJ_) following 24 h of PA14 exposure (initiated at the L4 larval stage). Quantification of tubular lysosomes were quantified by counting three 35 × 25 µm² regions per worm (n=20 worms). Scale bars: 10 µm.

**(B)** Confocal fluorescence imaging and quantitative analysis of DPY-7::sfGFP marked cuticle in wild-type (WT), *nsif-1(ok1337)* mutants, and *nsif-1(ok1337)* mutants expressing full-length NSIF-1::GFP or the signal-peptide-deleted variant (ΔSP-NSIF-1::GFP) under the control of the endogenous *nsif-1* promoter (P*nsif-1*), the ASI promoter (P_ASI_), or the ASJ promoter (P_ASJ_) following 24 h of PA14 exposure (initiated at the L4 larval stage). Quantification of abnormal DPY-7::sfGFP pattern performed on 300 worms(n=300). Scale bars: 10 µm.

Note: the data shown in (A-B) for the wild-type (WT), *nsif-1(ok1337)* mutants, and *nsif-1(ok1337)* mutants expressing full-length NSIF-1::GFP (driven by the endogenous *nsif-1* promoter /P*nsif-1*, the ASI promoter /P_ASI_, or the ASJ promoter/P_ASJ_) are the same as those presented in Figure 6C-6D. The S17 Figure extends these results by showing the corresponding data for *nsif-1(ok1337)* mutants expressing the signal-peptide-deleted variant (ΔSP-NSIF-1::GFP) under the control of the endogenous *nsif-1* promoter (P*nsif-1*), the ASI promoter (P_ASI_), or the ASJ promoter (P_ASJ_)

For all quantification, data are presented as mean ± SEM. Statistical comparisons were performed using one-way ANOVA followed by Tukey’s multiple comparisons test. *Indicates comparison with the wild-type, ^ indicates comparison with the nsif-1(ok1337) mutant; **/^^ p < 0.01, ***/^^^ p< 0.001; ns., not significant. All experiments were performed independently at least three times. The data underlying this figure can be found in S2 Data.

**S18 Fig. ASI/ASJ neuron ablation reduces NSIF-1 secretion to the epidermis and preserves lysosomal homeostasis during PA14 infection.**

(A) Confocal microscopy images and quantification of ASI/ASJ neuron ablation efficiency and epidermal lysosomal morphology. EGL-1-mediated genetic ablation of ASI and ASJ neurons resulted in a significant reduction of NSIF-1::GFP fluorescence in these neurons. Following 24 h of PA14 exposure (initiated at the L4 larval stage), ablation of ASI/ASJ neurons prevented the PA14-induced reduction in tubular lysosomes. NSIF-1::GFP fluorescence intensity was quantified in 30 worms. Tubular lysosomes were quantified by counting three 35 × 25 µm² regions per worm (n=20 worms).

(B) Confocal microscopy images and quantification of epidermally captured NSIF-1::GFP in WT, ASI-ablated, or ASJ-ablated worms following 24-h PA14 exposure in transgenic animals expressing P*_rgef-1_*NSIF-1::GFP+P*_col-12_*GBP::mKate2 (infection initiated at the L4 stage) Compared with control worms, genetic ablation of either ASI or ASJ neurons significantly reduced the epidermal NSIF-1::GFP signal (n=30 animals). Scale bars: 10 µm.

For all quantification, data are presented as mean ± SEM. Statistical comparisons were performed using Mann–Whitney U-test(A) and one-way ANOVA followed by Tukey’s multiple comparisons test(B). ***p<0.001, ns., not significant. All experiments were performed independently at least three times. The data underlying this figure can be found in S2 Data.

**S19 Fig. ASI/ASJ neuron ablation enhances epidermal lysosomal function.**

**(A)** Confocal fluorescence imaging of NUC-1::CHERRY in wild-type (WT), *nsfi-1(ok1337)*, and animals with ASI and ASJ neurons ablated, following 24 h of PA14 exposure (initiated at the L4 larval stage). Scale bars: 10 µm.

**(B)** Quantitative analysis of the number of tubular lysosomes in wild-type (WT), n*sfi-1(ok1337)*, and ASI/ASJ-ablated animals. The number of tubular lysosomes was quantified by counting within three 35×25(µm^2^) unit areas per worm (n=20 animals).

For all quantification, data are presented as mean ± SEM. Statistical comparisons were performed using one-way ANOVA followed by Tukey’s multiple comparisons test. *p<0.05, **p<0.01, ***p<0.001, ns., not significant. All experiments were performed independently at least three times. The data underlying this figure can be found in S2 Data.

**S20 Fig. Epidermal expression of NSIF-1 is sufficient to impact lysosomal homeostasis and cuticle integrity.**

**(A)** Confocal fluorescence imaging and quantification of NUC-1:: CHERRY-labeled lysosomes in wild-type (WT), *nsif-1(ok1337)* mutants, and WT animals expressing NSIF-1::mKate2 or the signal-peptide-deleted variant (ΔSP-NSIF-1::mKate2) under the control of the endogenous *nsif-1* promoter (P*nsif-1*) or the epidermal promoter (P*col-12*). Quantification was performed on 20 animals (n=20). Scale bars: 10 µm.

**(B)** Confocal fluorescence imaging and quantification of DPY-7::sfGFP-marked cuticle in WT, *nsif-1(ok1337)* mutants, and WT animals expressing NSIF-1::mKate2 or the signal-peptide-deleted variant (ΔSP-NSIF-1::mKate2) under the control of the endogenous *nsif-1* promoter (P*nsif-1*) or the epidermal promoter (P*col-12*). Quantification was performed on 300 animals (n=300). Scale bars: 10 µm.

For all quantification, data are presented as mean ± SEM. Statistical comparisons were performed using one-way ANOVA followed by Tukey’s multiple comparisons test. ***p<0.001, ns., not significant. All experiments were performed independently at least three times. The data underlying this figure can be found in S2 Data.

**S21 Fig. Identification of a putative NLS sequence in NSIF-1 using cNLS Mapper.** The online tool (https://nls-mapper.iab.keio.ac.jp/) predicted a candidate sequence with a score of 3.3.

**S22 Fig. ELT-3 functions downstream of NSIF-1 to regulate epidermal lysosomal function.**

**(A)** Confocal fluorescence imaging of localization of ELT-3::GFP and P*_col-12_*ΔSPNSIF-1::mKate2::3×HA in adult animals. Scale bars, 10µm.

**(B)** Transcription factor binding analysis identified ∼200 ELT-3-regulated genes among transcripts upregulated in *nsif-1* mutants.

**(C)** qRT-PCR analysis of *elt-3* expression changes in adult (L4+24h) WT and *nsif-1(ok1337)* animals. Three biological replicates, each with three technical replicates, were analyzed for *elt-3*.

**(D-E)** Analysis of hypodermal lysosomes in WT, *nsif-1(ok1337), elt-3(vp1),* and *nsif-1(ok1337);elt-3(vp1)* animals expressing NUC-1:: CHERRY following 24 h of PA14 exposure. The number of vesicular lysosomes (D) was quantified within three 35 × 25 µm² regions per worm (n = 20 animals). The diameter of vesicular lysosomes

**(E)** was determined by measuring the six largest vesicles within a 35 × 25 µm² region per worm (n = 10 animals)

**(F)** Immunoblots and quantification showing cleavage of CHERRY from NUC-1::CHERRY expressed in the epidermis of adult wild type(WT), *nsif-1(ok1337), elt-3(gk121), nsif-1(ok1337);elt-3(vp1)*. N=3 biological replicates.

For all quantification, data are presented as mean ± SEM. Statistical comparisons were performed using Mann–Whitney U-test (C) and one-way ANOVA followed by Tukey’s multiple comparisons test (D-F). *Indicates comparison with the wild-type, ^ indicates comparison with the *nsif-1(ok1337)* mutant; */^p< 0.05, **/^^p< 0.01, ***/^^^p< 0.001; ns., not significant. All experiments were performed independently at least three times. The data underlying this figure can be found in S2 Data.

## Legends for Supplemental Table

**S1 Table.** The secreted proteins identified by mass spectrometry.

**S2 Table.** Differentially expressed genes (DEGs) in *nsif-1(ok1337)* mutants versus wild-type (RNA-seq analysis).

**S3 Table.** Gene enrichment results corresponding to S15A-B Fig, S22B Fig.

**S4 Table.** Oligonucleotide primers used in this study.

**S1 Raw Images.** Data supplement for Western Blot Full Gel Images.

## Notes

### Competing Interest Statement

The authors have declared no competing interest.

### Summary of Updates

In this revision, we add supplemenary fiures (S1-S22 Figures), and updated Figure 1, Figure 3, Figure 4, Figure 5, Figure 6, Figure 7 and Figure 8.

