## Supplemental Figures for "Pathogen Subversion of Neuro-Epidermal Signaling Impairs Lysosomal Function to Disrupt Collagen Homeostasis"

#### Table of contents

| Supplemental Figure Name | Page Number |
|---|---|
| **S1 Fig.** | 2 |
| **S2 Fig.** | 3-4 |
| **S3 Fig.** | 5-6 |
| **S4 Fig.** | 7-8 |
| **S5 Fig.** | 9 |
| **S6 Fig.** | 10-11 |
| **S7 Fig.** | 12-13 |
| **S8 Fig.** | 14 |
| **S9 Fig.** | 15 |
| **S10 Fig.** | 16-17 |
| **S11 Fig.** | 18 |
| **S12 Fig.** | 19 |
| **S13 Fig.** | 20 |
| **S14 Fig.** | 21 |
| **S15 Fig.** | 22 |
| **S16 Fig.** | 23-24 |
| **S17 Fig.** | 25-26 |
| **S18 Fig.** | 27-28 |
| **S19 Fig.** | 29 |
| **S20 Fig.** | 30 |
| **S21 Fig.** | 31 |
| **S22 Fig.** | 32 |

### S1 Fig.

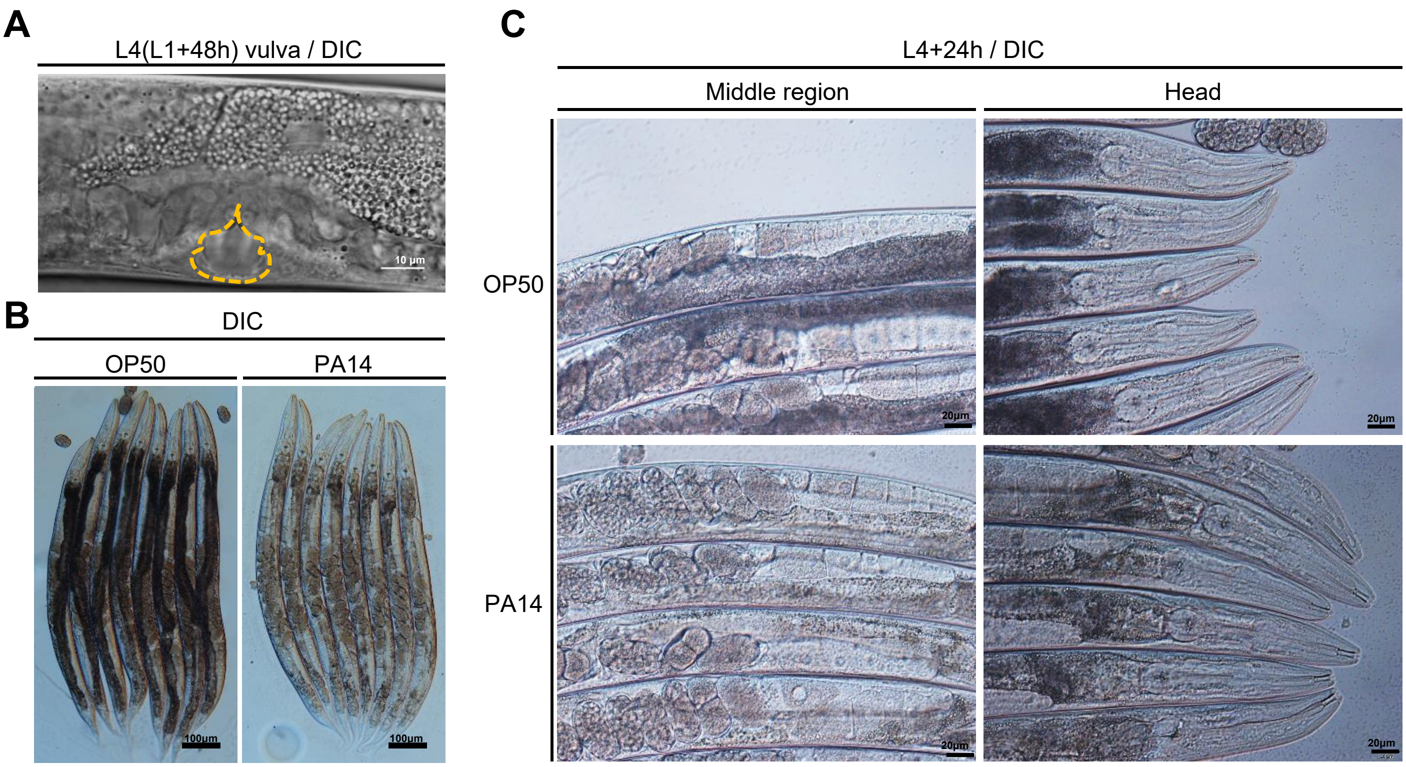

**S1 Fig. PA14 infection does not cause significant developmental retardation.**
**(A)** Confocal microscopy images of the vulva of animals in L4 stage. Scale bars: 10 μm.
**(B)** Confocal microscopy images of the body length of L4 animals after 24-hour OP50 or PA14 exposure. Scale bars: 100 μm.
**(C)** Confocal microscopy images of the egg morphology of L4 animals after 24-hour OP50 or PA14 exposure. Scale bars: 20 μm.

### S2 Fig.

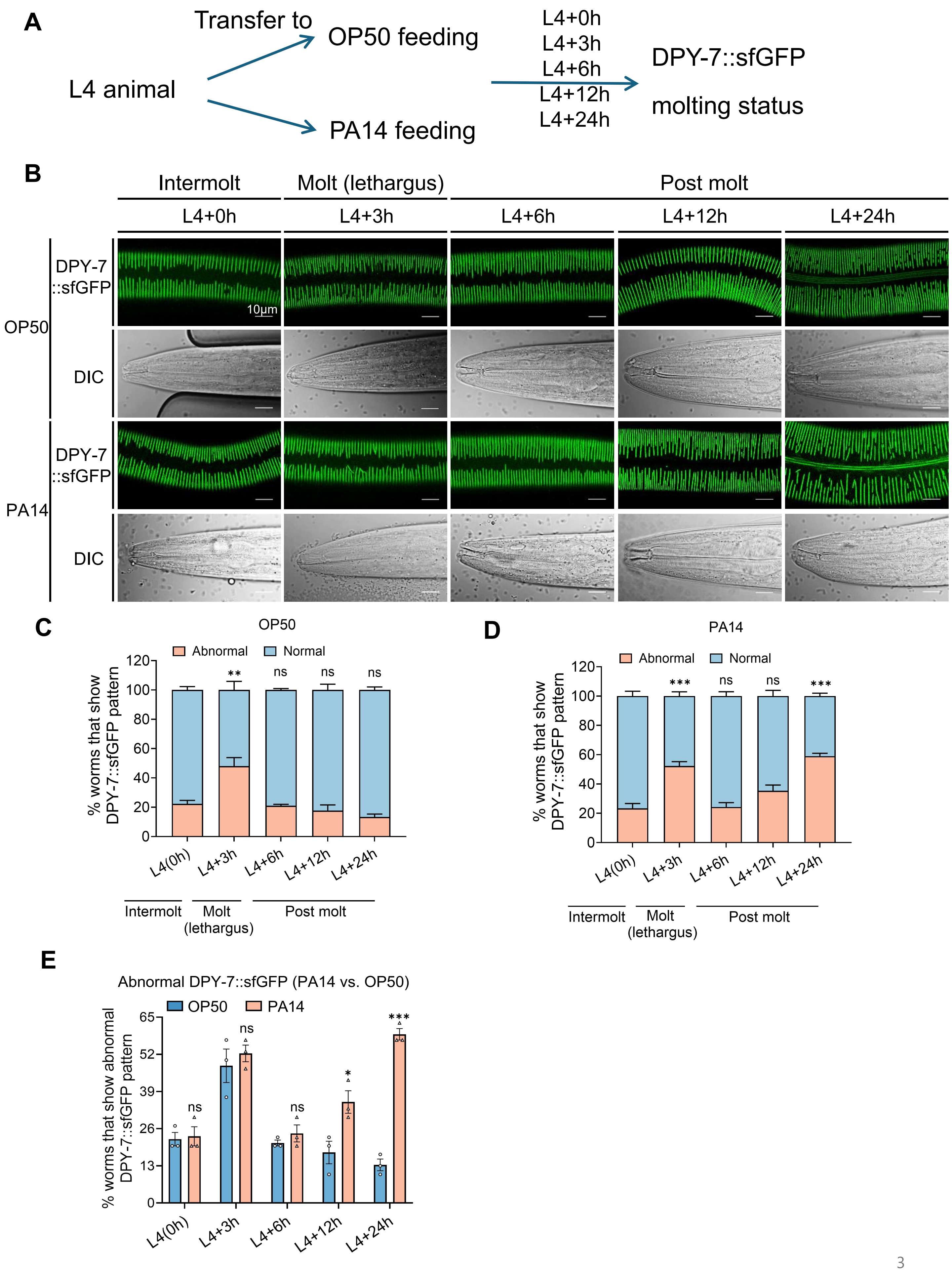
A
Transfer to
OP50 feeding
L4 animal
PA14 feeding
L4+0h
L4+3h
L4+6h
L4+12h
L4+24h
DPY-7::sfGFP
molting status
B
Intermolt
Molt (lethargus)
Post molt
L4+0h
L4+3h
L4+6h
L4+12h
L4+24h
OP50
DPY-7 ::sfGFP
DIC
PA14
DPY-7 ::sfGFP
DIC
10μm
C
OP50
Abnormal
Normal
% worms that show DPY-7::sfGFP pattern
**
ns
ns
ns
L4(0h)
L4+3h
L4+6h
L4+12h
L4+24h
Intermolt
Molt (lethargus)
Post molt
D
PA14
Abnormal
Normal
% worms that show DPY-7::sfGFP pattern
***
ns
ns
***
L4(0h)
L4+3h
L4+6h
L4+12h
L4+24h
Intermolt
Molt (lethargus)
Post molt
E
Abnormal DPY-7::sfGFP (PA14 vs. OP50)
OP50
PA14
% worms that show abnormal DPY-7::sfGFP pattern
ns
ns
ns
*
***
L4(0h)
L4+3h
L4+6h
L4+12h
L4+24h

**S2 Fig. PA14-induced cuticle damage is not confined to molting cycles but occurs progressively in post-molt, intact cuticles.**

**(A)** Schematic diagram of the timelapse microscopy method for observing cuticle morphological changes in wild type (WT) animals from the L4 stage upon PA14 infection. The molting status and the DPY-7::sfGFP was observed after transferring L4 animals to OP50 or PA14 for 0, 3, 6, 12, 24 hours.

**(B)** Confocal microscopy images of the molting status and the DPY-7::sfGFP of L4 animals after 0, 3, 6, 12, 24 hours OP50 or PA14 exposure. Scale bars: 10 µm.

**(C-E)** Quantitative analysis of the worms with abnormal DPY-7::sfGFP pattern on OP50(C) and PA14(D) in different stage. And quantitative analysis of the worms with abnormal DPY-7::sfGFP pattern percentage between OP50 and PA14 in the same stage (E). n=90 animals.

For all quantification, data are presented as mean ± SEM. Statistical comparisons were performed using one-way ANOVA followed by Tukey's multiple comparisons test (C, D). **p<0.01, ***p<0.001, n.s., not significant. All experiments were performed independently at least three times. The data underlying this figure can be found in S2 Data.

### S3 Fig.

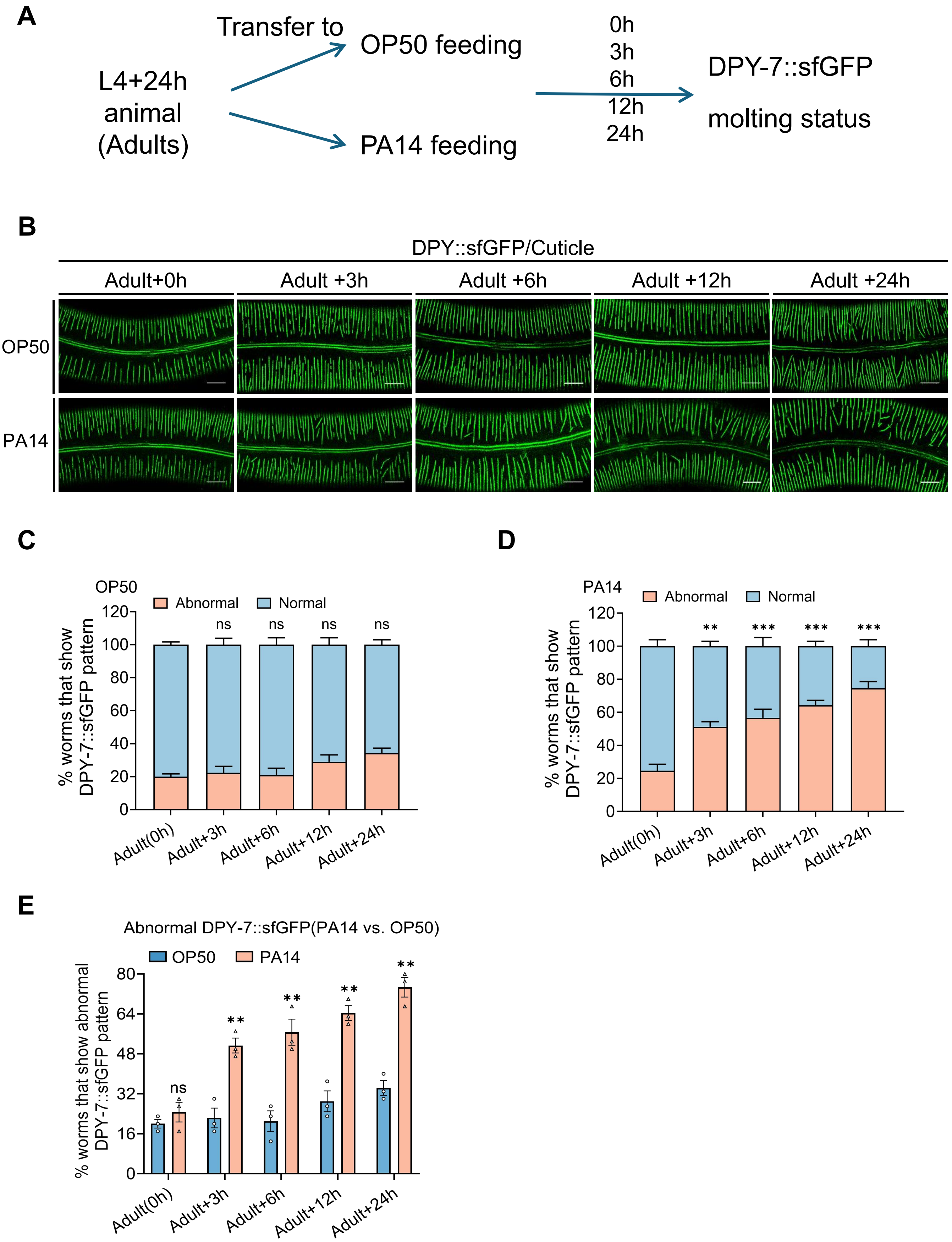
A
L4+24h animal (Adults)
Transfer to
OP50 feeding
PA14 feeding
0h
3h
6h
12h
24h
DPY-7::sfGFP
molting status
B
DPY::sfGFP/Cuticle
Adult+0h
Adult +3h
Adult +6h
Adult +12h
Adult +24h
OP50
PA14
C
OP50
Abnormal
Normal
% worms that show DPY-7::sfGFP pattern
ns
Adult(0h)
Adult+3h
Adult+6h
Adult+12h
Adult+24h
D
PA14
Abnormal
Normal
% worms that show DPY-7::sfGFP pattern
**
***
Adult(0h)
Adult+3h
Adult+6h
Adult+12h
Adult+24h
E
Abnormal DPY-7::sfGFP(PA14 vs. OP50)
OP50
PA14
% worms that show abnormal DPY-7::sfGFP pattern
ns
**
Adult(0h)
Adult+3h
Adult+6h
Adult+12h
Adult+24h

**S3 Fig. PA14 induces cuticle damage during the adult stage.**

**(A)** Schematic diagram of the timelapse microscopy method for observing cuticle morphological changes in wild type (WT) animals from L4+24h(adult) stage upon PA14 infection. DPY-7::sfGFP was observed after transferring L4+24h(adult) stage animals to OP50 or PA14 for 0, 3, 6, 12, 24 hours.

**(B)** Confocal microscopy images DPY-7::sfGFP of L4+24h(adult) animals after 0, 3, 6, 12, 24 hours OP50 or PA14 exposure. Scale bars: 10 µm.

**(C-E)** Quantitative analysis of the worms with abnormal DPY-7::sfGFP pattern on OP50(C) and PA14(D) in different stage. And quantitative analysis of the worms with abnormal DPY-7::sfGFP pattern percentage between OP50 and PA14 in the same stage(E). n=90 animals.

For all quantification, data are presented as mean ± SEM. Statistical comparisons were performed using one-way ANOVA followed by Tukey's multiple comparisons test (C, D). *p<0.05, **p<0.01, ***p<0.001, n.s., not significant. All experiments were performed independently at least three times. The data underlying this Figure can be found in S2 Data.

### S4 Fig.

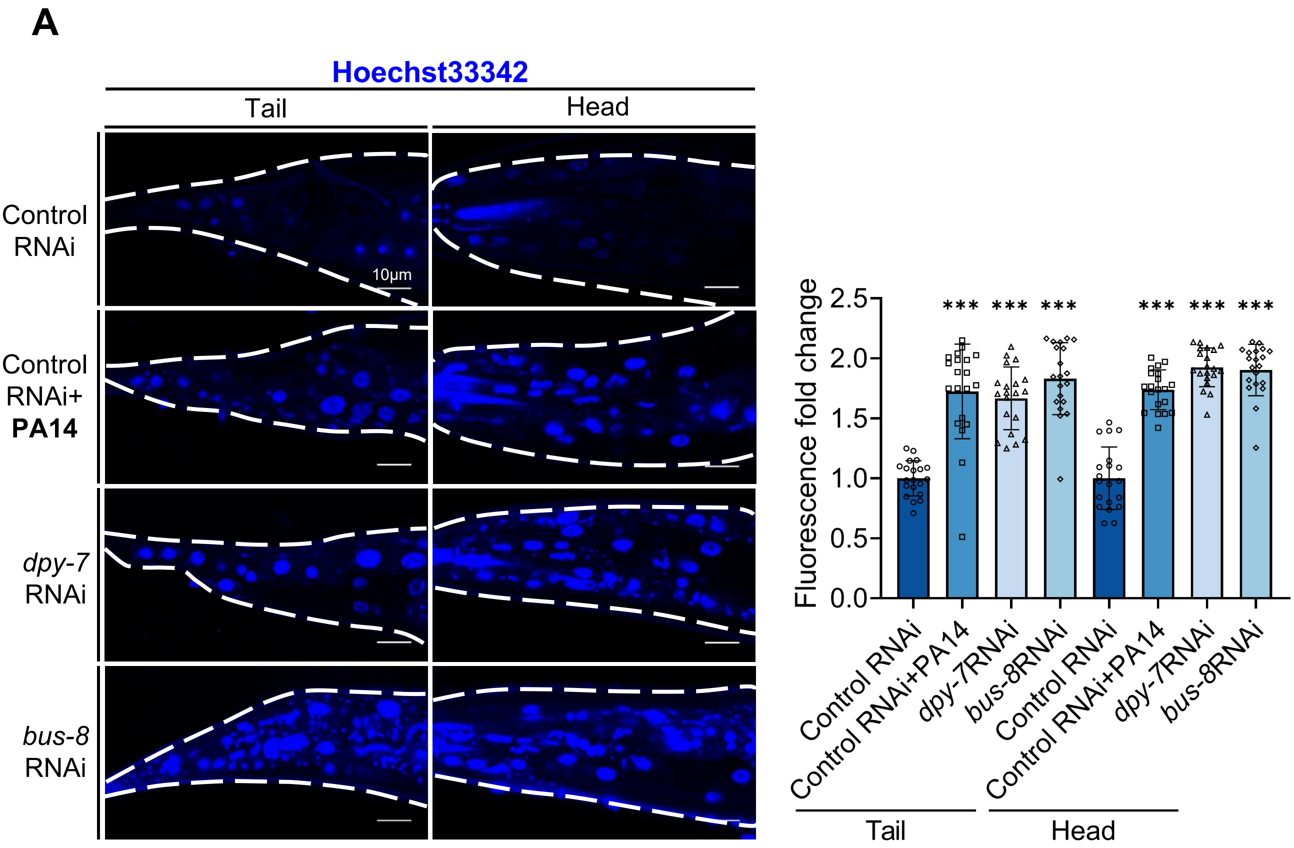

A
Hoechst33342
Tail
Head
Control RNAi
Control RNAi+ PA14
dpy-7 RNAi
bus-8 RNAi
10μm
Fluorescence fold change
2.5
2.0
1.5
1.0
0.5
0.0
***
Control RNAi
Control RNAi+PA14
dpy-7RNAi
bus-8RNAi
Tail
Head

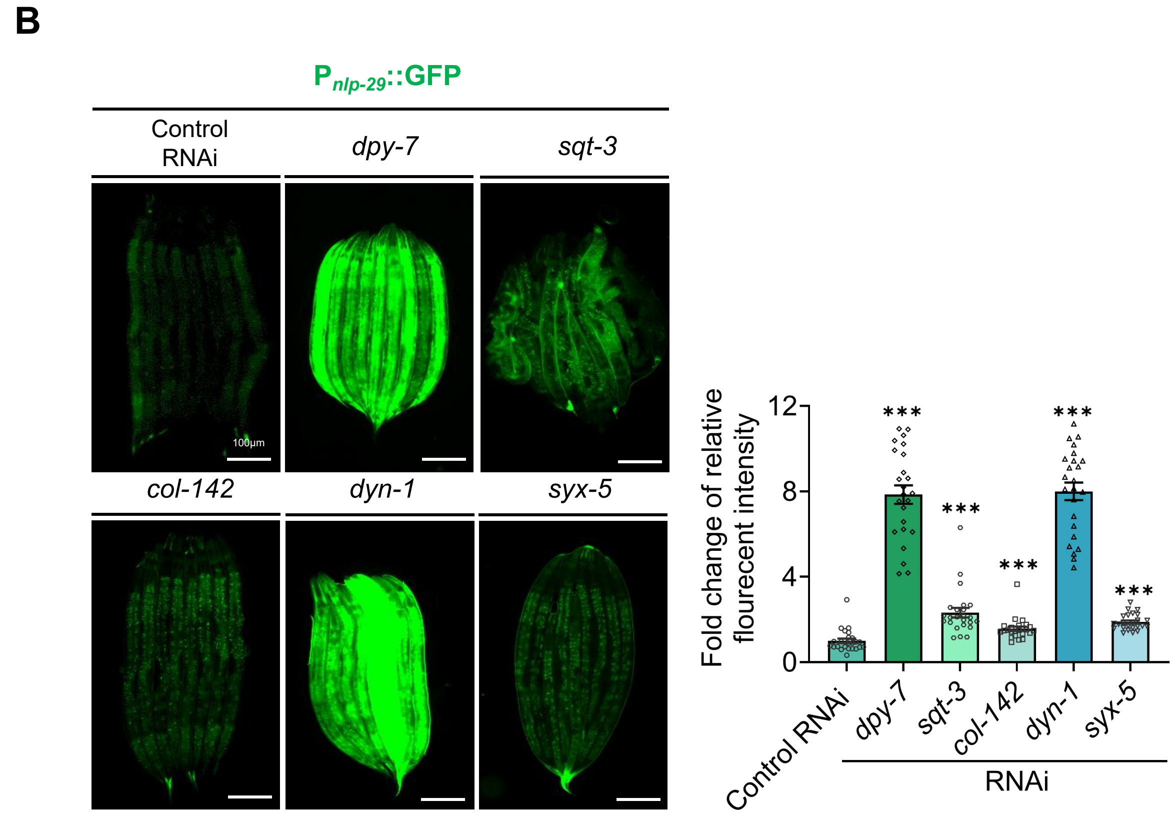

B
$P_{nlp-29}$::GFP
Control RNAi
dpy-7
sqt-3
col-142
dyn-1
syx-5
100μm
Fold change of relative flourecent intensity
12
8
4
0
***
Control RNAi
RNAi

**S4 Fig. PA14 induces cuticle damage during the adult stage.**

**(A)** Confocal fluorescence images and quantitative analysis of Hoechst 33342 fluorescence intensity in young adult animals after 24h PA14 exposure (infection initiated at L4 stage). RNAi knockdown of *dpy-7* and *bus-8* act as positive control. n=20 animals, Scale bars: 10 µm.

**(B)** Representative fluorescence microscopy images and quantification of $P_{nlp\text{-}29}$::GFP intensity in adult wild-type worms subjected to RNAi targeting *dpy-7*, *sqt-3*, *col-142*, *dyn-1*, and *syx-5*, compared to control RNAi. n=24 animals, Scale bars: 100 µm.

For all quantification, data are presented as mean ± SEM. Statistical comparisons were performed using one-way ANOVA followed by Tukey's multiple comparisons test. ***p<0.001,all experiments were performed independently at least three times. The data underlying this figure can be found in S2 Data.

### S5 Fig.

**A**

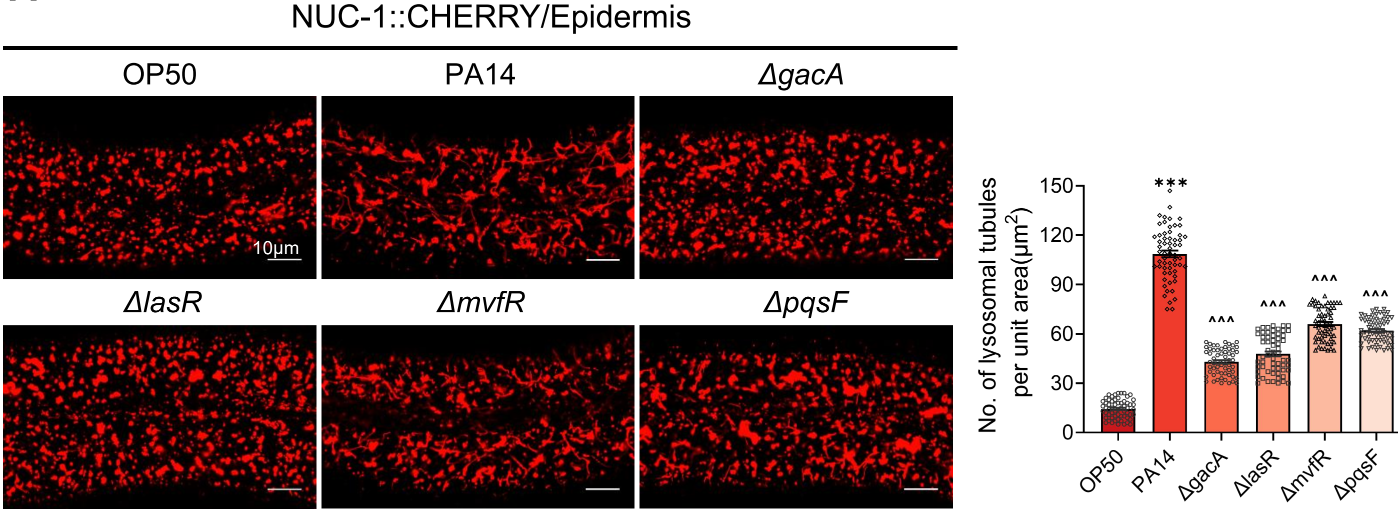

**B**

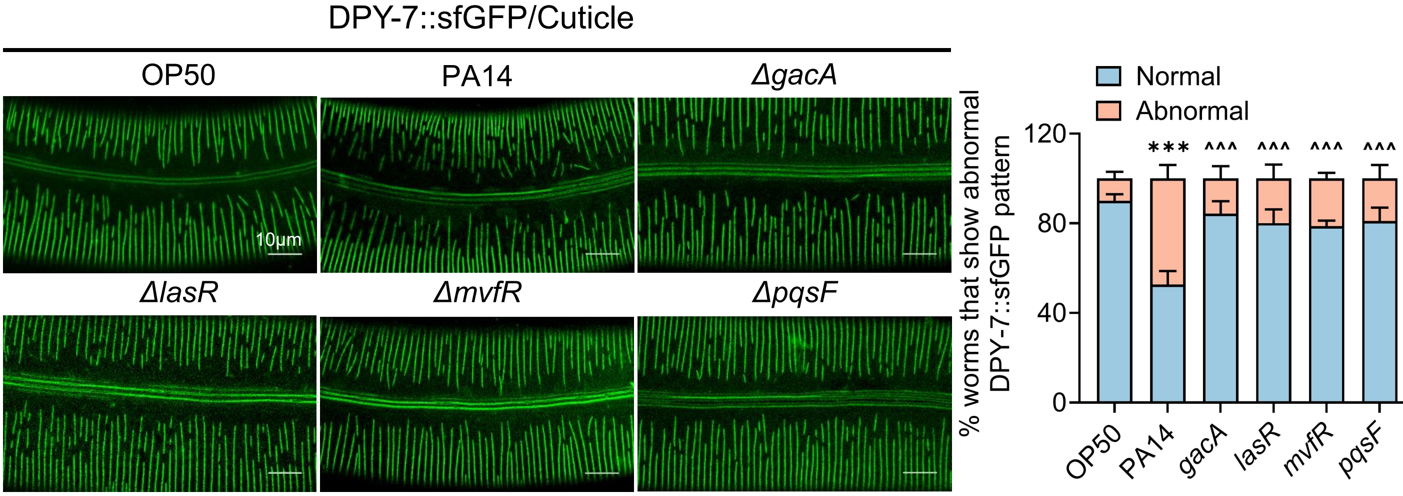

**S5 Fig. PA14-dependent dysregulation of lysosomal architecture and cuticle integrity requires functional virulence factor expression.**

**(A)** Confocal fluorescence imaging and quantitative analysis were performed on the epidermis of wild-type (WT) expressing NUC-1::CHERRY after 24h PA14 and PA14's key virulence regulators deficient strain *ΔgacA*, *ΔlasR*, *ΔmvfR*, *ΔpqsF* exposure. The number of tubular lysosomes was quantified by counting within 35×25(μm$^2$) unit areas per individual worm, with a total of 20 worms analyzed (n=20 animals). Scale bars: 10 μm.

**(B)** Confocal fluorescence imaging and quantitative analysis of abnormal DPY-7::sfGFP pattern in wild-type (WT) L4-stage animals following 24-hour exposure to PA14 and PA14's key virulence regulators deficient strain *ΔgacA*, *ΔlasR*, *ΔmvfR*, *ΔpqsF* . n=300 animals. Scale bars: 10 μm.

For all quantification, data are presented as mean ± SEM. Statistical comparisons were performed using one-way ANOVA followed by Tukey's multiple comparisons test. *Indicates comparison with the OP50, ^ indicates comparison with the PA14; ***/^^^p<0.001, ns., not significant. All experiments were performed independently at least three times. The data underlying this Figure can be found in S2 Data.

### S6 Fig.

**A**

| Gene names | Mol. weight [kDa] | Peptides | Signal Peptide (Yea r No) | Neuron |
|---|---|---|---|---|
| *rla-2* | 10.871 | 2 | Yes | RIG,ADA,NSM,RIH |
| *par-5* | 14.267 | 3 | No | - |
| *lbp-9* | 15.263 | 2 | No | - |
| *dao-2* | 15.298 | 5 | Yes | PHA,AQR,AVB,AVD,AFD |
| *lec-9* | 15.556 | 2 | No | - |
| *nlp-77* | 15.86 | 7 | Yes | AFD,RMG,NSM,RID |
| *rps-10* | 16.87 | 4 | No | - |
| *iff-1* | 17.867 | 3 | No | - |
| *icd-2* | 20.033 | 5 | No | - |
| *far-3* | 20.905 | 4 | Yes | - |
| *prdx-2* | 21.785 | 6 | No | - |
| *tag-293* | 26.156 | 4 | Yes | ASJ |
| *aldo-2* | 28.015 | 6 | No | - |
| *C31C9.2* | 34.691 | 7 | No | - |
| *lec-5* | 35.437 | 4 | Yes | DVC,PHB,AWC-OFF,AWB |
| *nap-1* | 35.664 | 6 | No | - |
| *nex-1* | 35.695 | 7 | No | - |
| *sgt-1* | 36.467 | 2 | No | - |
| *F35E12.6* | 38.736 | 3 | Yes | - |
| *pdi-2* | 41.619 | 3 | No | SIA,RIB,RMF,SIB,AVB |
| *clec-65* | 41.626 | 7 | Yes | - |
| *idh-1* | 45.959 | 6 | No | AVG,RMD |
| *cct-6* | 47.67 | 3 | No | ALM,PLM,PDE,PVN |
| *C49C8.5* | 48.68 | 3 | Yes | - |
| *gdi-1* | 50.024 | 3 | No | AVG,PVR,HSN,AVJ |
| *pars-1* | 51.586 | 2 | No | SIA,RIB,SIB,AVG |
| *spe-5* | 56.034 | 1 | No | URB,RMH,PDA RMF |
| *cct-7* | 58.372 | 5 | No | PDE,SDQ,PLM |
| *asns-2* | 61.736 | 2 | No | AVD,ADL,ASJ,PVQ |
| *bgal-1* | 74.63 | 3 | No | RIP,RIR,AVH |
| *noah-2* | 81.738 | 8 | Yes | - |
| *R08F11.7* | 83.991 | 4 | Yes | - |
| *noah-1* | 115.32 | 14 | Yes | - |
| *vit-2* | 180.92 | 28 | Yes | - |
| *fasn-1* | 289.22 | 6 | No | - |

**B**

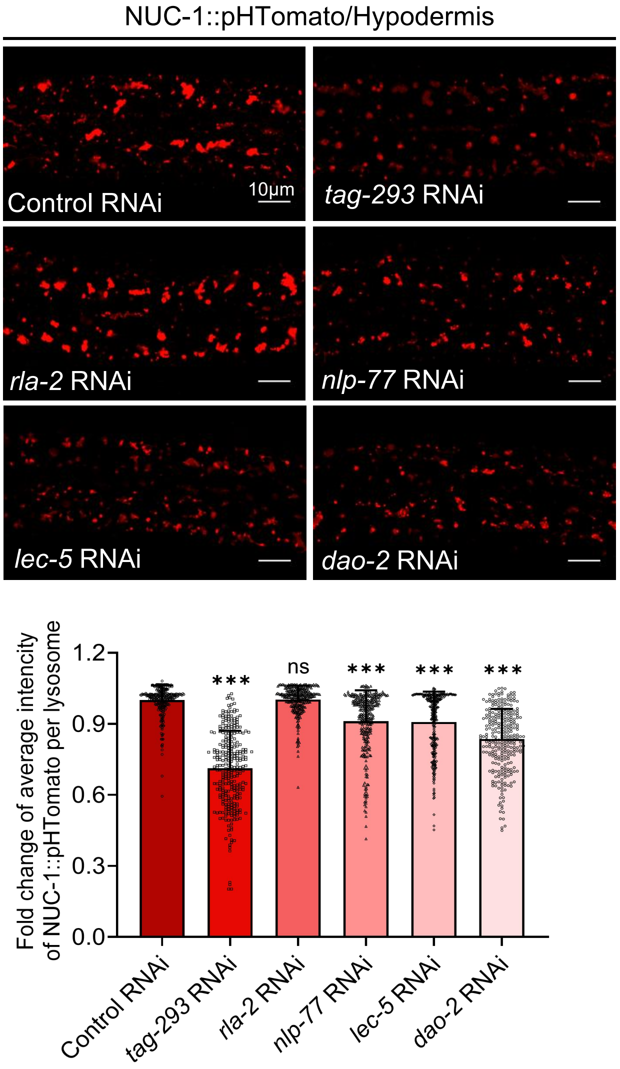

**C**

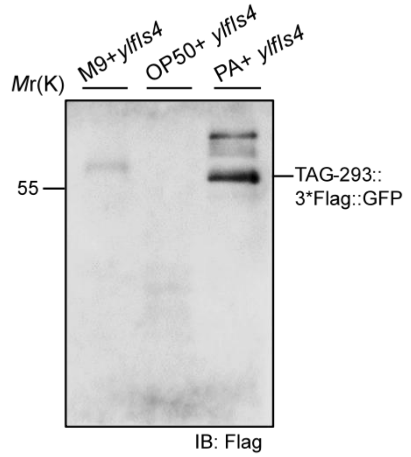

**S6 Fig. Screen of PA14 induced neuronal secretion proteins which modulates lysosomal function and susceptibility to PA14.**

**(A)** List of secreted proteins induced by PA14 infection, as identified by mass spectrometry. Candidate proteins were screened for signal peptides using SignalP-5.0 and evaluated for neuronal expression patterns via CeNGEN analysis.

**(B)** Confocal imaging and quantification of NUC-1::pHTomato fluorescence in hypodermal lysosomes of wild-type animals following RNAi targeting *rla-5, dao-2, nlp-77, tag-293,* and *lec-5*. *tag-293* RNAi significantly reduced lysosomal fluorescence intensity compared to control RNAi (150 lysosomes analyzed from 30 worms). n=30 animals. Scale bars: 10 μm.

**(C)** Immunoblot detection of secreted TAG-293 in liquid culture medium after 6-hour PA14 exposure using TAG-293::3xFLAG::GFP animals.

For all quantification, data are presented as mean ± SEM. Statistical comparisons were performed using one-way ANOVA followed by Tukey's multiple comparisons test. ***p<0.001, ns., not significant. All experiments were performed independently at least three times. The data underlying this Figure can be found in S2 Data.

### S7 Fig.

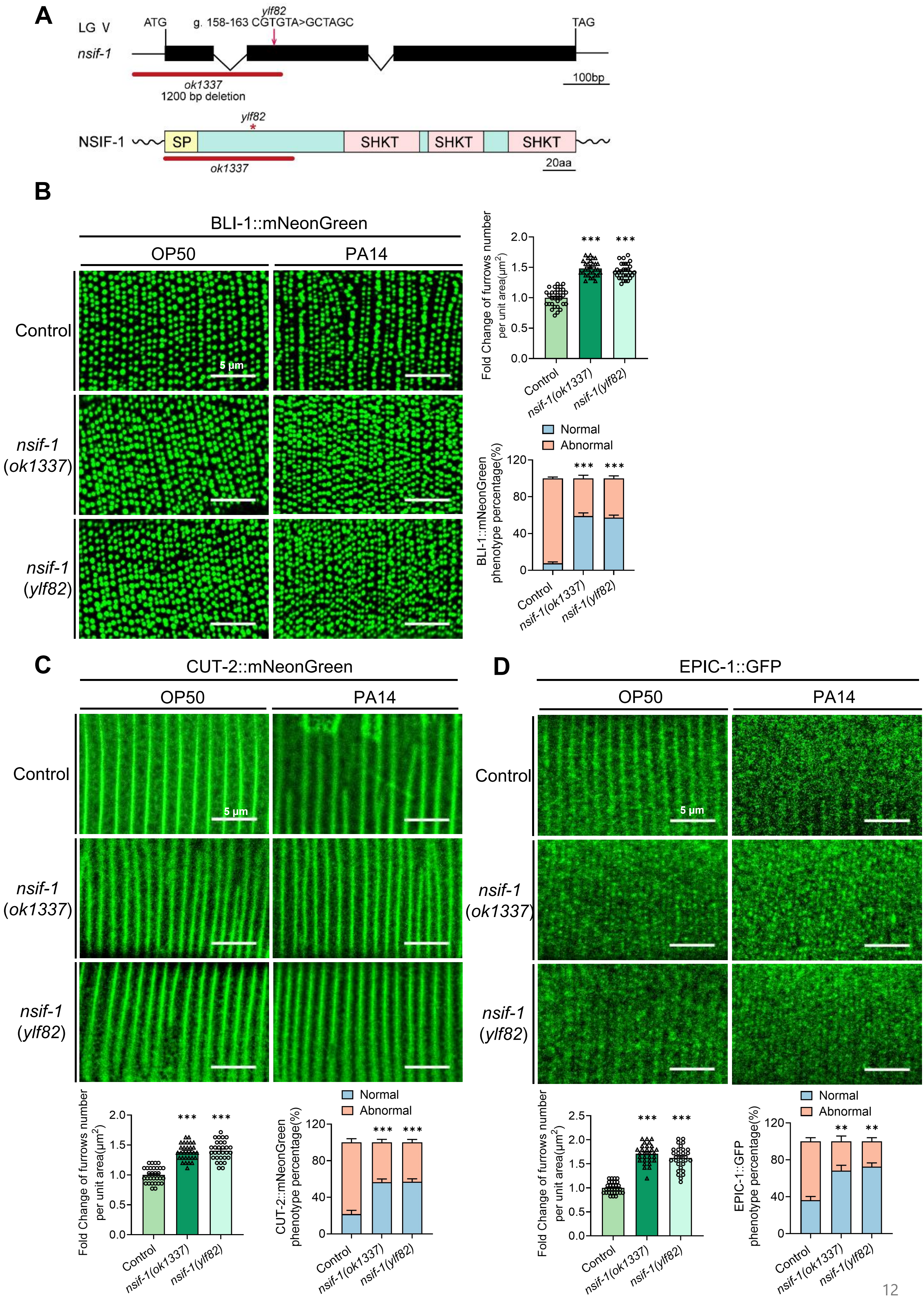
A
LG V
nsif-1
ATG
ylf82
g. 158-163 CGTGTA>GCTAGC
TAG
ok1337
1200 bp deletion
100bp
NSIF-1
SP
SHKT
SHKT
SHKT
20aa
B
BLI-1::mNeonGreen
OP50
PA14
Control
nsif-1 (ok1337)
nsif-1 (ylf82)
5 μm
Fold Change of furrows number per unit area(μm2)
Normal
Abnormal
BLI-1::mNeonGreen phenotype percentage(%)
C
CUT-2::mNeonGreen
CUT-2::mNeonGreen phenotype percentage(%)
D
EPIC-1::GFP
EPIC-1::GFP phenotype percentage(%)

**S7 Fig. NSIF-1 disrupts collagen integrity.**

**(A)** Schematic of the *nsif-1* gene and NSIF-1 protein. Exons are shown as filled boxes and introns as connecting lines. The positions of the *ok1337* deletion and the *ylf82* point mutation are indicated. The signal peptide (SP) and SHKT domains of NSIF-1 are shown.

**(B-D)** Confocal fluorescence images and quantitative analysis of BLI-1::mNeonGreen (B), CUT-2::mNeonGreen (C) and EPIC-1::GFP (D) in wild-type (WT), *nsif-1(ok1337)* and *nsif-1(ylf82)* following 24-hours PA14 exposure (infection initiated at L4 stage) or OP50. Quantification of cuticle furrows density were performed on 30 worms (n=30). And the quantitative analysis of abnormal pattern after 24h PA14 infection were performed on 100 worms with 3 times repeats (n=300 animals). Scale bar:5μm.

For all quantification, data are presented as mean ± SEM. Statistical comparisons were performed using one-way ANOVA followed by Tukey's multiple comparisons test. $^{**}p < 0.01$, $^{***}p < 0.001$. All experiments were performed independently at least three times. The data underlying this figure can be found in S2 Data.

### S8 Fig.

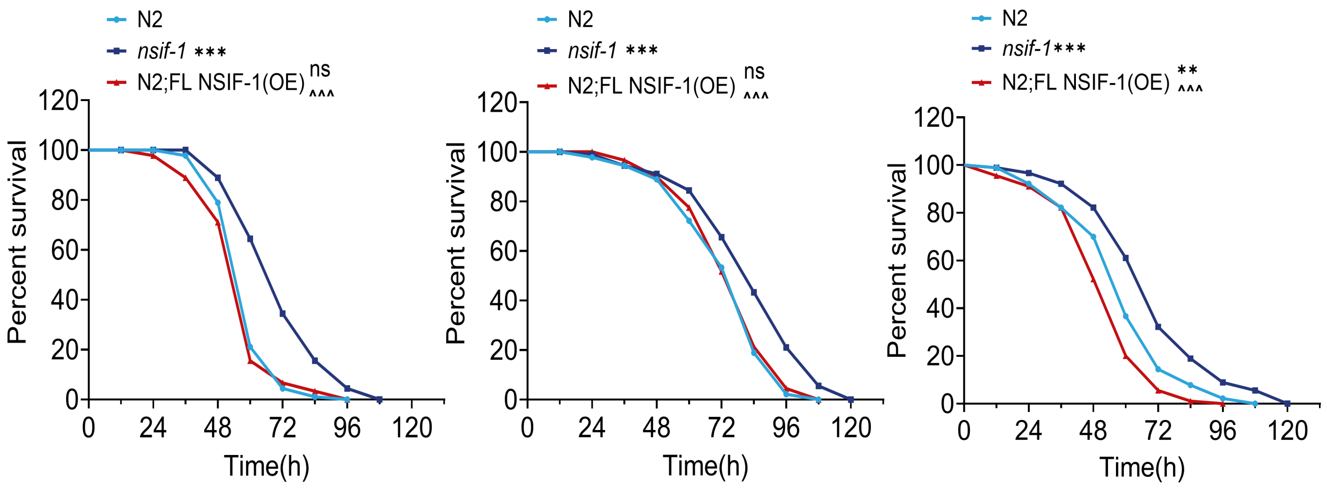

**S8 Fig. NSIF-1 enhances susceptibility to PA14**

Survival analysis of wild-type N2, *nsif-1(ok1337)* mutants, and N2 animals overexpressing full-length NSIF-1 driven by its own native promoter (P*nsif-1*) in the PA14 slow-killing assay. Data are representative of three independent biological replicates.

For all quantification, data are presented as mean ± SEM. Statistical comparisons were performed using Log rank (Mantel-Cox) test.* Indicates comparison with the wild-type, ^ indicates comparison with the *nsif-1(ok1337)* mutant. */^p<0.05, ***/^^^p<0.001; ns., not significant. All experiments were performed independently at least three times. The data underlying this figure can be found in S2 Data.

### S9 Fig.

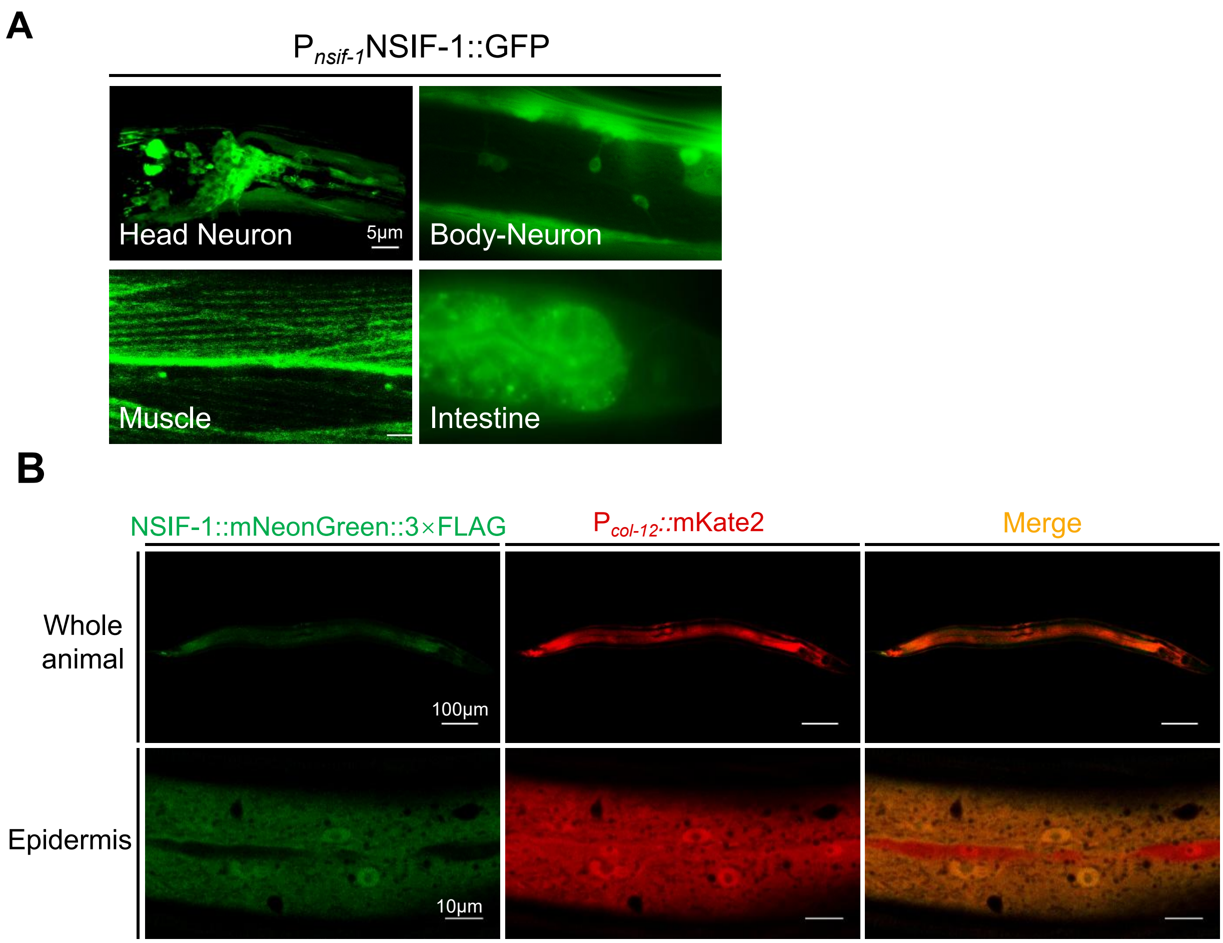

**S9 Fig. NSIF-1 expression pattern.**
**(A)** Confocal fluorescence imaging of transgenic animals expressing NSIF-1::GFP under the native *nsif-1* promoter ($P_{nsif\text{-}1}$NSIF-1::GFP). GFP expression was observed in various neurons, muscle cells, and the intestine. Scale bars: 5µm.
**(B)** Confocal fluorescence imaging of the CRISPR knock-in strain NSIF-1::mNeonGreen::3×FLAG (*syb9508*), showing that the tagged protein colocalizes with the epidermis, which is marked by $P_{col\text{-}12}$::mKate2. Scale bars: 100 µm (in whole animals) and 10 µm (in epidermis).

### S10 Fig.

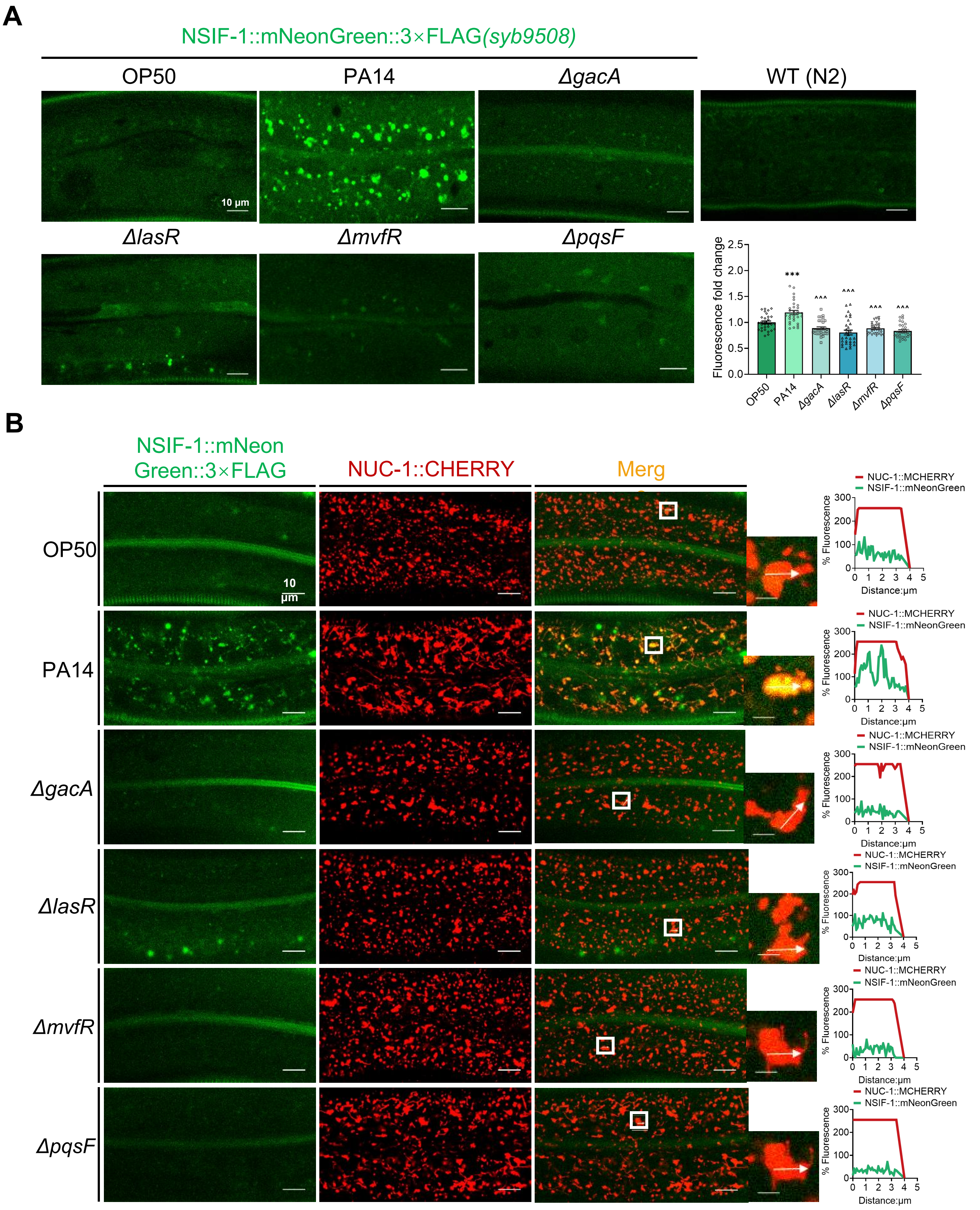
A
NSIF-1::mNeonGreen::3×FLAG(syb9508)
OP50
PA14
ΔgacA
WT (N2)
10 μm
ΔlasR
ΔmvfR
ΔpqsF
Fluorescence fold change
2.5
2.0
1.5
1.0
0.5
0.0
***
^^^
OP50
PA14
ΔgacA
ΔlasR
ΔmvfR
ΔpqsF
B
NSIF-1::mNeon Green::3×FLAG
NUC-1::CHERRY
Merg
OP50
PA14
ΔgacA
ΔlasR
ΔmvfR
ΔpqsF
10 μm
NUC-1::MCHERRY
NSIF-1::mNeonGreen
% Fluorescence
300
200
100
0
0 1 2 3 4 5
Distance:μm

**S10 Fig. PA14-induced upregulation and lysosomal localization of NSIF-1 are specific responses to active virulence.**

**(A)** Confocal fluorescence images and quantification of NSIF-1::mNeonGreen::3×FLAG(*syb9508*) knock-in worms following 24-h exposure to wild-type PA14 or the virulence regulator-deficient mutants *ΔgacA, ΔlasR, ΔmvfR,* and *ΔpqsF*. n=30 animals. Scale bars: 10 μm.

**(B)** Confocal fluorescence imaging of NSIF-1::mNeonGreen::3×FLAG and NUC-1::CHERRY following 24-h exposure to wild-type PA14 or the PA14 virulence regulator-deficient mutants *ΔgacA, ΔlasR, ΔmvfR,* and *ΔpqsF*. Scale bars: 10 μm.

For all quantification, data are presented as mean ± SEM. Statistical comparisons were performed using one-way ANOVA followed by Tukey's multiple comparisons test. *Indicates comparison with the OP50, ^indicates comparison with the PA14. ***/^^^p<0.001. All experiments were performed independently at least three times. The data underlying this figure can be found in S2 Data.

**S11 Fig.**

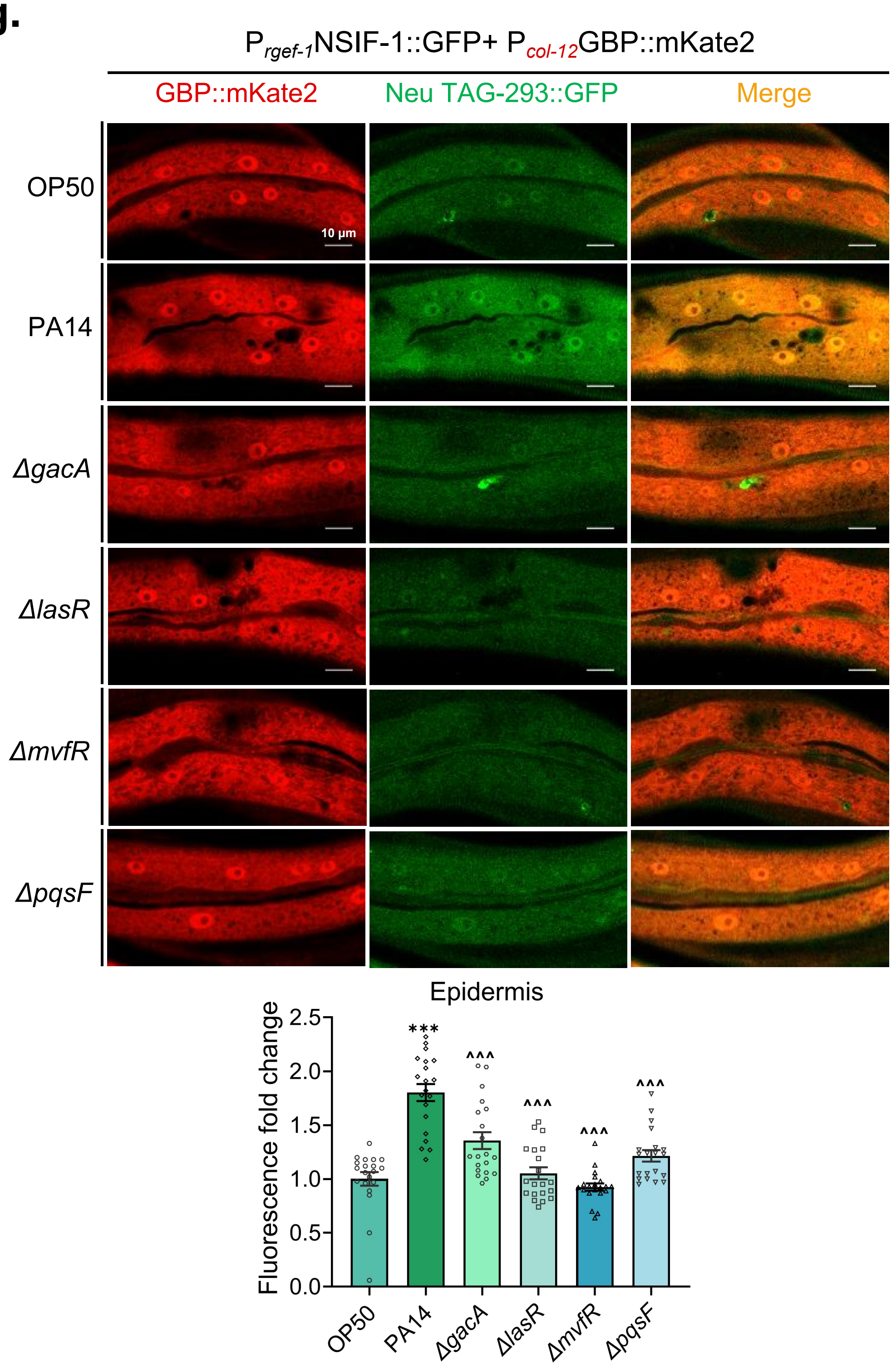

**S11 Fig. PA14 virulence regulators are required for neuronal NSIF-1 secretion to the epidermis.**

Confocal fluorescence images and quantification of neuronally secreted NSIF-1::GFP detected in the epidermis of transgenic animals expressing P$_{rgef-1}$NSIF-1::GFP (neuronal) and P$_{col-12}$GBP::mKate2 (epidermal) following 24 h of exposure to wild-type PA14 or the virulence-regulator-deficient mutants *ΔgacA, ΔlasR, ΔmvfR,* and *ΔpqsF*. n = 20 animals per condition. Scale bars: 10 µm.

For all quantification, data are presented as mean ± SEM. Statistical comparisons were performed using one-way ANOVA followed by Tukey's multiple comparisons test. *Indicates comparison with the OP50, ^indicates comparison with the PA14. ***/^^^p<0.001. All experiments were performed independently at least three times. The data underlying this figure can be found in S2 Data.

### S12 Fig.

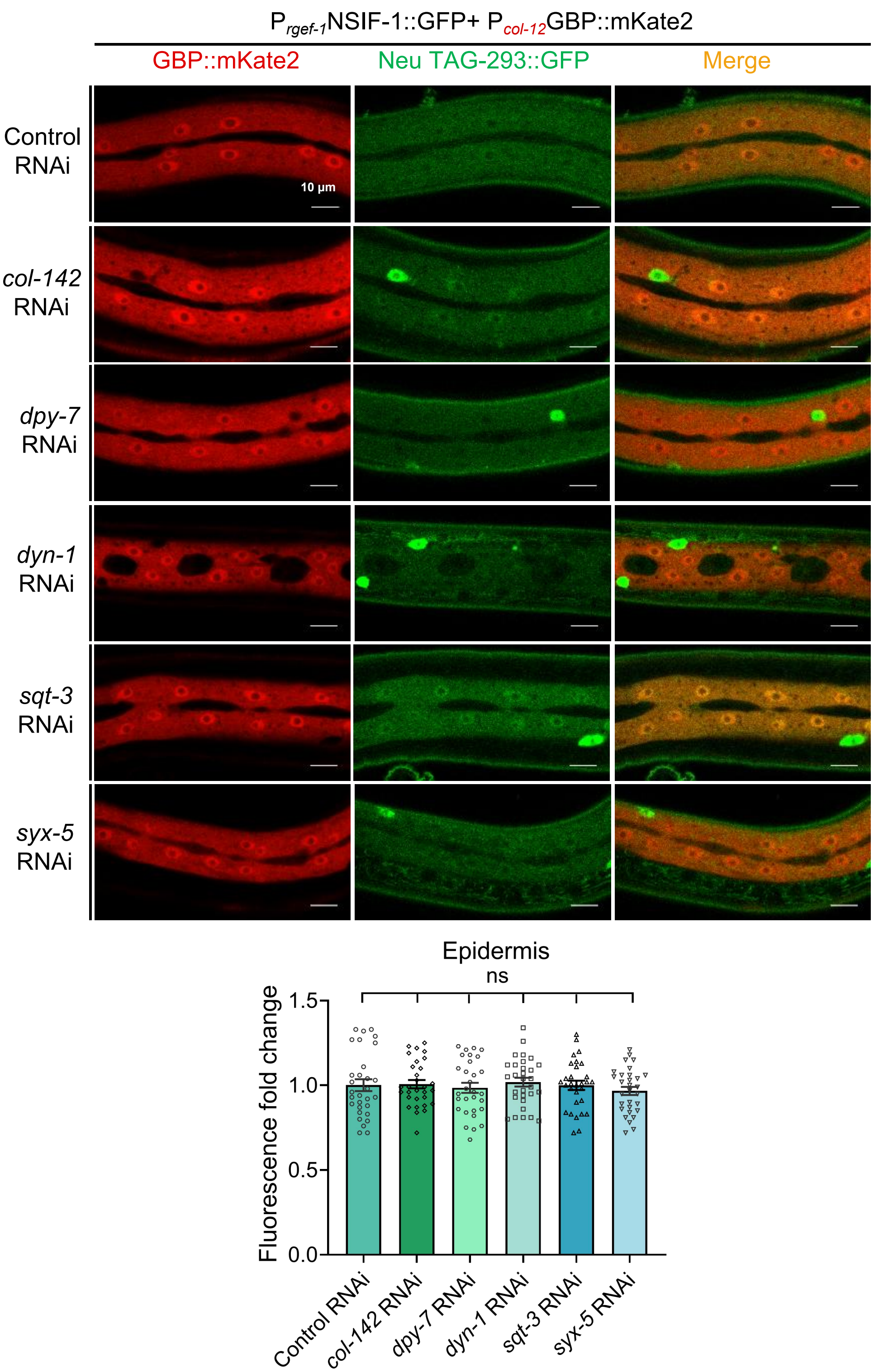

**S12 Fig. Collagen related genes disruption does not impair NSIF-1 secretion to the epidermis.**

Confocal fluorescence images and quantification of epidermal NSIF-1::GFP fluorescence in $P_{rgef\text{-}1}$NSIF-1::GFP;$P_{col\text{-}12}$GBP::mKate2 worms following *col-124, dpy-7, dyn-1, sqt-3,* or *syx-5* RNAi. No significant difference in epidermal NSIF-1::GFP fluorescence was observed compared with the control RNAi group. n=30 animals. Scale bars: 10 µm.

For all quantification, data are presented as mean ± SEM. Statistical comparisons were performed using one-way ANOVA followed by Tukey's multiple comparisons test. ns., not significant. All experiments were performed independently at least three times. The data underlying this figure can be found in S2 Data.

### S13 Fig.

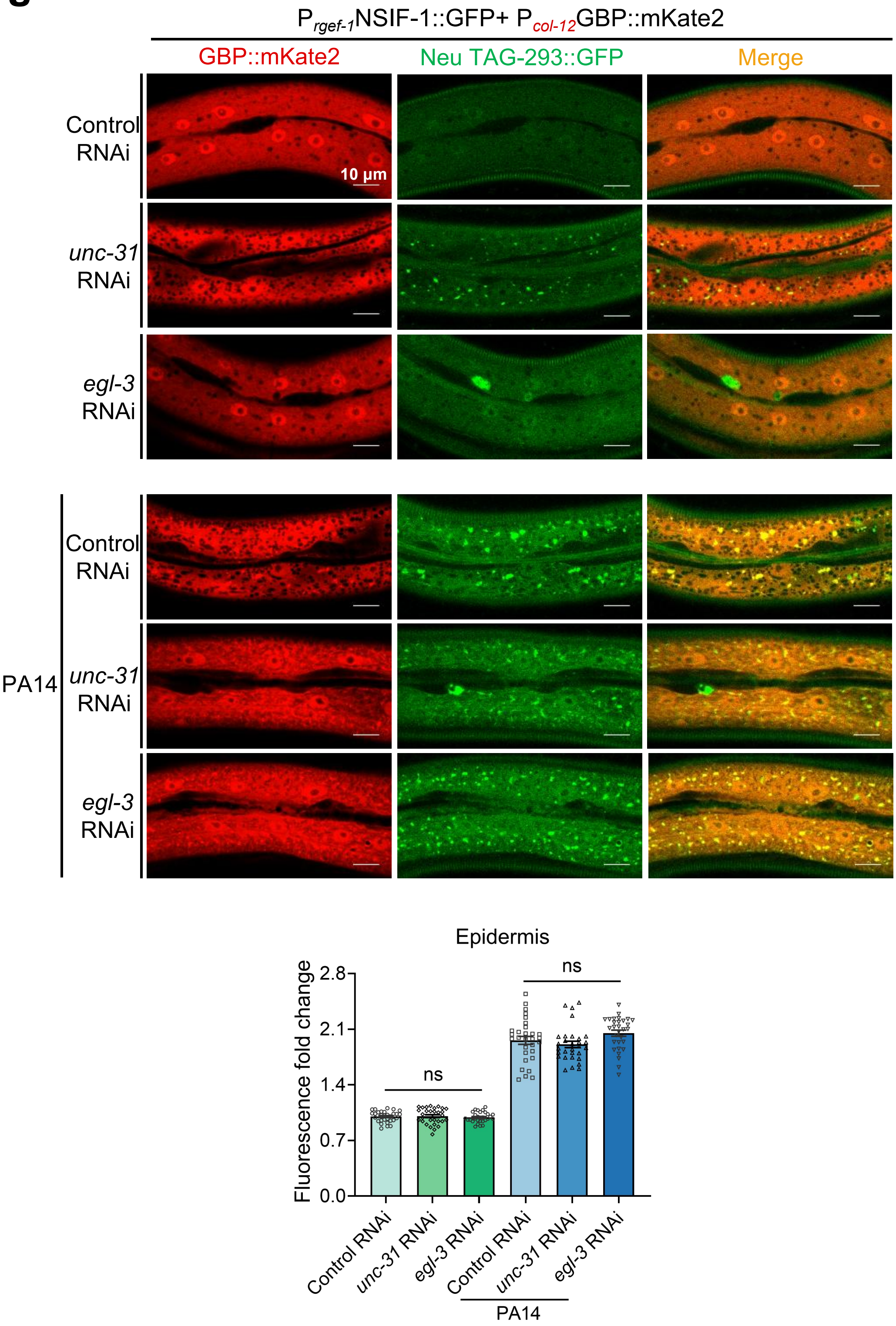

**S13 Fig. NSIF-1 release occurs through a non-canonical pathway independent of traditional neuropeptide processing mechanisms.**

Confocal fluorescence images and quantification of epidermal NSIF-1::GFP fluorescence in P*rgef-1*::NSIF-1::GFP;P*col-12*::GBP::mKate2 worms following 24-h exposure to OP50 or PA14 with *unc-31* or *egl-3* RNAi. Neither *unc-31* nor *egl-3* RNAi significantly affected epidermal NSIF-1::GFP fluorescence. n=30 animals. Scale bars: 10 µm.

For all quantification, data are presented as mean ± SEM. Statistical comparisons were performed using one-way ANOVA followed by Tukey's multiple comparisons test. ns., not significant. All experiments were performed independently at least three times. The data underlying this figure can be found in S2 Data.

### S14 Fig.

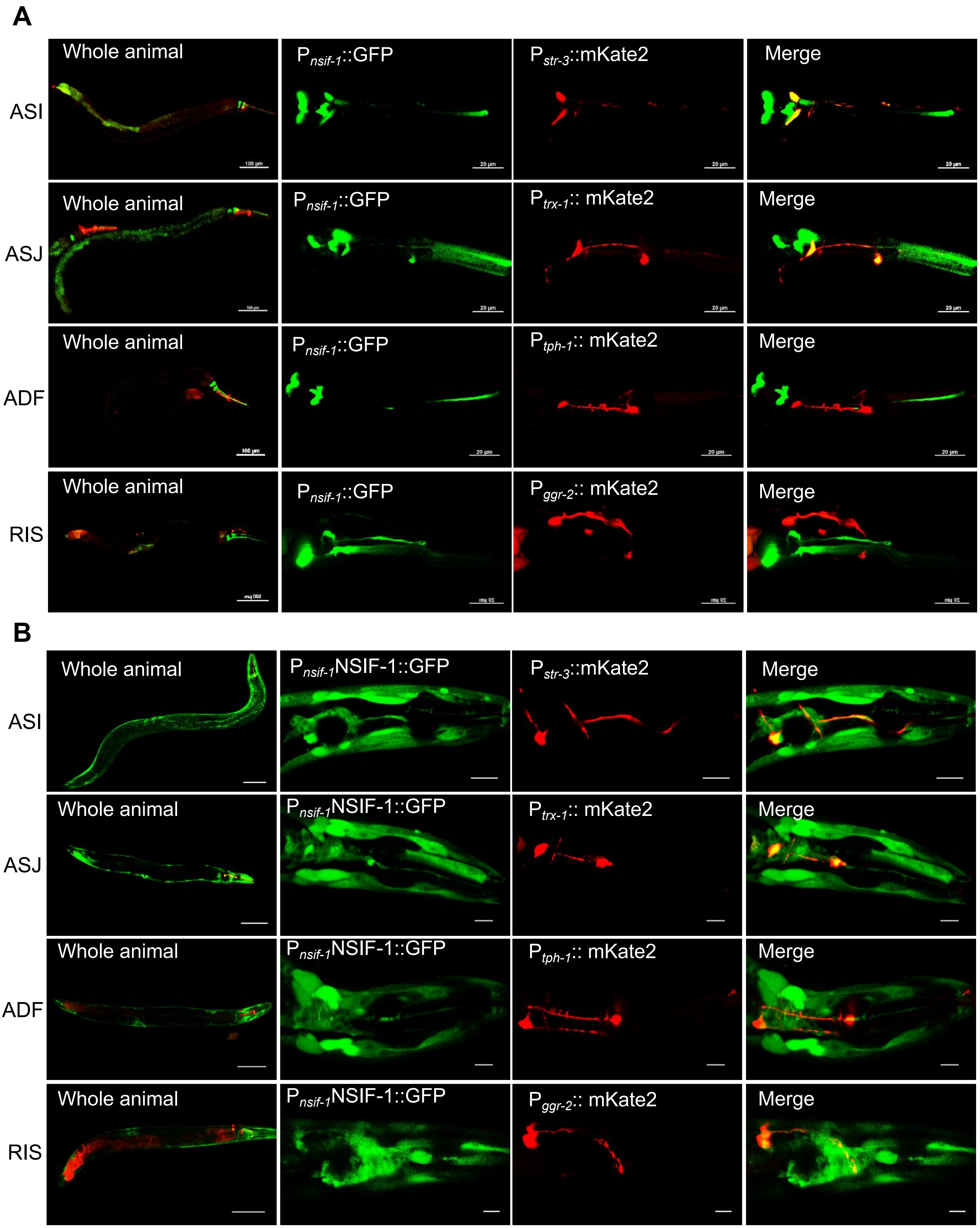

**S14 Fig. Co-localization analysis of NSIF-1 transcriptional and translational reporters with ASI, ASJ, ADF, and RIS neurons.**

**(A)** Localization of the NSIF-1 transcriptional reporter ($P_{nsif-1}$::GFP) in ASI ($P_{str-3}$::mKate2), ASJ ($P_{trx-1}$::mKate2), ADF ($P_{tph-1}$::mKate2), and RIS ($P_{ggr-2}$::mKate2) neurons. Scale bars: whole animal, 100 µm; others, 20 µm.

**(B)** Localization of the NSIF-1 transcriptional reporter ($P_{nsif-1}$NSIF-1::GFP) in ASI ($P_{str-3}$::mKate2), ASJ ($P_{trx-1}$::mKate2), ADF ($P_{tph-1}$::mKate2), and RIS ($P_{ggr-2}$::mKate2) neurons. Scale bars: whole animal, 100 µm; others, 10 µm.

### S15 Fig.

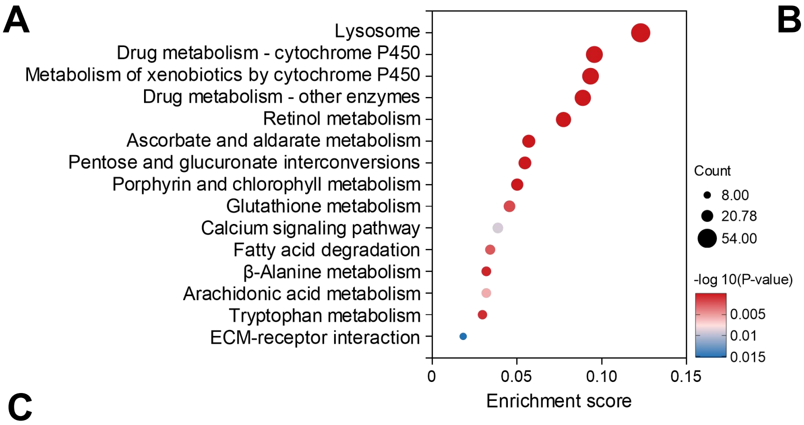

B

| Expression of Lysosome-related genes [*nsif-1(ok1337)* VS N2] | |
|---|---|
| Catalogs | Up regulated(54) |
| Membrane protein | 32 |
| v-ATPase | 0 |
| Protease | 12 |
| Glucocerebrosidase | 3 |
| Acid phosphatase | 1 |
| Non-protease hydrolases | 4 |
| Other hydrolase | 2 |

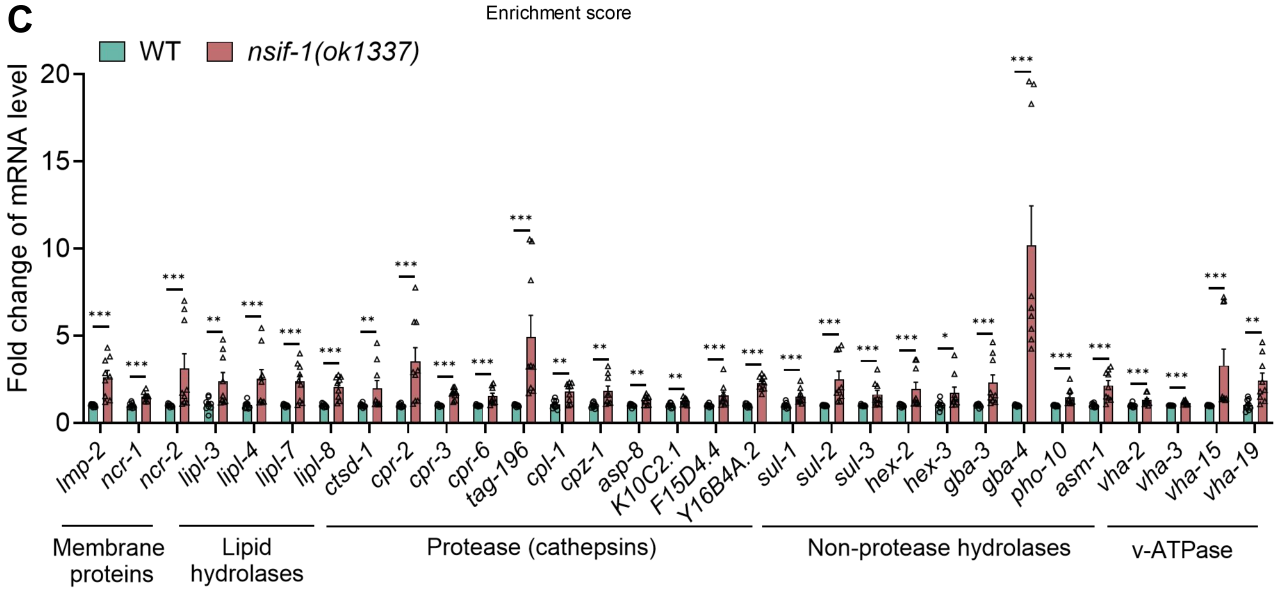

**S15 Fig. Loss of *nsif-1* upregulates lysosome-associated gene expression.**

**(A)** KEGG pathway analysis of differentially upregulated genes in *nsif-1(ok1337)* relative to wild-type (WT) from RNA-seq. Upregulated genes for KEGG pathway analysis were defined as those with a false discovery rate (FDR) < 0.05 and a fold change≥1.5.

**(B)** Functional categorization of 54 lysosomal genes upregulated in *nsif-1(ok1337)* (identified in A) from RNA-seq.

**(C)** qRT-PCR validation of lysosomal gene expression changes in WT and adult *nsif-1(ok1337)*, grouped by functional class: membrane proteins, lipid hydrolases, proteases, non-proteases, and v-ATPase components.

For all quantification, data are presented as mean±SEM. Statistical comparisons were performed using Mann–Whitney U-test. *p<0.05, **p<0.01, ***p<0.001; ns., not significant. All experiments were performed independently at least three times. The data underlying this figure can be found in S2 Data

### S16 Fig.

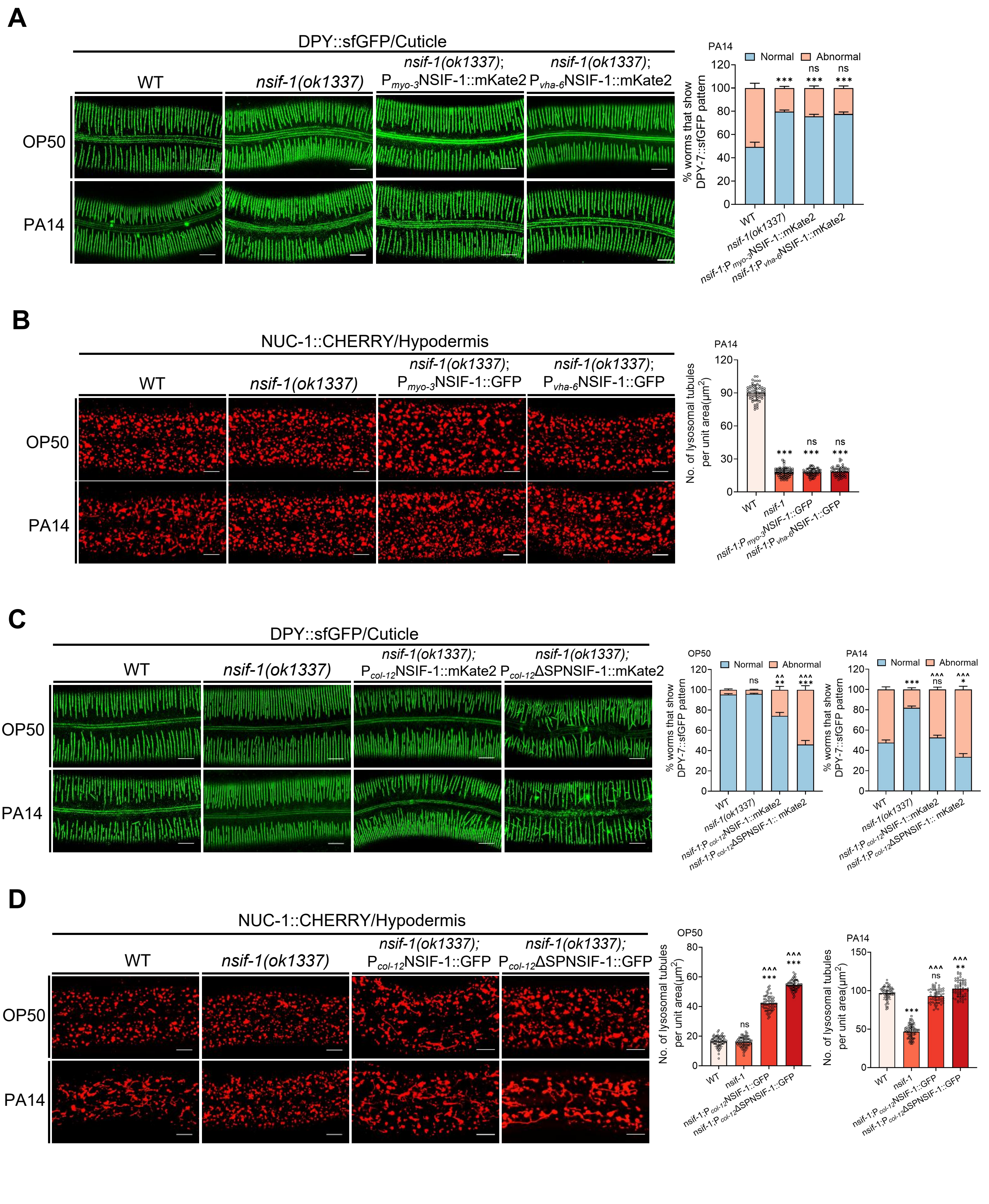
A
DPY::sfGFP/Cuticle
WT
nsif-1(ok1337)
nsif-1(ok1337); Pmyo-3NSIF-1::mKate2
nsif-1(ok1337); Pvha-6NSIF-1::mKate2
OP50
PA14
Normal
Abnormal
% worms that show DPY-7::sfGFP pattern
ns
***
nsif-1;P myo-3NSIF-1::mKate2
nsif-1;P vha-6NSIF-1::mKate2
B
NUC-1::CHERRY/Hypodermis
nsif-1(ok1337); Pmyo-3NSIF-1::GFP
nsif-1(ok1337); Pvha-6NSIF-1::GFP
No. of lysosomal tubules per unit area(μm2)
nsif-1
nsif-1;P myo-3NSIF-1::GFP
nsif-1;P vha-6NSIF-1::GFP
C
nsif-1(ok1337); Pcol-12NSIF-1::mKate2
nsif-1(ok1337); Pcol-12ΔSPNSIF-1::mKate2
nsif-1;P col-12NSIF-1::mKate2
nsif-1;P col-12ΔSPNSIF-1:: mKate2
^^
**
^^^
*
D
nsif-1(ok1337); Pcol-12NSIF-1::GFP
nsif-1(ok1337); Pcol-12ΔSPNSIF-1::GFP
nsif-1;P col-12NSIF-1::GFP
nsif-1;P col-12ΔSPNSIF-1::GFP

**S16 Fig. Functional analysis of multi-tissue rescue of NSIF-1 in regulating lysosomal homeostasis and cuticle integrity.**

**(A)** Confocal fluorescence imaging and quantitative analysis of DPY-7::sfGFP marked cuticle in wild-type(WT), *nsfi-1(ok1337)*, *nsfi-1(ok1337)*;$P_{myo-3}$NSIF-1::mKate2, *nsfi-1(ok1337)*;$P_{vha-6}$NSIF-1::mKate2 animals following 24-h PA14 exposure (infection initiated at the L4 stage). Quantification of abnormal DPY-7::sfGFP pattern performed on 300 worms (n=300 animals). Scale bars: 10 μm.

**(B)** Confocal fluorescence imaging and quantitative analysis of NUC-1::CHERRY marked lysosome in wild-type(WT), *nsfi-1(ok1337)*, *nsfi-1(ok1337)*; $P_{myo-3}$NSIF-1::mKate2, *nsfi-1(ok1337)*;$P_{vha-6}$NSIF-1::mKate2 animals following 24-h PA14 exposure (infection initiated at the L4 stage). Quantification of tubular lysosomes were quantified by counting three 35 × 25 μm² regions per worm (n=20 worms) . Scale bars: 10 μm.

**(C)** Confocal fluorescence imaging and quantitative analysis of DPY-7::sfGFP marked cuticle in wild-type(WT), *nsfi-1(ok1337)*, *nsfi-1(ok1337)*;$P_{col-12}$NSIF-1::mKate2, *nsfi-1(ok1337)*;$P_{col-12}$ΔSPNSIF-1::mKate2 animals following 24-h PA14 exposure (infection initiated at the L4 stage). Quantification of abnormal DPY-7::sfGFP pattern performed on 300 worms (n=300 animals). Scale bars: 10 μm.

**(D)** Confocal fluorescence imaging and quantitative analysis of NUC-1::CHERRY marked lysosome in wild-type(WT), *nsfi-1(ok1337)*, *nsfi-1(ok1337)*;$P_{col-12}$NSIF-1::mKate2, *nsfi-1(ok1337)*;$P_{col-12}$ΔSPNSIF-1::mKate2 animals following 24-h PA14 exposure (infection initiated at the L4 stage). Quantification of tubular lysosomes were quantified by counting three 35 × 25 μm² regions per worm (n = 20 worms). Scale bars: 10 μm.

For all quantification, data are presented as mean ± SEM. Statistical comparisons were performed using one-way ANOVA followed by Tukey's multiple comparisons test. *Indicates comparison with the wild-type, ^ indicates comparison with the *nsif-1(ok1337)* mutant; **/^^ $p < 0.01$, ***/^^^ $p < 0.001$; ns., not significant. All experiments were performed independently at least three times. The data underlying this figure can be found in S2 Data.

### S17 Fig.

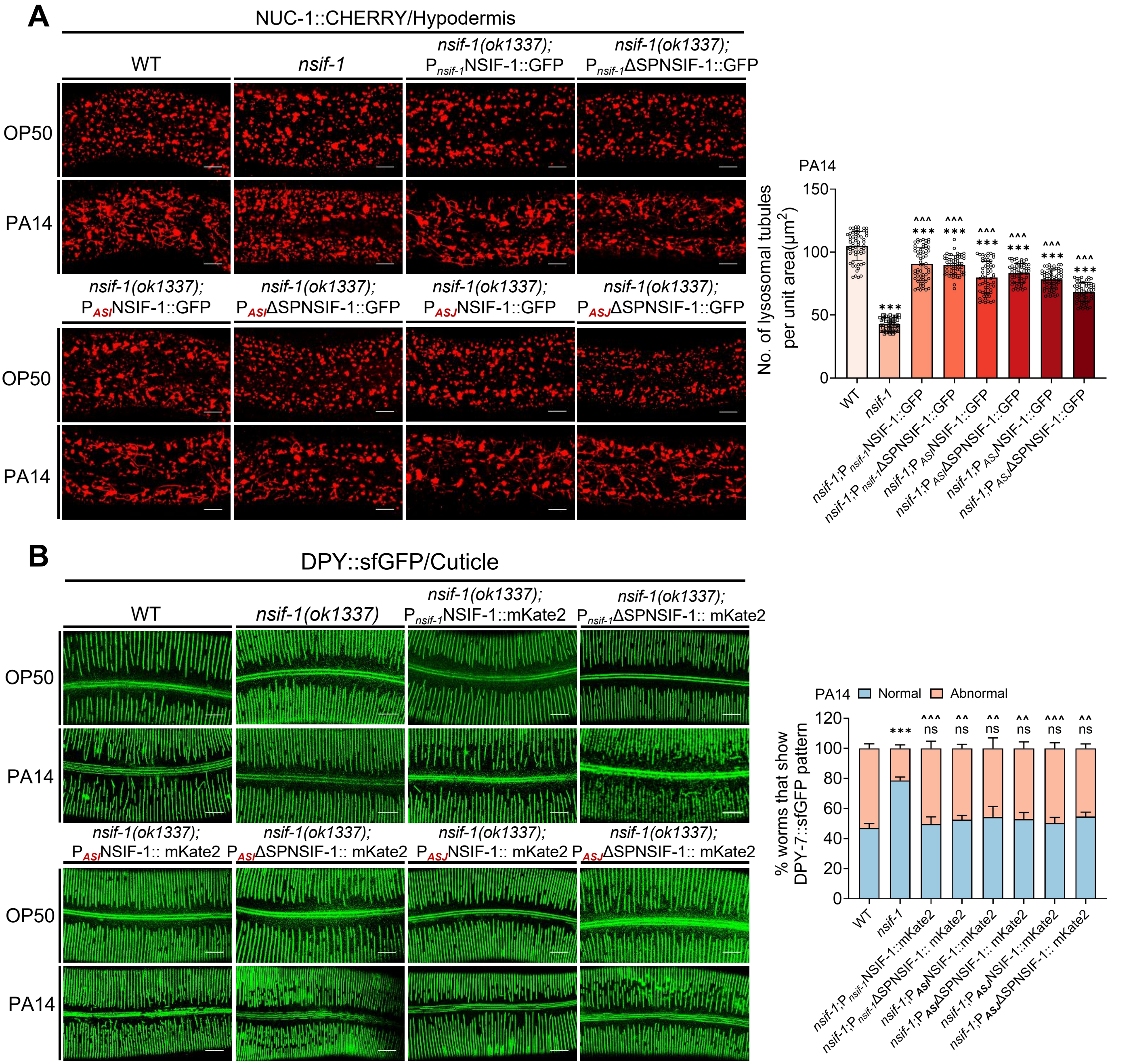

A
NUC-1::CHERRY/Hypodermis
WT
nsif-1
nsif-1(ok1337); PnsifNSIF-1::GFP
nsif-1(ok1337); PnsifΔSPNSIF-1::GFP
OP50
PA14
nsif-1(ok1337); PASINSIF-1::GFP
nsif-1(ok1337); PASIΔSPNSIF-1::GFP
nsif-1(ok1337); PASJNSIF-1::GFP
nsif-1(ok1337); PASJΔSPNSIF-1::GFP
PA14
No. of lysosomal tubules per unit area(μm2)
B
DPY::sfGFP/Cuticle
WT
nsif-1(ok1337)
nsif-1(ok1337); Pnsif-1NSIF-1::mKate2
nsif-1(ok1337); Pnsif-1ΔSPNSIF-1:: mKate2
nsif-1(ok1337); PASINSIF-1:: mKate2
nsif-1(ok1337); PASIΔSPNSIF-1:: mKate2
nsif-1(ok1337); PASJNSIF-1:: mKate2
nsif-1(ok1337); PASJΔSPNSIF-1:: mKate2
PA14 Normal Abnormal
% worms that show DPY-7::sfGFP pattern

### S18 Fig.

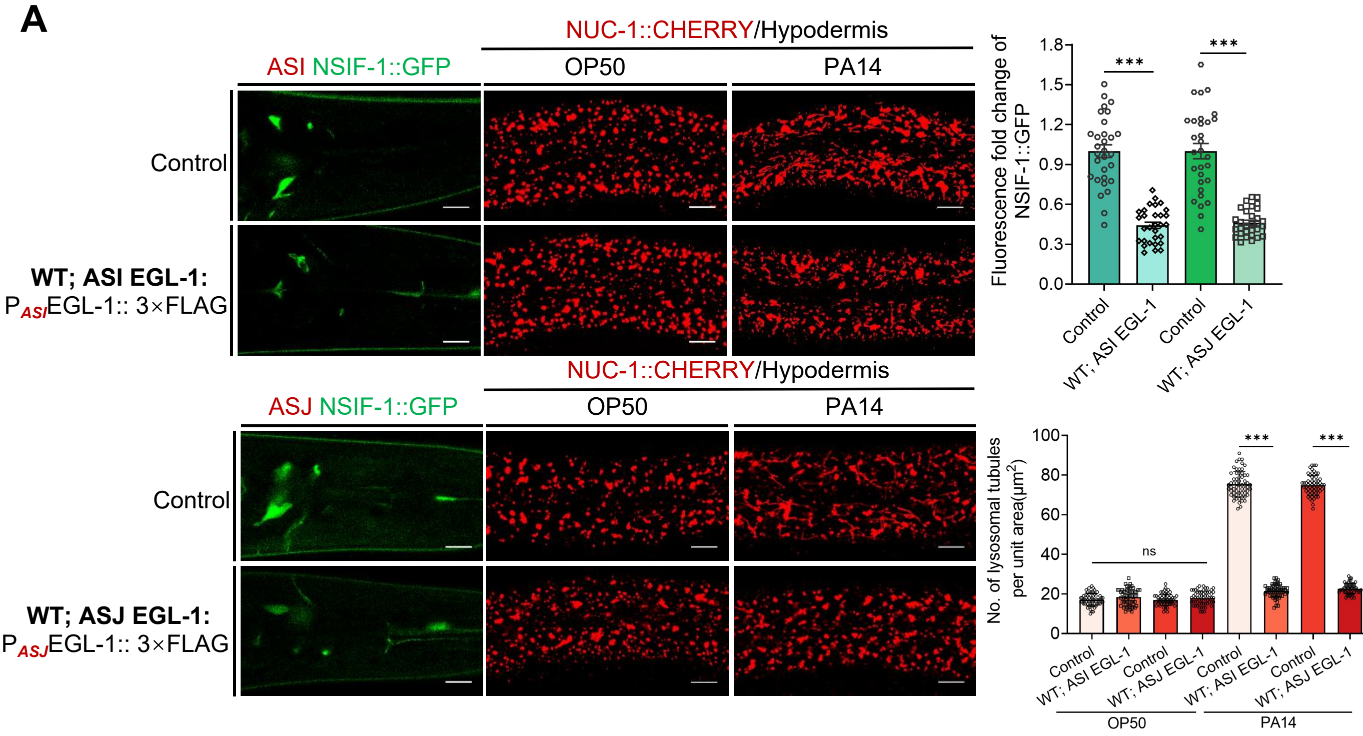

A
NUC-1::CHERRY/Hypodermis
ASI NSIF-1::GFP
OP50
PA14
Control
WT; ASI EGL-1:
$P_{ASI}$EGL-1:: 3×FLAG
ASJ NSIF-1::GFP
WT; ASJ EGL-1:
$P_{ASJ}$EGL-1:: 3×FLAG
Fluorescence fold change of NSIF-1::GFP
1.8
1.5
1.2
0.9
0.6
0.3
0.0
***
Control
WT; ASI EGL-1
WT; ASJ EGL-1
No. of lysosomal tubules per unit area(μm²)
100
80
60
40
20
0
ns

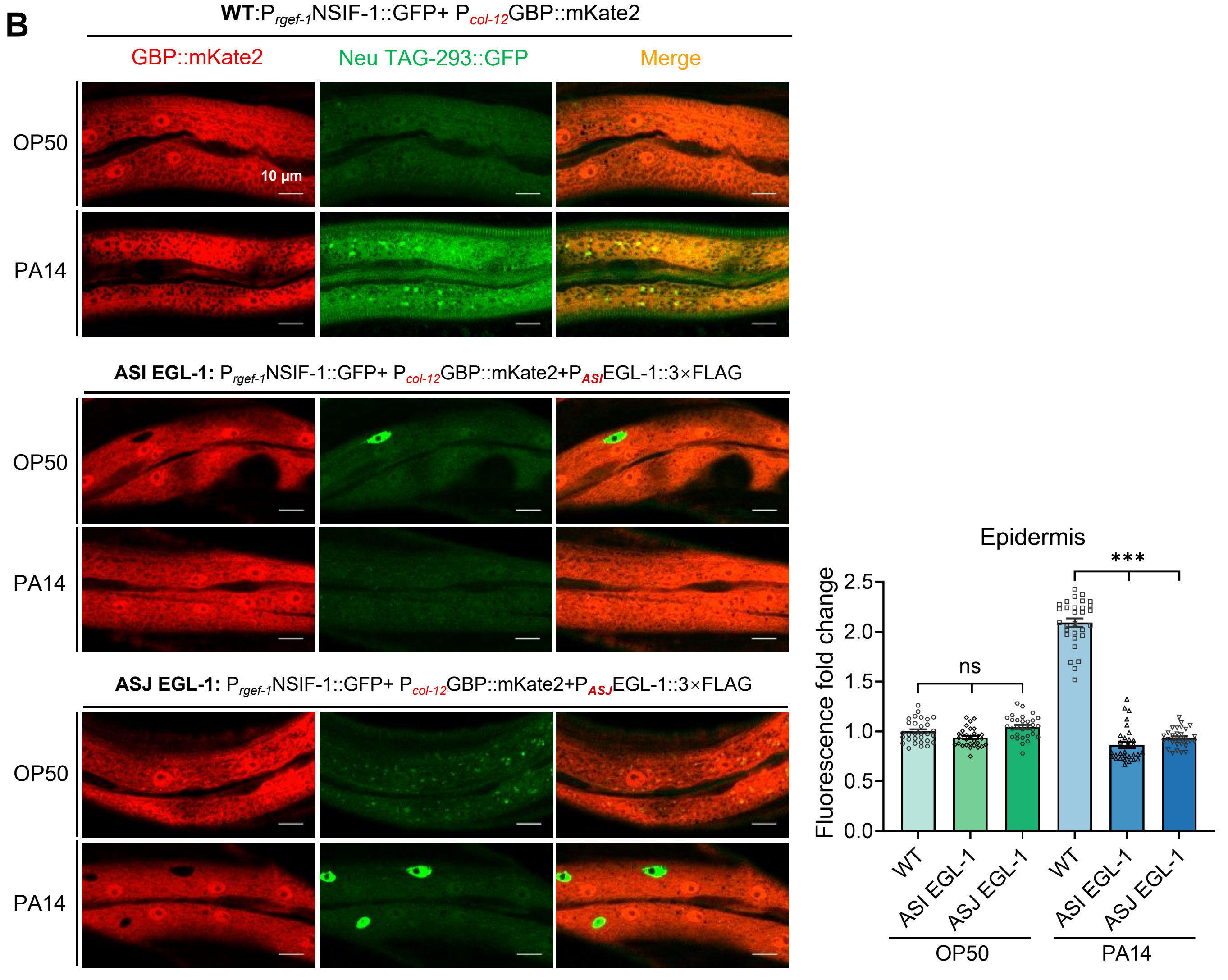

B
WT:$P_{rgef-1}$NSIF-1::GFP+ $P_{col-12}$GBP::mKate2
GBP::mKate2
Neu TAG-293::GFP
Merge
OP50
PA14
10 μm
ASI EGL-1: $P_{rgef-1}$NSIF-1::GFP+ $P_{col-12}$GBP::mKate2+$P_{ASI}$EGL-1::3×FLAG
ASJ EGL-1: $P_{rgef-1}$NSIF-1::GFP+ $P_{col-12}$GBP::mKate2+$P_{ASJ}$EGL-1::3×FLAG
Epidermis
Fluorescence fold change
2.5
2.0
1.5
1.0
0.5
0.0
ns
***
WT
ASI EGL-1
ASJ EGL-1

**S18 Fig. ASI/ASJ neuron ablation reduces NSIF-1 secretion to the epidermis and preserves lysosomal homeostasis during PA14 infection.**
**(A)** Confocal microscopy images and quantification of ASI/ASJ neuron ablation efficiency and epidermal lysosomal morphology. EGL-1-mediated genetic ablation of ASI and ASJ neurons resulted in a significant reduction of NSIF-1::GFP fluorescence in these neurons. Following 24 h of PA14 exposure (initiated at the L4 larval stage), ablation of ASI/ASJ neurons prevented the PA14-induced reduction in tubular lysosomes. NSIF-1::GFP fluorescence intensity was quantified in 30 worms. Tubular lysosomes were quantified by counting three 35 × 25 $\mu m^2$ regions per worm (n=20 worms).
**(B)** Confocal microscopy images and quantification of epidermally captured NSIF-1::GFP in WT, ASI-ablated, or ASJ-ablated worms following 24-h PA14 exposure in transgenic animals expressing $P_{rgef\text{-}1}$NSIF-1::GFP+$P_{col\text{-}12}$GBP::mKate2 (infection initiated at the L4 stage) Compared with control worms, genetic ablation of either ASI or ASJ neurons significantly reduced the epidermal NSIF-1::GFP signal . n=30 animals. Scale bars: 10 µm.
For all quantification, data are presented as mean ± SEM. Statistical comparisons were performed using Mann–Whitney U-test(A) and one-way ANOVA followed by Tukey's multiple comparisons test(B). ***$p<0.001$, ns., not significant. All experiments were performed independently at least three times. The data underlying this figure can be found in S2 Data.

### S19 Fig.

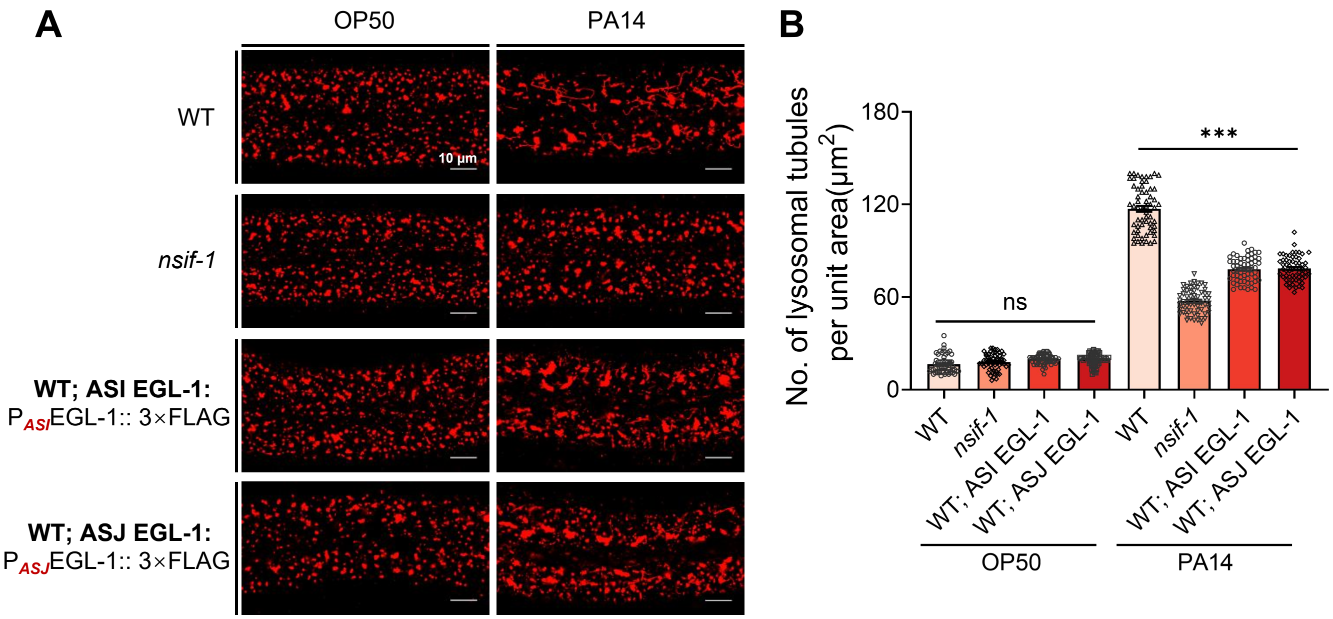

**S19 Fig. ASI/ASJ neuron ablation enhances epidermal lysosomal function.**

**(A)** Confocal fluorescence imaging of NUC-1::CHERRY in wild-type (WT), *nsfi-1(ok1337)*, and animals with ASI and ASJ neurons ablated, following 24 h of PA14 exposure (initiated at the L4 larval stage). Scale bars: 10 µm.

**(B)** Quantitative analysis of the number of tubular lysosomes in wild-type (WT), n*sfi-1(ok1337)*, and ASI/ASJ-ablated animals. The number of tubular lysosomes was quantified by counting within three 35×25(µm$^2$) unit areas per worm (n=20 animals).

For all quantification, data are presented as mean ± SEM. Statistical comparisons were performed using one-way ANOVA followed by Tukey's multiple comparisons test. *p<0.05, **p<0.01, ***p<0.001, ns., not significant. All experiments were performed independently at least three times. The data underlying this figure can be found in S2 Data.

### S20 Fig.

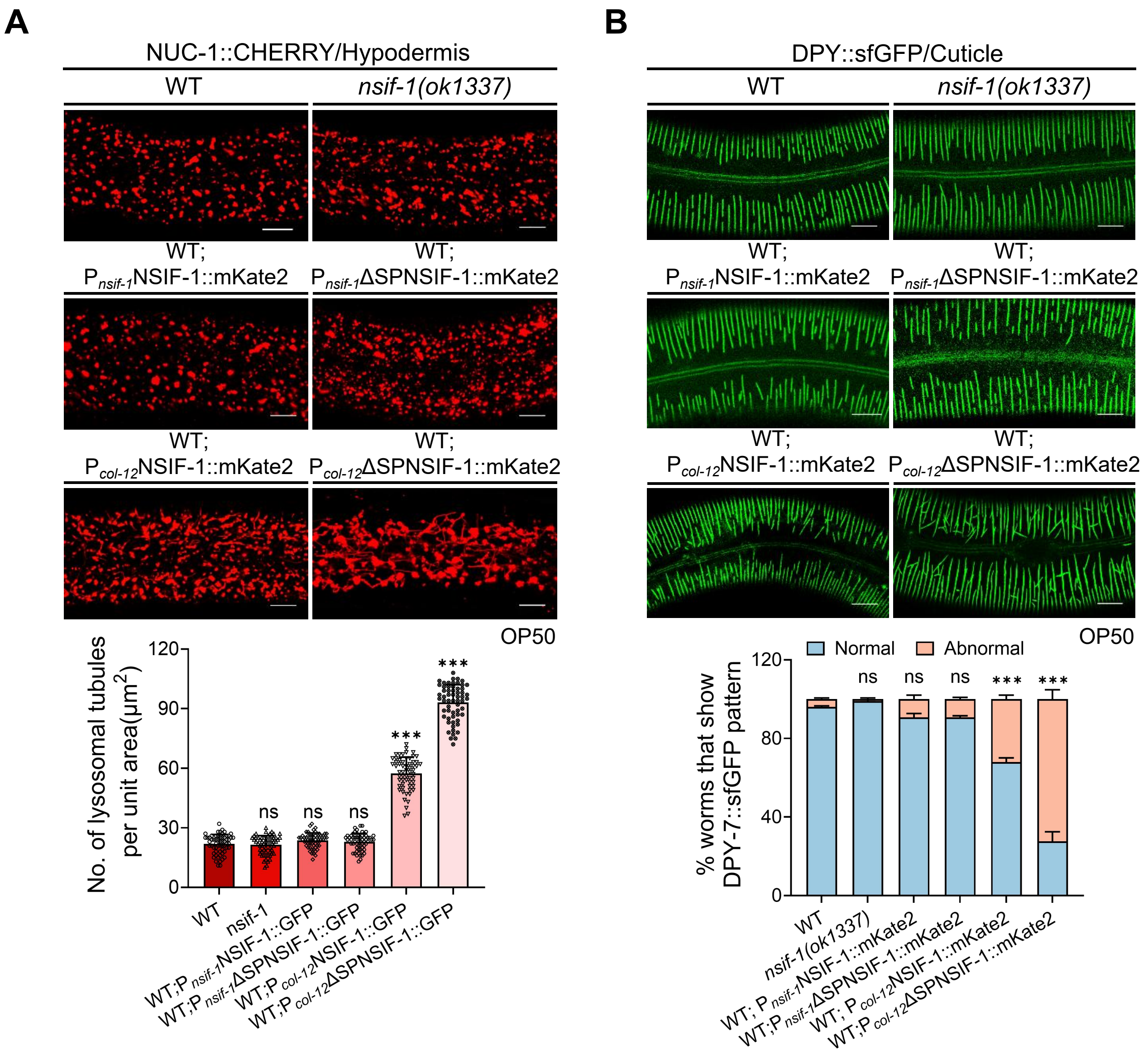

**S20 Fig. Epidermal expression of NSIF-1 is sufficient to impact lysosomal homeostasis and cuticle integrity.**

**(A)** Confocal fluorescence imaging and quantification of NUC-1::CHERRY-labeled lysosomes in wild-type (WT), *nsif-1(ok1337)* mutants, and WT animals expressing NSIF-1::mKate2 or the signal-peptide-deleted variant (ΔSP-NSIF-1::mKate2) under the control of the endogenous *nsif-1* promoter (P*nsif-1*) or the epidermal promoter (P*col-12*). Quantification was performed on 20 animals (n=20). Scale bars: 10 μm.

**(B)** Confocal fluorescence imaging and quantification of DPY-7::sfGFP-marked cuticle in WT, *nsif-1(ok1337)* mutants, and WT animals expressing NSIF-1::mKate2 or the signal-peptide-deleted variant (ΔSP-NSIF-1::mKate2) under the control of the endogenous *nsif-1* promoter (P*nsif-1*) or the epidermal promoter (P*col-12*). Quantification was performed on 300 animals (n=300). Scale bars: 10 μm.

For all quantification, data are presented as mean ± SEM. Statistical comparisons were performed using one-way ANOVA followed by Tukey's multiple comparisons test. ***p<0.001, ns., not significant. All experiments were performed independently at least three times. The data underlying this figure can be found in S2 Data.

### S21 Fig.

#### cNLS Mapper Result

| Predicted NLSs in query sequence |
|---|
| MFGLLVSCILAFTVPESAFADISGDLNCTQYNGTFFVWTPAAVACSNAVS 50 |
| DASCTALYPTEDEQGYPAAGNNAGRPLACFTTAAETPAPVDGDMKKAALT 100 |
| NCAKTCGFCCNTDDYSCPNAQFPRLNCDTITNNQCKDPNWRTIIATDCPS 150 |
| ACGFCNQGGCVDAVVDCANDRSICQSVGMQEFVNQNCQRTCGRCGSSTGN 200 |
| PSVPGGGSCTNYQADSSTACAAWAGNGFCTNTFYTEAQRKASCATTCRLC 250 |

| Predicted monopartite NLS | | |
|---|---|---|
| Pos. | Sequence | Score |

| Predicted bipartite NLS | | |
|---|---|---|
| Pos. | Sequence | Score |
| 91 | DGDMKKAALTNCAKTCGFCCNTDDYSCPNAQFP | 3.3 |

**S21 Fig. Identification of a putative NLS sequence in NSIF-1 using cNLS Mapper.**

The online tool (https://nls-mapper.iab.keio.ac.jp/) predicted a candidate sequence with a score of 3.3.

### S22 Fig.

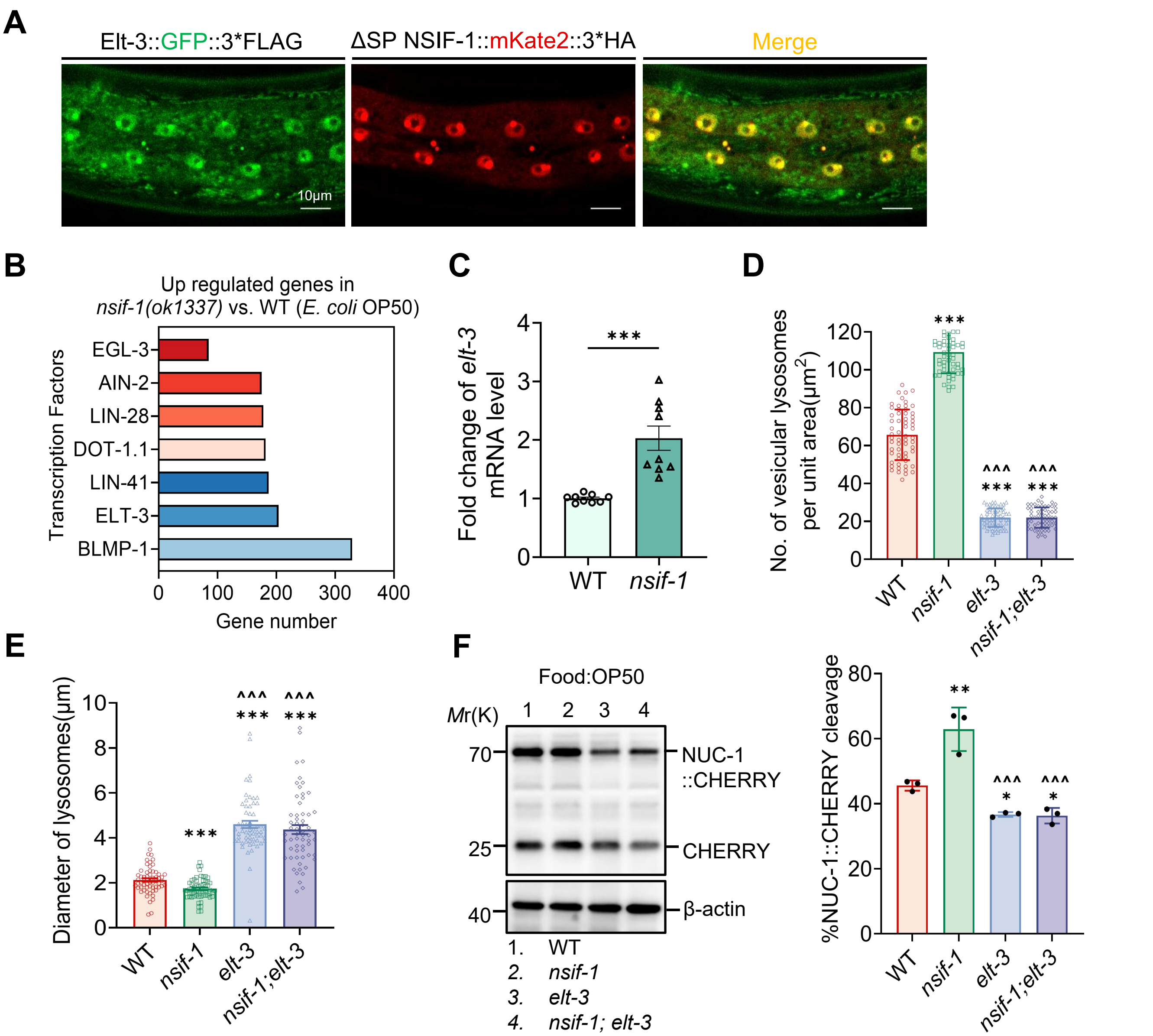

**S22 Fig. ELT-3 functions downstream of NSIF-1 to regulate epidermal lysosomal function.**

**(A)** Confocal fluorescence imaging of localization of ELT-3::GFP and $P_{col-12}$ΔSPNSIF-1::mKate2::3×HA in adult animals. Scale bars, 10μm.

**(B)** Transcription factor binding analysis identified ~200 ELT-3-regulated genes among transcripts upregulated in *nsif-1* mutants.

**(C)** qRT-PCR analysis of *elt-3* expression changes in adult (L4+24h) WT and *nsif-1(ok1337)* animals. Three biological replicates, each with three technical replicates, were analyzed for *elt-3*.

**(D-E)** Analysis of hypodermal lysosomes in WT, *nsif-1(ok1337), elt-3(vp1),* and *nsif-1(ok1337);elt-3(vp1)* animals expressing NUC-1::CHERRY following 24 h of PA14 exposure. The number of vesicular lysosomes (D) was quantified within three 35 × 25 μm² regions per worm (n = 20 animals). The diameter of vesicular lysosomes (E) was determined by measuring the six largest vesicles within a 35 × 25 μm² region per worm (n = 10 animals)

**(F)** Immunoblots and quantification showing cleavage of CHERRY from NUC-1::CHERRY expressed in the epidermis of adult wild type(WT), *nsif-1(ok1337), elt-3(gk121), nsif-1(ok1337);elt-3(vp1)*. N=3 biological replicates.

For all quantification, data are presented as mean ± SEM. Statistical comparisons were performed using Mann–Whitney U-test (C) and one-way ANOVA followed by Tukey's multiple comparisons test (D-F). *Indicates comparison with the wild-type, ^ indicates comparison with the *nsif-1(ok1337)* mutant; */^p< 0.05, **/^^p< 0.01, ***/^^^p< 0.001; ns., not significant. All experiments were performed independently at least three times. The data underlying this figure can be found in S2 Data.
